# Effect of Genotype and Age on a Defined Microbiota in Gnotobiotic SCID Piglets

**DOI:** 10.1101/2024.09.03.611011

**Authors:** Katherine M. Widmer, Faith Rahic-Seggerman, Ahlea Forster, Amanda Ahrens-Kress, Mary Sauer, Shankumar Mooyottu, Akhil Vinithakumari, Aaron Dunkerson-Kurzhumov, Brett Sponseller, Matti Kiupel, Stephan Schmitz-Esser, Christopher K. Tuggle

**Affiliations:** Department of Animal Science, Iowa State University, Ames, IA 50011, USA; Laboratory Animal Resources, Iowa State University, Ames, IA 50011, USA; Department of Veterinary Pathology, College of Veterinary Medicine, Iowa State University, Ames, IA 50011, USA; Department of Pathobiology, Auburn University, Auburn, AL, 36849, USA; Department of Veterinary Microbiology and Preventive Medicine, College of Veterinary Medicine, Iowa State University, Ames, IA 50011, USA; Department of Pathobiology and Diagnostic Investigation, College of Veterinary Medicine, Michigan State University, East Lansing, MI 48824, USA

## Abstract

Severe combined immunodeficient (SCID) individuals lack functional T and B lymphocytes, leading to a deficient adaptive immune system. SCID pigs are a unique large animal biomedical model as they possess many similarities to humans, allowing for the collection of translatable data in regenerative medicine, cancer, and other biomedical research topics. While many studies suggest early gut microbiota development is necessary for developing the intestinal barrier and immune system, these animals are often cesarian section derived, leaving them uncolonized for normal intestinal microflora. The hypothesis was that an increase in complexity of microbiota inoculum will allow for more stability in the composition of the gut microbiota of SCID piglets. This was tested across multiple litters of SCID piglets with three different defined microbiota consortium (2-strain, 6-strain, 7-strain). All piglets received their designated defined microbiota by oral gavage immediately after birth and again 24 hours later. There was no effect of SCID genotype on the composition of the gut microbiota, but there was a significant effect due to piglet age. Additionally, all three defined microbiota consortia were deemed safe to use in SCID piglets, and the 7-strain microbiota was the most stable over time. Based on these results, the 7-strain defined microbiota will be added to the SCID pig husbandry protocol, allowing for a more reproducible model.

## 1 Introduction

Severe combined immunodeficient (SCID) animal models have become important tools in biomedical research. The SCID genotype can be produced naturally due to mutations or by genetic modification of specific genes. These mutations lead to a SCID phenotype where T and B cells are deficient or non-functional (Cossu, 2010). Due to their compromised immune system, SCID animals allow for studies in cancer therapies, drug trials, and xenotransplants, making them a valuable component of pre-clinical research (Boettcher et al., 2018; Singer et al., 2019).

SCID mice are well established models that are cheaper and easier to care for than a large animal SCID model; however, research involving mice does not always translate efficiently to human medicine (Uhl & Warner, 2015). Boettcher and colleagues (2019) demonstrated that tumors grown in SCID mice were phenotypically different from the human tumor the cells were derived, while the tumors grown in the SCID pigs were much more similar. The pig genome is another area that is more similar to humans than the mouse genome (Junhee Seok et al., 2013; Wernersson et al., 2005). These similarities lead to the expectation that using a SCID pig model would allow for better data translation to human medicine. Additionally, in specific disciplines, such as tissue generation research on repair of cardiac disease, the FDA states that data obtained from small animal models is insufficient (FDA, 2009). This regulation demonstrates the benefit of a SCID pig model in biomedical research as opposed to SCID mice.

SCID pigs can occur naturally due to mutation in the Artemis gene, which is essential for DNA repair during the stages of development for T and B lymphocytes. When two mutations occur on different alleles of the Artemis gene (ART ^-/-^) in piglets, the resulting phenotype is a deficiency in T and B cells with functioning natural killer (NK) cells (Waide et al., 2015).

Genetic modification is another method to create SCID pigs. While there are many different methods to create genetically modified SCID pigs, this article highlights the use of a mutation in the IL-2 receptor subunit gamma chain (IL2RG ^-/Y^), which is necessary for lymphoid development. The resulting phenotype is a deficiency in T and NK cells with the presence of B cells that are not functional (Suzuki et al., 2012).

SCID pigs require a large, specific pathogen free (SPF) environment for long term survival (Lee et al., 2014), an example of which was previously described (Powell et al., 2018). Snatch farrowing was utilized to obtain the SCID piglets in aseptic manner prior to entry into the housing environment. Snatch farrowing involves catching piglets as they exit the vaginal canal preventing piglets from encountering contaminated surfaces (Huang et al., 2013). This method had some challenges, as the vaginal microbiota can be host to opportunistic pathogens, risking infection in the piglets. There was a change in protocol to avoid this risk of infection and it involves the use of cesarian section (Boettcher et al., 2020). The cesarian section method does not allow SCID piglets to receive microbiota from the sow like they would with a vaginal delivery (Wang et al., 2013), leading to an uncolonized gut where opportunistic pathogenic bacteria can thrive. The lack of colonization presents challenges, as it is known that early colonization of the gut is essential for piglet health (Kelly et al., 2007; Olszak et al., 2012). We tested three different defined microbiota that varied in complexity over multiple litters of SCID piglets and hypothesized that the most complex defined microbiota would be the most stable in the gut microbiome of the piglets.

With the knowledge of the significance of a colonized gut for piglet health, alternatives to snatch farrowing were sought, including providing the cesarian-derived SCID piglets with an artificial microbiome. We created three defined microbiota consortia, each with increased complexity. The selected bacteria in the defined microbiota were adapted from a defined commensal microbiota (DMF) designed to mimic the infant gut (Huang et al., 2018). This study aimed to determine the effect of genotype and piglet age on the defined microbiota in the gut of SCID piglets. Additionally, the goal was to create a safe, defined microbiota that could be implemented into our husbandry protocol to aid in the reproducibility of the SCID pig model.

## 2 Materials and Methods

### 2.1 Ethics statements

All procedures involving animals were conducted after approval by the Iowa State University Institutional Animal Care and Use Committee.

### 2.2 Experimental Animals

Yorkshire-Landrace gilts were raised on site at the Iowa State University campus. These females were artificially inseminated with thawed, frozen semen previously collected from boars also raised at Iowa State University. Specific genotypes used for each litter are discussed below.

Sows were moved at day 114 of gestation to the Iowa State University Laboratory Animal Resource (LAR) facility. They were prepared for cesarean section by Iowa State University LAR care staff, which included washing the sow.

Sows for cesarean section were sedated with intramuscular acepromazine (MWI Animal Health, Boise, ID) in the housing room 30-45 minutes prior to moving to the surgery preparation room. Once transported to the preparation room, sows received 0.2 mg/kg intramuscular morphine sulphate (Spectrum Chemical, New Brunswick, NJ) and topical lidocaine cream (Alembic Pharmaceuticals Ltd., Vadodara, Gujarat, India) was applied to the ears and epidural site. After approximately 10-15 minutes contact time for the lidocaine cream, about 35 ml of lidocaine (MWI Animal Health, Boise, ID) was administered via epidural, as described (Swindle & Smith, 2015). An ear catheter was placed (Kaiser-Vry et al., 2023) and propofol (Zoetis Inc., Kalamazoo, MI) administered IV to effect (500-800 mg). Once a sufficient anesthetic plane was achieved, the sow was intubated, and anesthesia maintained on isoflurane administered with oxygen. The surgical site was shaved, and an initial scrub performed in the preparation room.

The sow was moved to the surgery room and placed in lateral recumbency on the surgery table. The sow was maintained on IV fluids of Lactated Ringers Solution +/- 5% dextrose (Baxter International, Deerfield, IL) for the duration of the surgery. Once properly positioned, a complete surgical scrub was performed. Spray-on adhesive was used to secure a homemade flexible film surgical isolator to the sow. The surgical isolator and contents had been previously autoclaved where possible (surgical instruments and metal transfer cylinder) and sterilized with Clidox spray (Pharmacal Research Laboratories, Waterbury, CT). The surgical isolator allowed gloved access for the surgeon and surgery assistant as well as four people to revive piglets. A paramedian incision was made into the abdominal cavity and the uterus exteriorized. Incisions were made over each piglet, as each piglet was removed the umbilicus was clamped, and 1-2 drops naloxone (Somerset Therapeutics LLC, Hollywood, FL) placed on the tongue to reverse the anesthetic allowing piglets to be revived more promptly. Piglets were stimulated to assist with revival, warmed with towels and handwarmers. Any piglet that was still sedated approximately 10 minutes after removal from the uterus, received an additional 1-2 drops naloxone. The sow was euthanized with Fatal-Plus Solution (Vortech Pharmaceuticals, Dearborn, MI) after all piglets had been removed. Once all piglets were stable, piglets were moved to a transfer cylinder for transport to their housing environment.

The isolators used for designated litters were previously described (Aluthge et al., 2020). The BioBubble biocontainment housing environment for SCID pigs and the sterilization protocol of the biocontainment environment have also been previously detailed (Powell et al., 2018). A modification to the previously described sterilization protocol was the addition of C. diff Solution Tablets (3M, Athlone, Ireland). The tablets were dissolved in water and, using a foam sprayer, was sprayed on all surfaces in the bubble. After all surfaces air dried, the room was sprayed down with water. This change to the sterilization protocol was added prior to the third animal trial discussed in this article.

Once piglets were received into their selected environment, they were processed. All piglets received one milliliter of Gleptoforte injectable pig iron (Ceva Animal Health, Services, Lenexa, KS) intramuscularly, had needle teeth clipped, temperature programmable microchips (Unified Information Devices, Lake Villa, IL) injected sub-dermally for identification purposes, temperatures collected via microchips, and umbilical cords tied off with suture. A small piece of umbilical tissue was collected using scissors, which were disinfected with Oxivir Tb Wipes (Diversity, Fort Mill, SC) between collections, then placed in sterile 1.5 ml microcentrifuge tubes. Genotyping was performed using the DNeasy Blood and Tissue Kit (Qiagen, Germantown, MD) to purify genomic DNA and determine the genotype of the piglets. As genotyping results take more than a day to be returned, a preliminary method to rapidly determine which piglets were SCID and non-SCID was utilized. One milliliter of blood was drawn from the jugular vein of each piglet and a complete blood count (CBC) was performed to look at lymphocyte levels of each piglet. Within the experimental herd at ISU, we have found that if a piglet has less than 1.0 lymphocytes per microliter it is very likely to be SCID, which allows placing an animal in a treatment group when immediately necessary. An initial fecal swab was also collected prior to feeding and microbiota inoculation. The rest of the fecal swab collection protocol will be discussed in the sample collection section.

After piglets were processed, the microbiota consortium samples were given by oral gavage catheterization. Five milliliters of inoculum were syringed through the catheter followed by five milliliters of colostrum to ensure all inoculum was flushed through the catheter into the stomach. This process was then performed again approximately 24 hours later.

After the initial fecal swab was collected, irradiated colostrum powder (Bovine IgG Calf’s Choice Total Gold Colostrum, SCCL, Saskatoon, Saskatchewan) was mixed with sterile water and piglets were fed using baby bottles. The goal was to feed 20 to 25 milliliters of colostrum to each piglet, every thirty minutes to provide up to 250-300 milliliters in the first 36 hours. The amount of colostrum the piglets received at each feeding was recorded. The SCID piglets received 100% colostrum by care staff for the first 36 hours of life. A bowl of colostrum was added into the piglet’s environment at 12 hours of life to encourage drinking on their own. Once a piglet was observed drinking from the bowl, they were no longer offered the bottle. After 36 hours, piglets started receiving a mixture containing 50% colostrum and 50% milk replacer (irradiated Birthright Pig Milk Replacer powder (Ralco, Marshall, MN) appropriately mixed with water) until day 7. From one week to three weeks of age, piglets received a mixture of 25% colostrum and 75% milk replacer. Table 1 outlines the feeding timeline protocol.

**Table 1.** Feeding timeline for cesarean-section derived SCID piglets.

|  | Day 0 | Day 1-before<br>36 hours | Day 1-after 36<br>hours | Day 2 | Day 3 | Day 4 | Day 5 | Day 6 |
| --- | --- | --- | --- | --- | --- | --- | --- | --- |
| Colostrum | 100% | 100% | 50% | 50% | 50% | 50% | 50% | 50% |
| Milk<br>Replacer | 0% | 0% | 50% | 50% | 50% | 50% | 50% | 50% |
|  | Day 7 | Day 8 | Day 9 | Day 10 | Day 11 | Day 12 | Day 13 | Day 14 |
| Colostrum | 25% | 25% | 25% | 25% | 25% | 25% | 25% | 25% |
| Milk<br>Replacer | 75% | 75% | 75% | 75% | 75% | 75% | 75% | 75% |

Note: Bowls of colostrum and milk replacer were introduced to piglets at 12 hours of life.

Throughout the first 72 hours, piglets were continuously monitored. Piglet temperatures were recorded every hour to monitor for signs of infection and ensure the environment was an appropriate temperature for the piglets. If any abnormal behavior was noted, an LAR on-call veterinarian was alerted and further action was directed (Powell et al., 2018). Piglets were weighed once a day to ensure they were gaining weight appropriately.

The litters included for analysis were chosen based on husbandry similarities and their genetic heritage. The only major change between litters is the defined microbiota inoculum they received (Table 2).

**Table 2.** Microbiota treatment and genotype for each litter.

|  | Genotype |  | Microbiota |
| --- | --- | --- | --- |
| Litter 17 | A-SCID | n=2 | 2-strain |
|  | G-SCID | n=2 |  |
|  | Non-SCID | n=1 |  |
| Litter 23 | A-SCID | n=2 | 6-strain |
|  | Non-SCID | n=2 |  |

|  | Genotype |  | Microbiota |
| --- | --- | --- | --- |
| Litters 33 & 34 | A-SCID | n=2 | High<br>concentration<br>7-strain |
|  | G-SCID | n=5 |  |
|  | Non-SCID | n=1 |  |

Note: Artemis SCID denoted as A-SCID and IL2RG SCID denoted as G-SCID.

#### 2.2.1 Litter 17

A double heterozygous dam (ART^+/-^ IL2RG ^+/-^) was bred to an Artemis heterozygous sire (ART ^+/-^). This breeding resulted in two ART^-/-^ SCID piglets (A-SCID), two Il2RG^-/-^ SCID piglets (G-SCID), and one non-SCID pig used for this project. Piglets were placed into isolators for the duration of this trial. All piglets received the 2-strain microbiota inoculum.

#### 2.2.2 Litter 23

A heterozygous dam for the Artemis gene (ART ^+/-^) was bred to the same sire described in litter 17 (ART ^+/-^). Offspring from this breeding were two A-SCIDs and two non-SCID pigs used for this project. Like litter 17, this litter resided in isolators for the duration of this trial. All four piglets received the 6-strain microbiota inoculum.

#### 2.2.3 Litters 33 and 34

The dams used to create litters 33 and 34 were double heterozygous for the Artemis gene (ART^+/-^ IL2RG ^+/-^). These females were bred to a double mutant (BMT)- rescued SCID male (ART^-/-^ IL2RG ^-/Y^). This SCID male was born in a litter created by embryo transfer then had a bone marrow transplant performed which led to the reconstitution of T, B, and NK cells (Boettcher et al., 2020). Piglets for this trial were placed into the BioBubble biocontainment room and remained there for the duration of the trial.

The total number of piglets born across these two litters was 20 piglets. A fecal swab was collected during day one processing from all 20 piglets, to determine gnotobiotic status of the piglets. For the overall analysis, we only used piglets that met all timepoints we wanted to include. This included two A-SCIDs, five G-SCIDs, and one non-SCID pig. All piglets received the 7-strain microbiota consortium that had an increased concentration of bacteria, compared to the other two groups. The 7-strain microbiota inoculum contained ∼10^8^ CFU/mL for each isolate while the 2-strain and 6-strain inoculum contained ∼10^6^ CFU/mL for each isolate.

Environmental swabs were collected and analyzed for this trial to determine if environmental bacterial contaminants were present. These swabs were collected prior to the entry of piglets and again from the same location approximately six hours later. Samples were collected by swabbing a 10 cm x 10 cm section of all four floor locations in the bubble, the cart in which supplies were stored and the buses where the piglets are raised.

### 2.3 Development of the 2, 6, and 7-strain Inocula

*Clostridioides difficile* is a frequent member of neonatal pig gut microbiota and is present in two-thirds of litters and one-third of piglets (Songer, 2004). Using the snatch farrowing method mentioned above, the piglet’s microbiota was largely acquired from the sow and the farrowing environment (Boettcher & Tuggle, 2022). This method resulted in numerous cases of *C. difficile* infection which prompted a switch to sterile cesarean delivery and the development of a defined microbiota that protected against *C. difficile* proliferation. Huang and colleagues (2018) designed a defined commensal microbiota (DMF) for gnotobiotic piglets, to represent the infant gut microbiota, to further their studies (Huang et al., 2018). Their methods guided us towards adapting their DMF to a defined microbiota for use in SCID pig.

### 2.4 Cultivation of the inoculum

All handling and incubation steps were completed in anaerobic conditions (5/5/90% H2/CO2/N2 Mixed Gas, Airgas, Radnor, PA). -80°C stock cultures (preserved with either 20% glycerol or 7% DMSO) of the isolates were plated on the media listed in Table 3. De Man – Rogosa – Sharpe (MRS) agar (Thermo Fisher Scientific, 52.0 g/L, + 20 g/L agar), Brain Heart Infusion (BHI) agar (Thermo Fisher Scientific, 37 g/L, + 20 g/L agar), and BHI + 1% Yeast Extract (BD, 10 g/L + 20 g/L agar) agar plates were pre-reduced in anaerobic conditions prior to use. Anaerobic liquid media was prepared following the methods described (Uchino & Ken- Ichiro, 2013).

**Table 3.**
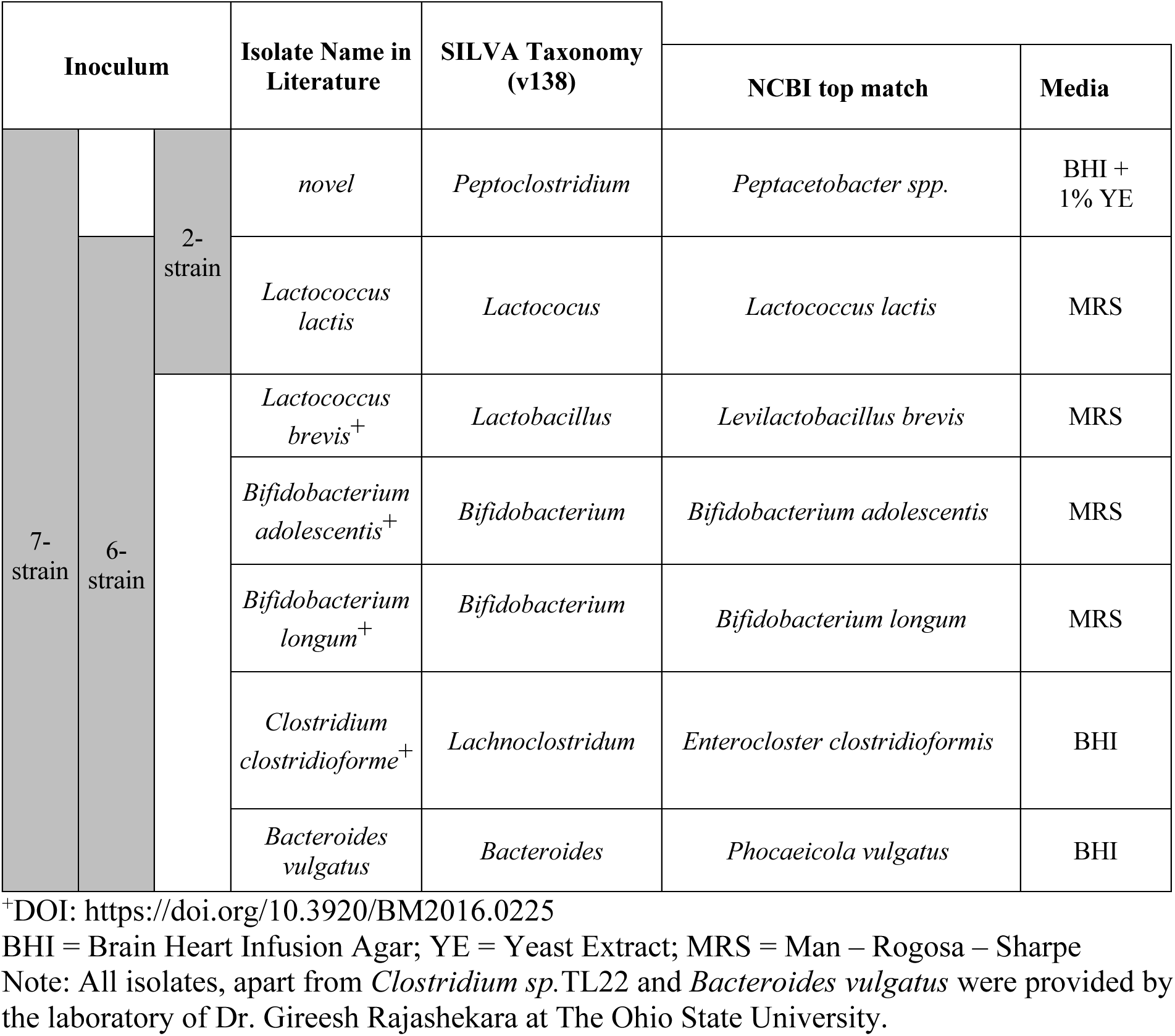
Classification of each isolate in the inoculum from 16S rRNA gene sequencing.

Culture purity was routinely confirmed by sequencing the 16S rRNA gene of each isolate. To obtain genomic DNA, a small scraping of cells from an agar plate was suspended in 20 μL of 10 mM EDTA in a microtube. After vortexing, the microtube was heated at 100°C for 10 minutes. The microtube was then centrifuged for 10 minutes at 14.6 x g. After centrifugation, 15 μL of supernatant was transferred to a new microtube containing 75 μL of PCR-grade diethyl pyrocarbonate (DEPC) water. 16S rRNA gene PCR was completed using the primers 616F (5’ –AGA GTT TGA TYM TGG CTC – 3’) and 1492R (5’ – GGY TAC CTT GTT ACG ACT T –3’) following the standard Platinum II Taq Hot-Start DNA polymerase protocol (Thermo Fisher Scientific). The PCR products were confirmed using agarose gel electrophoresis, then purified using the Thermo Scientific GeneJET PCR Purification Kit (Thermo Fisher Scientific). The purified DNA was sent to the ISU DNA facility where it underwent Sanger sequencing. The resulting16S rRNA gene sequences were assigned to known species using NCBI BLAST. The top match for each isolate can be found in Table 3.

The 2-strain and 6-strain inocula were created using the following method. For each isolate individually, a 10 μL loopful of cells was used to inoculate 10 mL of pre-reduced broth (corresponding media type for each isolate listed in Table 3). The liquid cultures were incubated for 20 hours at 37°C yielding ∼10^8^ – 10^9^ CFU/mL. To check for contamination, 10 μL of each liquid culture was used to inoculate agar plates in triplicate. The plates were incubated in aerobic and anaerobic conditions for 48 hours at 37°C and monitored for growth. If abnormal colony morphologies were observed, the identities of the potential contaminants would be confirmed by sequencing the 16S rRNA gene. Cultures were discarded if contamination was found. The clean liquid cultures were then combined in a sterile 500 mL bottle and diluted with an equivalent volume of BHI broth. Sterile glycerol was then added to reach a 20% (v/v) glycerol solution.

Aliquots of the mixed inoculum were stored in 50 mL centrifuge tubes at -80°C. Prior to use, the tubes were thawed at 37°C for 10 minutes and were administered by oral gavage within 30 minutes of thawing.

Once thawed, the surviving count of each isolate was ∼10^6^ CFU/mL. For the 7-strain inoculum, the initial bacterial load was increased to ∼10^8^ CFU/mL for each isolate. To achieve this, 2.5 mL of the initial 10 mL broth cultures were used to inoculate 50 mL of broth (Table 3). The 50 mL cultures were incubated at 37°C for 36 hours yielding ∼10^8^ – 10^9^ CFU/mL. To confirm culture purity, samples were plated on solid media as described above. The liquid cultures were then mixed in a sterile disposable 1 L plastic bottle, and sterile glycerol was added to reach a 20% (v/v) glycerol solution. 25 mL aliquots of the mixed inoculum were stored in 50 mL centrifuge tubes at -80°C.

### 2.5 Sample collection

Swabs were collected at six timepoints: while piglets were being processed after cesarean section, prior to microbiota inoculation which is designated as day 0, 24 hours after the second inoculation of microbiota, three days after the second inoculation, seven days after the second inoculation, and 14 days after the second inoculation. See Table 4 for the sample collection timeline.

**Table 4.** Fecal swab collection timeline.

| Day 0 (Date of birth) | Day 1 | Day 2 | Day 4 | Day 8 | Day 15 |
| --- | --- | --- | --- | --- | --- |
| Fecal swab collected prior to inoculation | **2<sup>nd</sup> microbiota inoculation** | Fecal swab collected | Fecal swab collected | Fecal swab collected | Fecal swab collected |
| **1<sup>st</sup> microbiota inoculation** |  |  |  |  |  |

Puritan Fecal Opti-Swabs Elongated Tip Swab, 2 ml Cary Blair Medium (Harmony, Garden Grove, CA) were used for sample collection. Fecal swabs were inserted into the anus, taking care to not touch anywhere else, until the tip was fully inserted, approximately 2 cm. The swab was then gently spun, removed from the anus, inserted into the collection vial, and tightly secured. Immediately after collection, the fecal swabs were stored at -80 °C until DNA extractions were performed. Fecal swabs collected during piglet processing were streaked on BHI media to determine gnotobiotic status.

Samples collected up to day 15 were used for analysis. These were the last swabs used for analysis as weaning occurs after this collection time point and dry feed is introduced to the piglets, which is known to change the gut microbiota composition (Frese et al., 2015; Guevarra et al., 2019). Fecal swabs were not used in the analysis if they were taken after piglets received antibiotics.

### 2.6 Processing of samples and DNA extraction

Fecal swabs were removed from the freezer, thawed at room temperature, and vortexed for 10 minutes to detach fecal material from the swab. Lids were then removed from the vials, and the swab was removed from the vial using forceps. The swab was discarded, and the cap replaced. Between each swab, forceps were dipped in bleach and rinsed in sterile deionized water to avoid cross contamination. Fecal swab tubes were then centrifuged for three minutes at 4,694 x g at 4°C (Cassas et al., 2024). The supernatant was then discarded, being careful not to disturb the pellet.

DNA extraction from the fecal swabs was then performed following the protocol provided in the DNeasy Powerlyzer Powersoil Kit (Qiagen). Upon completion of the DNA extraction protocol, the concentration of the samples was determined using a ND-100 Nanodrop spectrophotometer (NanoDrop Technologies, Dockland, DE). Many of the samples, especially from the earlier timepoints, were found to have low DNA concentrations (∼ 10 ng/µl or less). To ensure the DNA could be amplified prior to amplicon sequencing, samples with the lowest DNA concentrations underwent 16S rRNA gene PCR. Successful amplification was confirmed with agarose gel electrophoresis. Although many samples had low DNA concentrations, all samples showed DNA amplification allowing them to be submitted for Illumina MiSeq 16S rRNA gene amplicon sequencing. Samples with DNA concentrations higher than 25 ng/µL were diluted to reach the desired concentration of 25 ± 5 ng/µL.

The processing method described above was the same for all three trials and litters.

Additional control samples were added to the DNA extraction process for Litter 33 and Litter 34 samples. An unused fecal swab that was never frozen and an unused fecal swab that was frozen at -80°C were included for DNA extraction at the end of each kit as controls and were processed like the rest of the samples. This was to determine if contaminants were present in the blank fecal swabs. Two samples of PCR-grade DEPC water were also added to the end of the plate submitted for sequencing, to determine if there were any contaminants contracted during sequencing.

### 2.7 16S rRNA gene amplicon sequencing and data processing

16S rRNA gene amplicon sequencing of the DNA obtained from SCID piglet fecal swabs was completed using the Illumina MiSeq platform at the ISU DNA facility. The ISU DNA facility performed custom library preparation to amplify the 16S rRNA gene V4 region.

Universal 16S rRNA gene bacterial primers 515F (5′-GTGYCAGCMGCCGCGGTAA-3′) and 806RB (5′-GGACTACNVGGGTWTCTAAT-3′), and the Platinum Hot Start PCR Master Mix (2x) (Thermo Fisher Scientific) were used to amplify the V4 variable region with the following thermocycler conditions: initial denaturation step at 94 °C for 3 min; 45 s of denaturing at 94 °C; 60 s of annealing at 50 °C; 90 s of extension at 72 °C. This was repeated for 35 cycles and finished with a 10 min extension at 72 °C. After PCR, equal amounts of amplicons from each sample were pooled into a single tube and purified using the standard protocol of the UltraClean PCR Clean-Up Kit (MO BIO Laboratories). All described reagents were dispensed using a Mantis robot (Formulamatrix). The barcoded amplicons then underwent paired-end (2 x 250) 500-cycle sequencing on an Illumina MiSeq platform.

The 16S rRNA gene amplicon sequences for litters 17 and 23 were processed and analyzed separately from litters 33 and 34 using the following protocol. Raw sequences were processed using mothur (v1.43.0) (Schloss et al., 2009) following a protocol based on the MiSeq Standard Operating Procedure (Kozich et al., 2013). The “make.contigs” command was used to merge and filter the paired-end reads. Parameters including a maximum homopolymer run of eight bp, a minimum length of 252 bp, and a cutoff of zero ambiguities were applied for the “screen.seqs” command. Sequences were aligned using the SILVA reference database (v138) (Quast et al., 2013) and the “align.seqs” command. Chimeric sequences were removed using the “chimera.vsearch” command in combination with the SILVA.gold reference. *De novo* operational taxonomic unit (OTU) clustering at 99% gene similarity was completed and the resulting OTUs were classified using the SILVA reference database (v138). The resulting OTU data was imported into R where the package decontam (v1.20.0) (Davis et al., 2018) was used to identify and remove contaminating DNA. Microbial community visualization was completed using the Phyloseq (v1.38.0) (McMurdie & Holmes, 2013), vegan (v2.5.7) (Oksanen et al., 2022), and ggplot2 (v3.4.0) (Wickham, 2016) R packages.

### 2.8 Statistical Analysis

OTUs with fewer than 10 reads were removed prior to statistical analysis. The microbial raw abundance data was normalized to minimize sequencing depth biases by dividing by the corresponding sample library size, resulting in “relative abundance” values used for further analysis. The following model was used to evaluate the effects of genotype and time on the beta diversity and alpha diversity of the piglet’s microbiota:

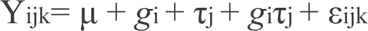

Where Y*ijk* is the observed value for k^th^ experimental unit within the i^th^ level of genotype (A- SCID, G-SCID, or non-SCID) at the j^th^ timepoint (Day 2, Day 4, Day 7-8, or Day 14-15); µ is the overall mean; *gi* is the fixed effect of the i^th^ genotype (i = A-SCID, G-SCID, or non-SCID); τ*j* is the fixed effect of the j^th^ timepoint (j = Day 2, Day 4, Day 7-8, or Day 14-15); *gi*τ*j* is the interaction of genotype and timepoint; and *εijk* is the error as described by the model for Yijk.

Differences in beta diversity were analyzed using Bray-Curtis distances and visualized using principal coordinate analysis (PCoA) plots using the Phyloseq package in R. A permutational multivariate analysis of variance (PERMANOVA) and a permutational multivariate analysis of dispersion was completed with the commands “adonis2” and “betadisper” from the vegan package in R (Oksanen et al., 2022). The resulting p-values were adjusted to account for multiple-comparisons using Bonferroni’s correction (Dunn, 1961).

The Phyloseq package in R was used to generate alpha diversity measurements for the number of observed species, Chao species richness, Simpson evenness, and Shannon diversity. In addition to the described model, Pig ID was included as a repeated measure to account for the covariance among the samples taken from the same animal. The least square means (LSmeans) of all alpha diversity measurements were compared using the PROC MIXED procedure in SAS (Version 9.4, SAS Inst., Cary, NC). The resulting p-values from the pairwise comparisons were corrected for multiple-comparisons using Tukey’s Honest Significant Difference test (Tukey, 1949) Adjusted p-values were considered significant if p < 0.05.

## 3 Results

### 3.1 Description of strains used in the defined microbiota and sample collection

To determine which defined microbiota was most stable in the gut microbiome, SCID piglets were c-sectioned and inoculated with different versions of a defined microbiota. The complexity of the microbiota was gradually increased over time, starting with a 2-strain microbiota, then a 6-strain microbiota, and finally a 7-strain microbiota. The list of strains included in each inoculum can be found in Table 3. We sought to balance the risk associated with opportunistic infections from otherwise normal porcine microflora while simultaneously mitigating the negative effects of an absent gastrointestinal microbial community.

*Peptacetobacter spp.*, applied in the 2-strain and 7-strain inoculum, is a novel bacterial strain that converts primary bile acids. To avoid potential overgrowth of a *Peptacetobacter spp.* monoculture, *Lactococcus lactis* was included in the 2-strain inoculum. Research has shown that *L. lactis* is beneficial for piglets post-weaning, resulting in improved growth performance and intestinal immunity among other benefits (Yu et al., 2021). In the 6 and 7- strain inocula, *Lactobacillus brevis*, *Bifidobacterium adolescentis*, *Bifidobacterium longum*, and *Clostridium clostridioforme,* were efficacious as part of a defined commensal microbiota, or DMF as described by Huang and colleagues (2018), in a neonatal gnotobiotic pig model. *Bacteroides thetaiotaomicron* was also included in the described DMF, but *Bacteroides vulgatus* was substituted based on availability from the Iowa State University Veterinary Diagnostic Laboratory (ISU VDL).

As outlined above, fecal swabs were collected from SCID piglets in each litter as part of processing after birth and these samples were denoted as day 0. Fecal samples were then collected according to the protocol outlined in Table 4. Samples collected up to day 15 were included in statistical analysis as weaning occurs after this timepoint. Litter 17, however, only included samples collected up to day 8 as antibiotics were introduced prior to collection at the last time point. Environmental swabs were also collected from the BioBubble biocontainment facility to confirm sterility of the housing environment prior to piglet entry. DNA extraction and 16S rRNA amplicon sequencing were then performed for all collected swabs.

### 3.2 Combined analysis of sample, control, and environment swabs led to exclusion of day 0 samples from further analysis

Figure 1 is a combined Principal Coordinate Analysis (PCoA) plot which includes data from all fecal swab collections for litters 17, 23, 33, and 34, controls samples, environmental samples and swabs collected from the sows of Litter 33 and 34. For statistical purposes, samples collected from Litter 17 and 23 were grouped and labeled with the same color and Litter 33 and 34 samples were grouped and labeled with the same color. This means control swabs from Litter 17 and 23 were labeled together as one item and the same for day 0 swabs and piglet’s swabs.

**Figure 1.**
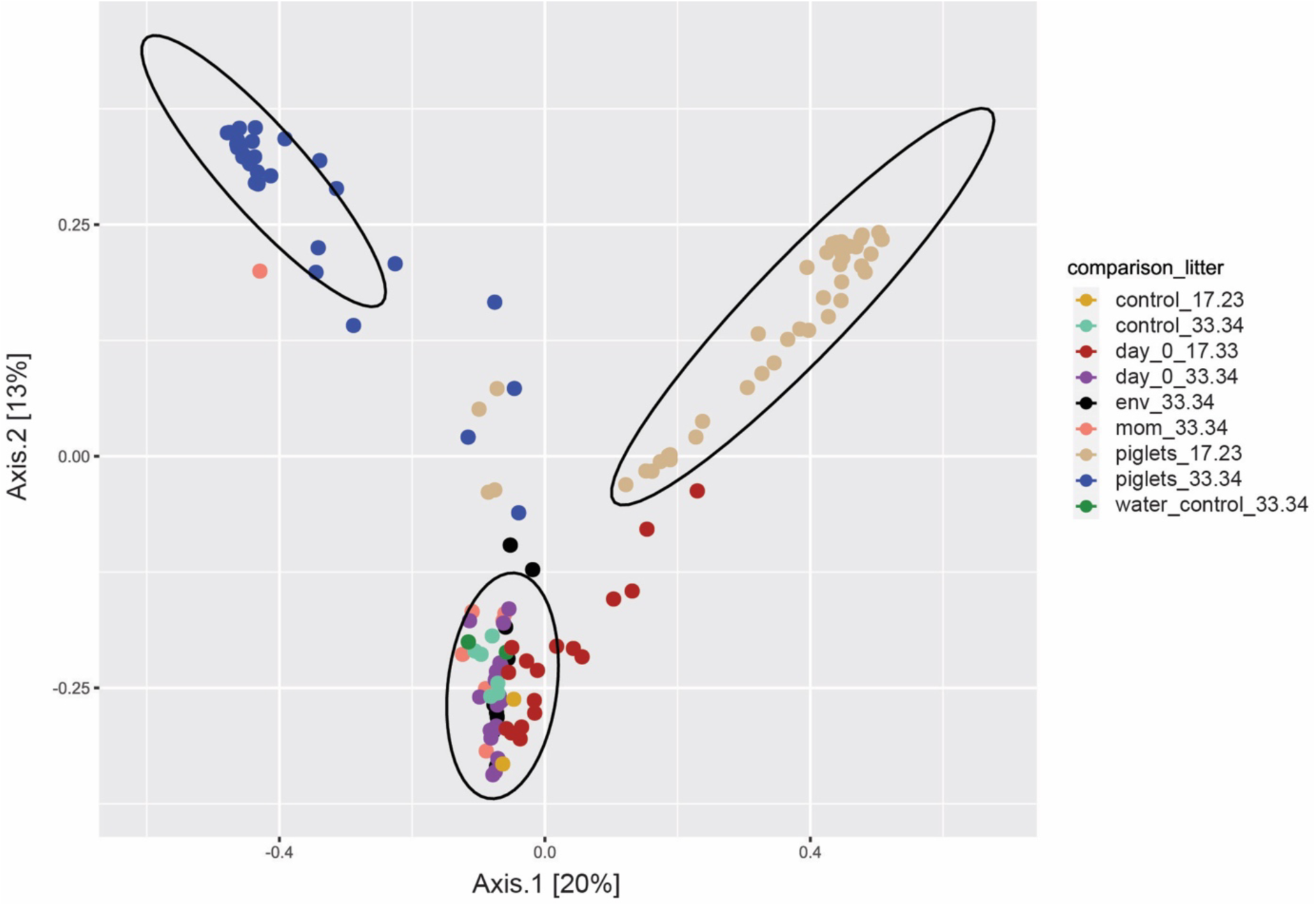
Principal coordinate analysis (PCoA) plot displays day 0 timepoint clustering with controls and environmental swabs leading to the exclusion of the day 0 timepoint from analysis. Distances between samples denote Bray-Curtis dissimilarity measures based on 16S rRNA gene amplicon sequencing. This plot shows that the composition of day 0 samples, taken immediately before inoculum was given, largely cluster with the control swabs, environmental swabs, and swabs from the mother’s placenta. There was a significant difference between the non-day 0 swabs and the day 0 and control swabs (P=0.001) that were detected using PERMANOVA. There was also a significant difference between Litter 17 and 23 swabs, labeled a “piglets_17.23”, compared to Litter 33 and 34, labeled as “piglets_33.34”, (P=0.001) detected using PERMANOVA, so they were analyzed separately. Because of this, we removed the day 0 samples from the following analysis. More information about the controls and environmental swabs are provided in the materials and methods.

The same labeling system was followed for Litter 33 and 34 samples. Day 0 swabs were the swabs initially collected from the piglets and the data points labeled as “piglets” were the swabs collected after this initial timepoint. Environment swabs were labeled as “env” in Figure 1 and the location of collection for these swabs is provided in the materials and methods. The samples labeled as controls are DEPC water samples to test for contaminants during DNA extraction. The samples labeled as sow, were collected from the anus and vulva from the sows of Litter 33 and 34. Lastly, the samples labeled as water control were the DEPC water samples included at the end of the plate submitted for 16S rRNA gene amplicon sequencing.

At day 0, prior to microbiota inoculation, piglets are expected to be free of microorganisms; however, results from 16S rRNA amplicon sequencing revealed a minimal number of reads, with an average of 30,892 reads pre-decontam package (discussed in materials and methods), from these swabs. These reads could indicate the piglets were not germ free at the start of the experiment. Since the data indicated a low presence of bacterial DNA in these samples, we analyzed day 0 swabs separately from the later timepoints. The purpose was to look at the relationship between day 0 swabs, the non-day 0 swabs labeled as “piglets”, the environmental swabs, and the control swabs. We found that the day 0 swabs, environmental swabs, and control swabs clustered together, and separately from the non-day 0 piglet swabs.

This indicated the read counts present for day 0 samples was likely background, indistinguishable from low levels of bacterial DNA present in the environment or process control swabs, and that no bacteria were present in the piglets or the environment. Thus, day 0 swabs were then omitted from future microbiome analysis.

There was a significant difference between the non-day 0 swabs and the day 0 and control swabs (P=0.001) that were detected using PERMANOVA. There was also a significant difference between Litter 17 and 23 swabs, labeled a “piglets_17.23”, compared to Litter 33 and 34, labeled as “piglets_33.34”, (P=0.001) detected using PERMANOVA. With these results, we removed the day 0 samples from analysis and the Litter 17 and 23 were analyzed separately from Litter 33 and 34.

Supplementary Figure 1 is an additional PCoA plot displaying only data collected during the third trial (litter 33 and 34). With the addition of environmental swabs and more control swabs, we wanted to identify the beta diversity of samples from this trial. This figure displays a clear difference in clustering of day 0, control, and environmental swabs when compared to the non-day 0 swabs denoted as “piglets” in the figure. Furthermore, the environmental and day 0 swabs collected from litter 33 and 34 were cultured, shortly after collection, to determine if any bacteria were present. This included six environmental swabs collected prior to piglet entry and all twenty swabs collected from the piglets on day 0. Two blank fecal swabs were also included to confirm their sterility. Each sample had one plate for aerobic growth and one plate for anaerobic growth. The environmental swabs collected prior to piglet entry, cultured in an aerobic environment, resulted in three plates with no growth, two plates with a single colony, and one plate that grew many colonies with many morphologies. The results from culturing in the anaerobic environment were two plates with no growth, one plate with three colonies, two plates that had multiple colonies, and one plate that had many colonies with many morphologies. Of the day 0 piglet swabs, one piglet sample out of twenty presented a single colony in aerobic conditions, it is possible this was contamination from collection the sample. The plates for the blank swabs did not have any growth. The cultivation results and addition of controls reinforced the decision to exclude day 0 swabs from further analysis.

### 3.3 No significant effect of SCID piglet genotype on beta diversity of piglet gut microbiota, regardless of piglet age

Figure 2 contains unconstrained PCoA plots for litter 17 (Figure 2A), litter 23 (Figure 2B), and litter 33 and 34 (Figure 2C) to examine the beta diversity of the microbiota of SCID piglets due to genotype, regardless of piglet age, due to the defined microbiota they received. Whole bacterial community beta diversity comparisons were performed using PERMANOVA for all groups.

**Figure 2.**
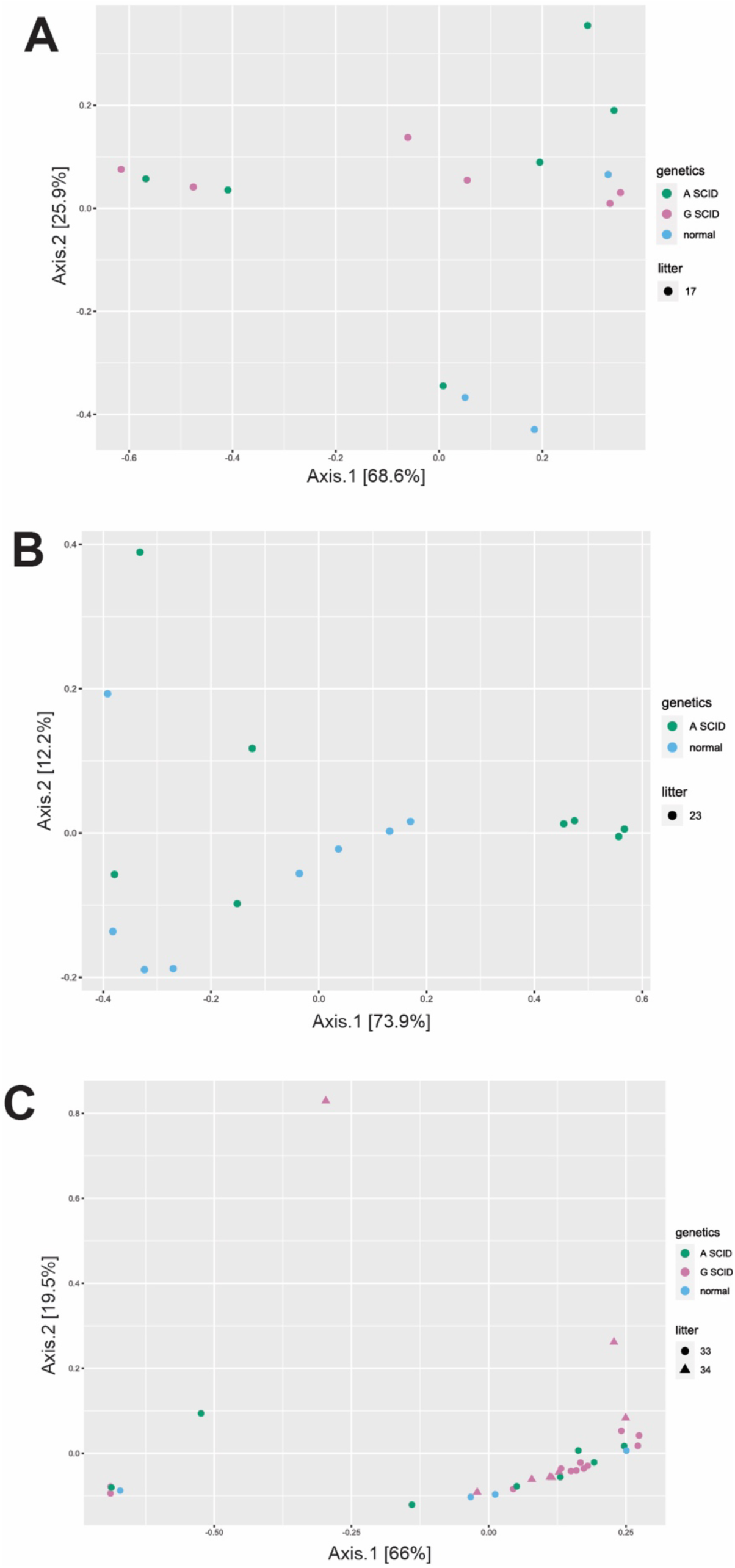
PCoA plots display no significant effect of SCID genotype on beta diversity of piglet gut microbiota when receiving the defined microbiota, regardless of piglet age. Based on 16S rRNA gene amplicon sequencing, distances between samples in PCoA plot denote Bray-Curtis dissimilarity measures. (A) Litter 17 received the 2-strain defined microbiota and there were significant differences due to genotype (P = 0.018) detected using PERMANOVA. However, no statistical differences in pairwise comparisons were observed across genotypes. This litter received antibiotics prior to collection of day 14-15 swabs so they were omitted from the analysis. (B) Litter 23 received the 6-strain defined microbiota and significant differences due to genotype (P = 0.028) were detected using PERMANOVA, however, unequal variance between groups was detected with PERMDISP2 (P = 0.009). (C) Litters 33 and 34 received the 7-strain defined microbiota and no significant differences due to genotype (P=0.340) were detected using PERMANOVA.

Litter 17 (2-strain microbiota inoculum) PERMANOVA results (P = 0.018) revealed a significant difference due to genotype. There were no significant pairwise comparisons noted in an attempt to determine specific genotype effects, which could be due to lack of power from the small sample size. The second trial, involving litter 23 (6-strain microbiota inoculum), revealed significant differences due to genotype using PERMANOVA (P = 0.028). A significant PERMDISP2 (P=0.009) test showed that there was unequal variance between the treatment groups, meaning assumptions for PERMANOVA testing were not met. Lastly litter 33 and 34 (7- strain microbiota inoculum) PERMANOVA results (P= 0.340) revealed no significant differences due to genotype.

### 3.4 Significant effect of piglet age on beta diversity of gut microbiota of SCID piglets, regardless of genotype

Figure 3 examines beta diversity of the gut microbiota of SCID piglets, regardless of genotype. PERMANOVA results for litter 17 (P = 0.001), litter 23 (P=0.008), and litter 33 and 34 (P= 0.001) revealed a significant different due to piglet age over time.

**Figure 3.**
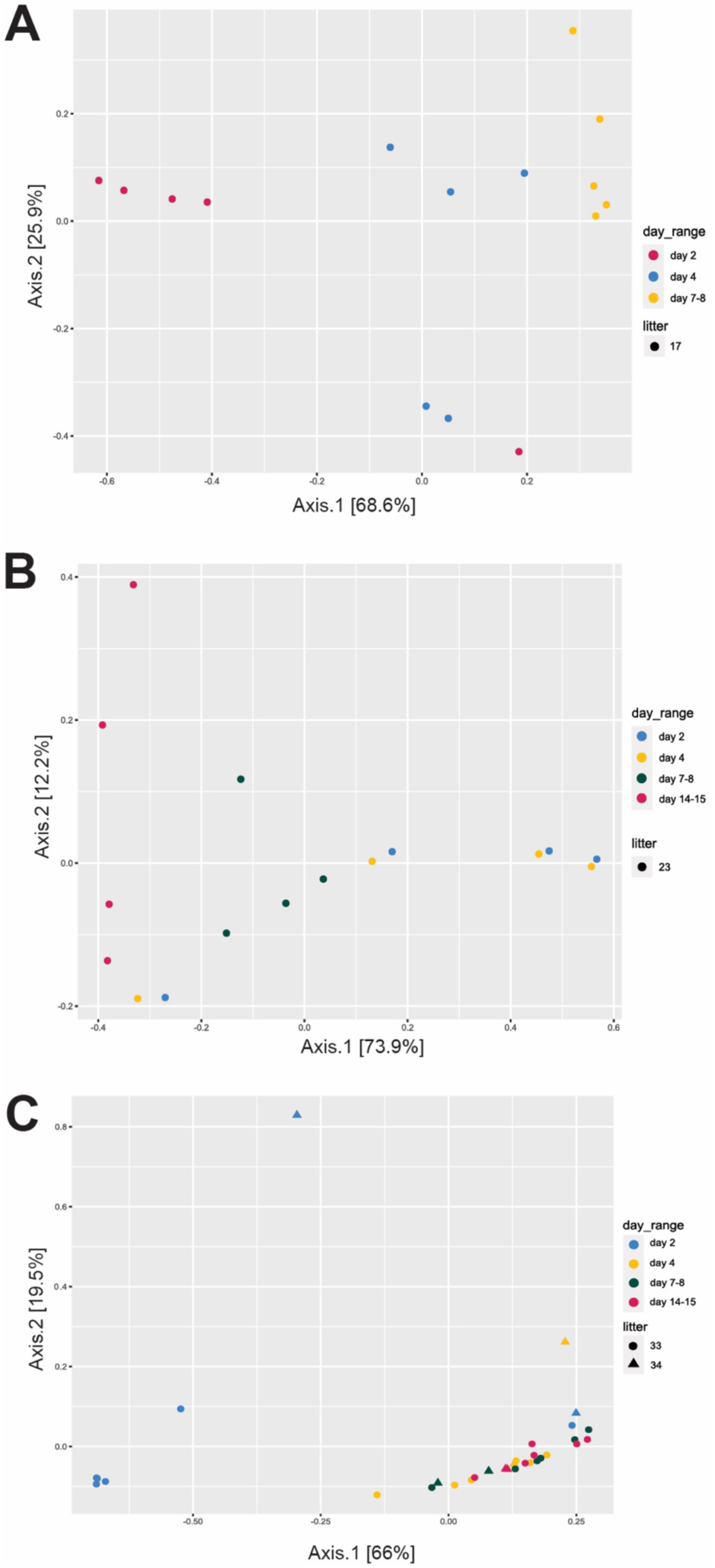
Significant effect of piglet age on beta diversity over time of gut microbiota of SCID piglets, regardless of genotype. Based on 16S rRNA gene amplicon sequencing, distances between samples in PCoA plot denote Bray-Curtis dissimilarity measures. (A) There were significant differences in the intestinal microbiota composition over time for litter 17, which received the 2-strain defined microbiota, due to piglet age (P = 0.001) detected using PERMANOVA. This litter received antibiotics prior to the collection of day 14-15 swabs, so they were omitted from analysis. Pairwise comparison results can be seen in the results section. (B) Significant differences in microbiota composition over time for litter 23, which received the 6-strain defined microbiota, due to piglet age (P = 0.008) were detected using PERMANOVA. Pairwise comparison results can be seen in the results section. (C) Significant differences in microbiota composition over time for litters 33 and 34, which received the 7-strain defined microbiota, due to piglet age (P = 0.001) were detected using PERMANOVA. Pairwise comparison results can be seen in the results section.

Pairwise comparisons for Litter 17 (Figure 3A) revealed a significant difference between day 4 and day 7-8 (P=0.027). Litter 23 (Figure 3B) did not have any significant pairwise comparisons noted for a direct comparison of piglet age, presumably due to lack of power from a small sample size. Lastly, litter 33 and 34 (Figure 3C) pairwise comparisons revealed a significant difference between day 2 and day 14-15 (P=0.018) and a significant difference between day 4 and day 14-15 (P=0.012).

### 3.5 Increase in inoculum complexity and concentration appears to offer greater microbiota stability over time, regardless of genotype

Based on 16S rRNA gene amplicon sequencing, the relative abundance of each inoculated bacteria for all timepoints from the three trials can be seen in Figure 4. This aids in the understanding of how the microbiome composition changes, over the sampling timepoints.

**Figure 4.**
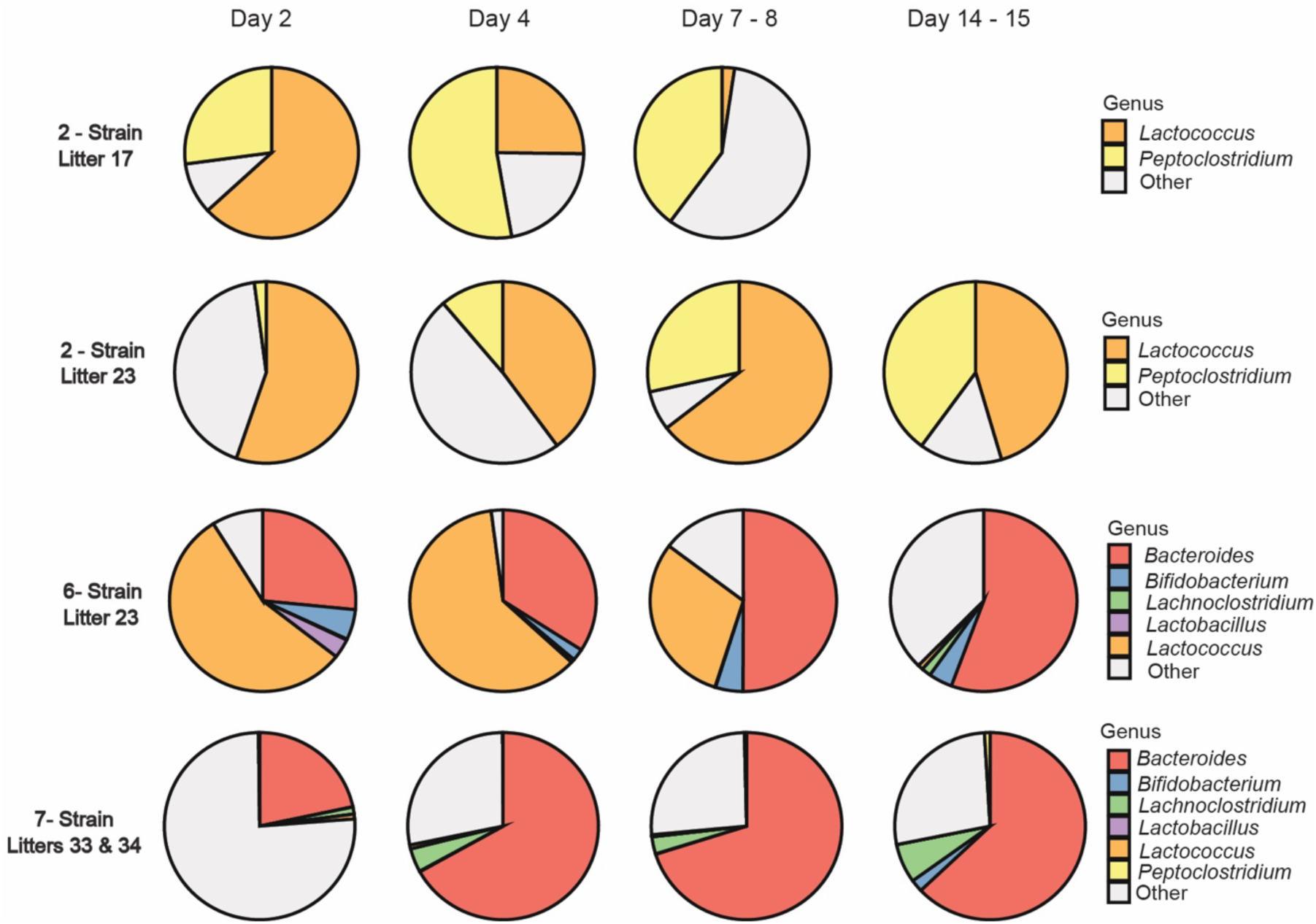
Increase in inoculum complexity and concentration appears to offer greater stability to the gut microbiota of SCID piglets over time. Based on 16S rRNA gene amplicon sequencing, this figure presents the relative abundance of each inoculated isolate when compared to the relative abundance of bacteria not administered in the initial inoculum (“Other”- see Supplementary Table 1, 2, and 3, for the list of bacteria included in this category for each group). Litter 17 (2-strain inoculum) received antibiotics prior to the collection of day 14-15 swabs. Due to the potential effects of antibiotic administration on the microbiota these samples were excluded. It should be noted that the initial administered concentration of the 7-strain inoculum is higher than the 2-strain and 6-strain inocula. The fraction of “Other” bacteria remained relatively stable after day 2 in the piglets that received the 7-strain inoculum.

Litter 17 received the 2-strain inoculum which included *Peptacetobacter spp.* and *Lactococcus lactis*. At day 2, both bacteria were present, with the relative abundance of *L. lactis* being the greatest. A shift is seen at day 4 where the relative abundance of *Peptacetobacter spp.* and other bacteria increased. At day 7-8, the relative abundance of *L. lactis* had decreased significantly, *Peptacetobacter spp.* decreased slightly, and other bacteria increased. By day 7-8, other bacteria, that were not part of the 2-strain inoculum, were most abundant.

Litter 23 received the 6-strain inoculum which included *Bacteroides vulgatus, Bifidobacterium adolescentis* and *Bifidobacterium longum* (which were grouped together in Figure 4), *Clostridium clostridioforme* (denoted as *Lachnoclostridium* in Figure 4, as this is the SILVA Taxonomy name*), Lactobacillus brevis*, and *Lactococcus lactis*. At day 2, the relative abundance of *L. lactis* was greatest followed by *B. vulgatus*. *Bifidobacterium* and *L. brevis* were present in small percentages and other bacteria that was not inoculated were also present. At day 4, the relative abundance of *L. lactis* and *B. vulgatus* increased while there was a decrease in *Bifidobacterium*, *L. brevis* and other bacteria. At day 7-8, *B. vulgatus*, *Bifidobacterium* and other bacteria increased in relative abundance, and *L. lactis* decreased. By day 14-15, the relative abundance of *B. vulgatus*, *L. clostridioforme* and other bacteria had increased, while there was a major decrease in *L. lactis*. *Bifidobacterium* abundance was unchanged. Other, not inoculated, bacteria appear to flourish by day 14-15; however, compared to the 2-strain microbiota at day 7- 8, the relative abundance of inoculated species increased when using the 6-strain inoculum.

Lastly, Litter 33 and 34 received the 7-strain inoculum which included all bacteria in the 6-strain as well as *Peptacetobacter spp.*, which was seen in the 2-strain. As well, a 100x increase in the concentration of the bacteria in this inoculum compared to the prior two was used. At day 2, which is after the second microbiota inoculation, the majority of the relative abundance was other bacteria; however, *B. vulgatus*, *L. clostridioforme* and *L. lactis* were present. By day 4, the relative abundance of *B. vulgatus* and *L. clostridioforme* increased while *L. lactis* and other bacteria decreased. Day 7-8 composition was like day 4 with the addition of *Peptacetobacter spp..* Finally, by day 14-15, the relative abundance of *Peptacetobacter spp.*, *L. clostridioforme*, and *Bifidobacterium* increased while *L. lactis* and *B. vulgatus* had decreased. The percentage of other bacteria varied little. *L.brevis,* was inoculated as part of the 7-strain defined microbiota but was not present in any of the relative abundance timepoints in Figure 4.

The relative abundance of other, not inoculated, bacteria was greater at day 2 when inoculated with the 7-strain, compared to day 2 for the 2-strain and 6-strain inoculum. However, the other three timepoints for the 7-strain inoculum were most consistent to each other compared to the other two groups.

## 4 Discussion

There has been minimal literature published using SCID pig models due to the recent discovery and creation of such models, as well as complexities involved in the rearing of these highly disease-susceptible pigs. It is known that early establishment of the gut microbiota is important for the development of a healthy gut microbiota, which also is known to have a significant interaction with the immune system (Kelly et al., 2007). To our knowledge, a defined microbiota has not been used in a protocol for rearing SCID piglets, but defined microbiota consortia have been used in other animal model systems, such as gnotobiotic pigs (Huang et al., 2018; Laycock et al., 2012) We investigated three different defined microbial consortia and the effect of SCID genotype and piglet age have on the gut microbial composition of piglets pre-weaning. The bacterial strains used for these consortia were adapted from a DMF previously used to create an environment that mimicked the infant gut microbiota (Huang et al., 2018).

The primary purpose of this study was to test different levels of complexity of defined microbiota consortia and determine if there was an effect on the gut microbiota due to genotype or piglet age. A secondary outcome from the results of this study was to develop a defined microbiota that was safe for use in SCID piglets and that could be standardized and reproduced easily.

All litters were delivered by cesarean section, then transported to their sterile housing environment where they received their respective defined microbiota consortia (2-strain, 6-strain, and 7-strain) by oral gavage. Fecal swabs were collected according to the protocol outlined in the materials and methods. Day 0 swabs were not included in the analysis due to their similarity to control and environmental swabs, indicating there was little to no live bacteria present in the day 0 samples (Figure 1).

The composition of the litter 33 gut microbiota compared to the litter 34 gut microbiota was not found to be significantly different. Because of this, and the fact that the litters were born on the same day, had a common environment, and received the same 7-strain inoculum, litter 33 and 34 data were combined for analysis. Along with being the most complex inoculum, the 7- strain defined microbiota inoculum also had the highest concentration of bacteria at ∼10^8^ CFU/mL while the 2-strain and 6-strain defined microbiota inoculum had a concentration of ∼10^6^ CFU/mL. The consortia developed by Huang and colleagues (2018), utilized in gnotobiotic piglets, had a concentration of about 7 x10^5^ CFU/mL. With their positive outcome, we utilized a higher concentration at 2 x 10^6^ CFU/mL for the 2-strain defined microbiota and 6 x 10^6^ CFU/mL for the 6-strain (one million of each bacteria per mL of inoculum). Cheng and colleagues (2022) designed a complex defined microbiota for use in mice at a concentration of 10^9^ CFU/mL for each bacterial strain, with a total of 10^10^ bacteria in the 0.2 mL inoculum. This prompted the increase in total bacterial concentration for the 7-strain defined microbiota at 7 x 10^8^ CFU/mL; with a total of 3.5 x 10^9^ CFU in the 5 mL inoculum. Thus, this increase was thought to bring the inoculum closer to the equivalent dose given to mice (Cheng et al., 2022). However, since these mice were immunocompetent, an important goal with the increase in bacterial concentration of the 7-strain inoculum was to determine if it was safe for use in SCID piglets. Additionally, this increase was thought to be closer to what the equivalent dose would be in comparison to what is given to mice, given the large difference in intestinal size (Cheng et al., 2022). All three defined microbiota consortia (2-strain, 6-strain, 7-strain) were determined safe for use in the SCID piglets, including the 7-strain with the increase concentration of bacteria, as the inoculated bacteria was never found in blood collected for culture.

### 4.1 Piglet genotype does not have a significant effect on piglet gut microbiota

For all three inoculum, a significant effect due to genotype on the microbial composition was not significant. Litter 17, which received the 2-strain microbiota inoculum, had a significant p-value for whole community level statistics for the effect of genotype across all genotypes on the beta diversity of the piglet gut microbiota. However, there were no significant (P > 0.05) pairwise comparisons between specific genotypes. This is likely due to a lack of statistical power. Due to antibiotic administration prior to collection of the day 15 swab, because of confirmed infection in this litter, this time point was not included in analysis based on the knowledge that antibiotics affect the gut microbial composition (Gao et al., 2018). This decreased the already small sample size.

For Litter 23, our evaluation of the effect of genotype on the bacterial community composition was inconclusive. The two groups (A-SCID and non-SCID) had unequal variance, and thus, not all necessary assumptions were met for the PERMANOVA test. Genotype was not found to have a significant effect on the beta diversity of the litters that received the 7-strain inoculum (Litters 33 and 34).

Statistical analysis of alpha diversity further confirmed the lack of effect of genotype on the piglet microbiota, as assessed by beta diversity analysis (Supplementary Figure 2). The young age of the piglets used in the study could contribute to this indifference. We did observe a significant difference in alpha diversity of piglet microbiota due to piglet age for all three treatment groups (Supplementary Figure 3). Prior research has shown that there is a decrease in alpha diversity of the gut microbiota for SCID mice compared to non-SCID mice (Zheng et al., 2019). It is important to note that these SCID mice had a different mutation leading to their SCID phenotype than the pigs used in our study. These mice were nonobese diabetic (NOD)/SCID mice which have low, but detectable, natural killer cell activity. This decrease in alpha diversity was found over a three-month period, while our study only considered the pre-weaning timeframe. There are known important functional differences between mice and pigs so it is unclear if the same outcome would be seen with SCID pigs at a later timepoint (Dawson et al., 2013; Meurens et al., 2012).

The lack of SCID genotype effect on the piglet gut microbiota that we observed could be due to the short timeframe in which fecal swabs were collected. We collected intestinal tissue at day one and day seven from some piglets in litter 33 and 34 that were not included in the microbiome analysis presented in this manuscript. These samples included tissues from four piglets at day one and three piglets at day seven, across all three genotypes. The tissues were prepared for histological analysis and there were no notable differences in the samples between the two timepoints as well as the different genotypes (data not shown). Lack of an effect by genotype was also seen in aspects of a 7-day study performed by Rajao and colleagues (2016) using this line of SCID pigs which analyzed the effect of influenza A on the immune system.

While they observed notable differences of immune system function in some respects, when compared to non-SCID pigs in the same environment, there were parameters measured that did not appear to be different between the SCID group and the non-SCID group, such as IL-2 levels in the lungs (Rajao et al., 2016). The similarities between the two genotypes could have been due to the short time frame they were reporting on, as we report here. It could also be due to the developmental stage of the innate immune system during this early period of neonatal development.

With the knowledge about the effect of the gut microbiota on the immune system, it could be hypothesized that non-SCID pigs, and SCID pigs raised in the same environment, would have similar immune development at early timepoints. Because the piglet bloodstream is not accessible to the maternal bloodstream as in humans, pigs are dependent on antibodies acquired at birth, otherwise they are highly susceptible to infection until they can create their own antibodies (Butler et al., 2009). The SCID and non-SCID piglets are raised and fed following the same protocol. Similar immune development in this time period could be due to the only antibodies they are acquiring is through the irradiated bovine colostrum they are fed.

This could also be due to the lack of a complete gut microbiota to aid in the development of the host adaptive immune system that non-SCID pigs possess, unlike SCID pigs (Hill et al., 2012; Honda & Littman, 2016). Further studies performed for a longer duration of time are needed to determine if eventually there is an effect of SCID genotype on the gut microbiota of pigs.

Regardless of the limitations present for these trials, the lack of a significant effect of SCID genotype on the piglet gut microbiota in our system is an important observation.

### 4.2 Piglet age has a significant effect on the SCID piglet gut microbiota

On a whole community scale, there were significant differences in bacterial microbiota composition between sampling timepoints, regardless of genotype. Litter 17 (2-strain) pairwise comparisons revealed a significant difference in day 2 samples compared to day 7-8, which is reflected in Figure 3A. More specifically, the relative abundance of uninoculated bacteria present at day 7-8 is much higher than the previous two timepoints as seen in Figure 4.

Litter 23 (6-strain inoculum) did not have any significant differences in microbiota in pairwise comparisons, after adjustments, due to lack of statistical power from the available samples (n=4), seen in Table 2. There was a higher relative abundance of bacteria, that were not inoculated, at day 14-15 when compared to the previous three timepoints. Litter 23 had a high relative abundance of other bacteria present at day 14-15 compared to the inoculated bacteria, this is also the timepoint that antibiotics were necessary for litter 17. The high relative abundance of other bacteria at the day 14-15 timepoint could be an indication of infection in the piglets.

Analysis for Litter 33 and 34 (7-strain inoculum), revealed the most consistency across the day 4, day 7-8, and day 14-15 timepoints, leading to the conclusion that the 7-strain is the most stable overtime in these piglets. When comparing relative abundance of the bacterial strains in the 6-strain and 7-strain defined microbiota over time, *Bacteroides* had the highest relative abundance for both groups by day 7-8. The relative abundance of the phylum *Bacteroidota* is second, behind *Firmicutes*, in the composition of gut microbiota in piglets (Wang et al., 2022), consistent with the high relative abundance we observed in our samples. The function of *Bacteroides* in the piglet gut is to degrade polysaccharides derived from ingested milk (Guevarra et al., 2019; Ye et al., 2021) and its relative abundance increases during the pre-weaning timeframe (Fukuda et al., 2021). This means *Bacteroides* can thrive in the gut microbiome of pre-weaned pigs due to milk intake. Inclusion of *Bacteroides* in the 6-strain and 7-strain defined microbiota, appears to have allowed the SCID piglet gut microbial composition to follow this trend.

Beta diversity statistics for litter 33 and 34 (Figure 3C) showed a significant difference between day 2 and day 14-15, as well as between day 4 and day 14-15. Alpha diversity (Supplementary Figure 2C) for observed species, richness, and diversity present a statistically significant difference between both pairwise comparisons. The only exception was for the alpha diversity statistic of evenness as there was not a significant difference in evenness between day 4 and day 14-15. Overall, it appears an increase in the complexity and concentration of the inoculum, offers greater stability as piglets age. This was observed in previous research analyzing succession of the gut microbial community of pigs (Dong et al., 2023; Luo et al., 2022). Dong and colleagues (2023) determined the composition of the gut microbiota at an early timepoint varies between animals; however, as piglets get older, the composition of the gut microbiota was more similar across the group. This finding also supports the data Luo and colleagues (2022) reported, that there is a “core” microbiota, bacterial species consistently present, found in the gut of healthy piglets. Our findings support this published data which is a positive outcome for our results.

### 4.3 Implications

Pre-weaning was the time frame of focus for this study due to our concern of piglet loss from *C. difficile* or other rapid systemic infections. We did not see *C. difficile* infections while using any of the defined microbiota consortia tested, which suggests the inoculated bacteria is not allowing for the proliferation of *C. difficile*, if it was present. Additionally, the defined microbiotas did not cause harm to the piglets as none of the inoculated bacteria was cultured from the bloodstream of any piglets. Based on these results, we believe this is a promising initial step for the development of a defined microbiota, that can support the SCID pig gastrointestinal tract. The 7-strain inoculum is the most complex defined microbiota tested and it appears to offer increased stability to the piglet gut microbiota over the first two weeks of life, when compared to the 2-strain and 6-strain inoculum.

### 4.4 Limitations

More studies are necessary to examine the addition of a defined microbiota in SCID pigs as these biomedical models are important to gain insight in human processes. When working with a large immunodeficient biomedical animal model, there are many limitations present. The costs associated with creating and raising these litters is very high due to substantial labor input and lack of established husbandry protocols. Another challenge with SCID pigs is small litter sizes. The number of offspring in a SCID pig litter can be significantly smaller than a sow on a production farm which means there are less animals available for studies. This is, in part, the reason this study has a low sample size.

Creating a biocontainment room that can house SCID pigs safely is costly, and so is the effort required to keep these environments clean. Even though much effort was put towards keeping microorganisms out of the biocontainment facility during the piglet trials, it is difficult to keep the environment completely free of microorganisms that could be pathogenic to the SCID pigs. As seen in Figure 4, a portion of the overall microbiota present were bacteria that were not inoculated into the piglets. This leads to the conclusion that husbandry methods need to be improved to keep unwanted bacteria out of the piglets including environmental decontamination throughout the research study. There is also potential that the contamination present is from the sow and not from the environment, however, more research is needed in this area. Preventing the infiltration of pathogenic bacteria is difficult with a large animal model as care staff must enter the SCID pig biocontainment space to care for them. This makes it more challenging to keep an SPF environment when compared to smaller models, like SCID mice. The size of SCID mice allow them to live in small environments and they can be handled in a biosafety cabinet limiting potential for pathogen exposure, which is not possible for a large animal model.

Another challenge we encountered is the short sampling period. While we were focused on the pre-weaning time frame, that only allows 14-21 days for sample collection. We intentioned to continue collecting data after weaning, as well as giving a third dose of microbiota inoculum post weaning; however, medical intervention was necessary, hindering the use of data past the weaning timepoint. It is expected that the SCID piglet gut microbiota composition would shift after weaning, which is something to be analyzed in the future. Inoculating SCID piglets with a third or additional doses of microbiota after weaning or after antibiotic use would also be important to determine how different feeds or antibiotic use affects the inoculated bacteria.

Additionally, we used 16S rRNA gene amplicon sequencing. It is well known that amplicon sequencing is only semi-quantitative. The ideal method for greater accuracy when quantifying the amount of inoculated and uninoculated bacteria in the gut of the piglets is qPCR, of the extracted DNA, from the collected fecal swabs. By using amplicon sequencing, we were only able to present relative abundances of bacterial species (and not absolute abundances). We used DNA for sequencing, because of this we cannot prove that the obtained sequences were indeed from living organisms. Lastly, these were low biomass samples. To control for this in the sequencing process, we included many controls throughout the process. This included controls at the end of each DNA extraction kit using PCR grade water, blank fecal swabs, and PCR grade water at the end of the plate submitted for sequencing. These controls allowed us to account for possible contamination throughout the processing of samples. Then, for the analysis of samples, the decontam package, discussed in the materials and methods section of this manuscript, was used to detect contaminating DNA, and remove it. These methods were utilized to make our data the best quality they could be, despite their low biomass.

### 4.5 Future studies

In the future, a further increase in concentration of the inoculum or an addition of more bacterial strains to the inoculum could be implemented. These changes could lead to a decrease in the percent of uninoculated bacteria present in the SCID pigs. A more complex, but still defined, microbiota for use in gnotobiotic mice has been created (Cheng et al., 2022) which would be important to explore to improve the SCID pig model. The outcome of this mouse study was a gut microbiome that saw an increase in stability when challenged and displayed resistance against pathogenic *E. coli* (Cheng et al., 2022). As previously mentioned, a third (or additional) inoculation time point could be utilized post weaning or post antibiotics as both events lead to a change in the pig gut microbiota (Gao et al., 2018; Yu et al., 2021). With the changes to the SCID piglet husbandry protocol discussed in this section, the SCID pig could become a more reproducible and accessible model.

## 5 Conclusions

This study looked at the effect of SCID genotype and piglet age on the defined microbiota in the gut of piglets. The results revealed there was no significant effect due to SCID genotype while piglet age had a significant effect on the gut microbiota of the piglets over time. More research is necessary to examine a defined microbiota in the husbandry of SCID piglets over a longer length of time, however, it appears the consortia we created were safe to administer to SCID piglets and may be beneficial in preventing *C. difficile*. The 7-strain defined microbiota, which had not only the greatest number of bacteria strains but also the highest concentration of bacteria. This appeared to offer more stability to the gut microbiota composition over time making it the most ideal candidate to give to SCID piglets. It will be added to our SCID pig husbandry protocol allowing for rearing of SCID piglets to be more standardized, enabling reproducibility by other research groups.

## Acknowledgements

Katherine Widmer performed the collection of data and wrote the manuscript. Faith Rahic- Seggerman performed statistical analysis, wrote portions of the manuscript, and prepared the defined microbiota for each litter. Ahlea Forster was the project manager for the animal trials. Amanda Ahrens-Kress and Mary Sauer were responsible for the veterinary care the animals received. Amanda Ahrens-Kress wrote the section describing cesarean-section of the piglets. Shankumar Mooyottu and Brett Sponseller were the project leaders responsible for the *Peptacetobacter spp.* Akhil Vinithakumari trained FRS on the methods for preparation of the defined microbiota, and Akhil Vinithakumari and Aaron Dunkerson-Kurzhumov provided early aliquots of the strain *Peptacetobacter spp.* Matti Kiupel performed histological analysis on piglet intestinal samples we provided. Stephan Schmitz-Esser was responsible for conceptualization, supervision as the major professor of FRS, and resources. Christopher Tuggle was responsible for conceptualization, funding acquisition, project administration, resources, and supervision as the major professor for KW. We thank Linda Saif and Gireesh Rajashekara from The Ohio State University for providing several bacteria strains, as seen in Huang et al., that were included in the defined microbiota. This research was supported by NIH Office of the Director, R24OD028748-04.

## Supplementary Figures and Tables

**Supplementary Figure 1.**
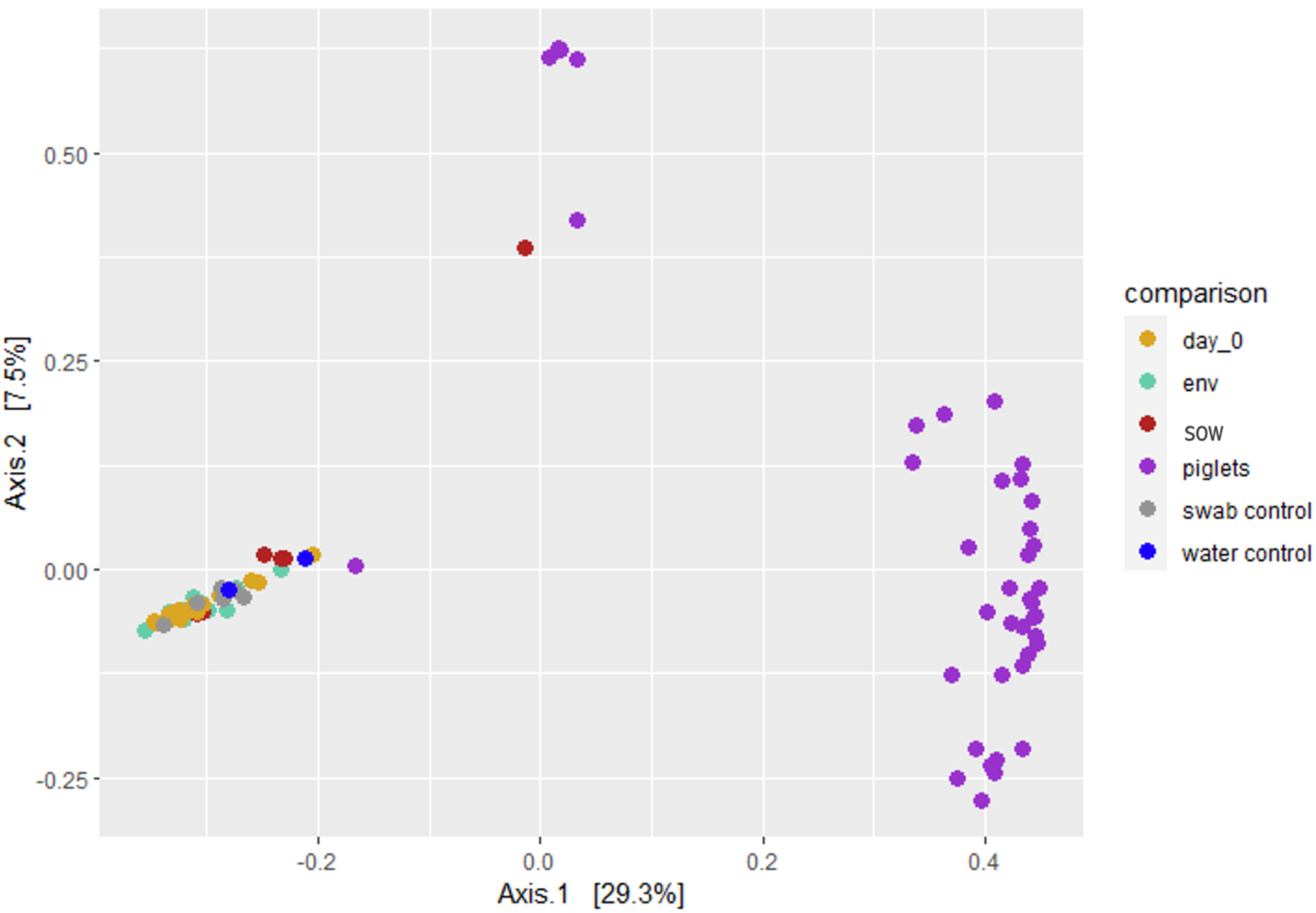
Beta diversity of non-day 0 swabs are significantly different from the day 0 and control swabs for litters 33 and 34 and controls, defending the decision to not include day 0 swabs in analysis. Due to the additional collection of multiple control samples from the cesarean section of Litter 33 and Litter 34 compared to the previous two trials involving Litter 17 and Litter 23, this PCoA plot was generated to further reinforce the decision to exclude the day 0 timepoint swabs. Distances between samples denote Bray-Curtis dissimilarity measures based on 16S rRNA gene amplicon sequencing. Swabs collected from every time point, other than day 0, were pooled together and labeled as “piglets” to determine the relationship between the day 0 swabs, control swabs and the rest of the time points. The “piglets” swabs are clearly clustered together, separate from the clustering of the rest of the swabs collected. This aids in the understanding that the swabs collected at day 0 are most like the control swabs, leading to the understanding that the Illumina MiSeq data from the day 0 samples is likely background noise and does not contain data that would need to be included in analysis.

**Supplementary Figure 2.**
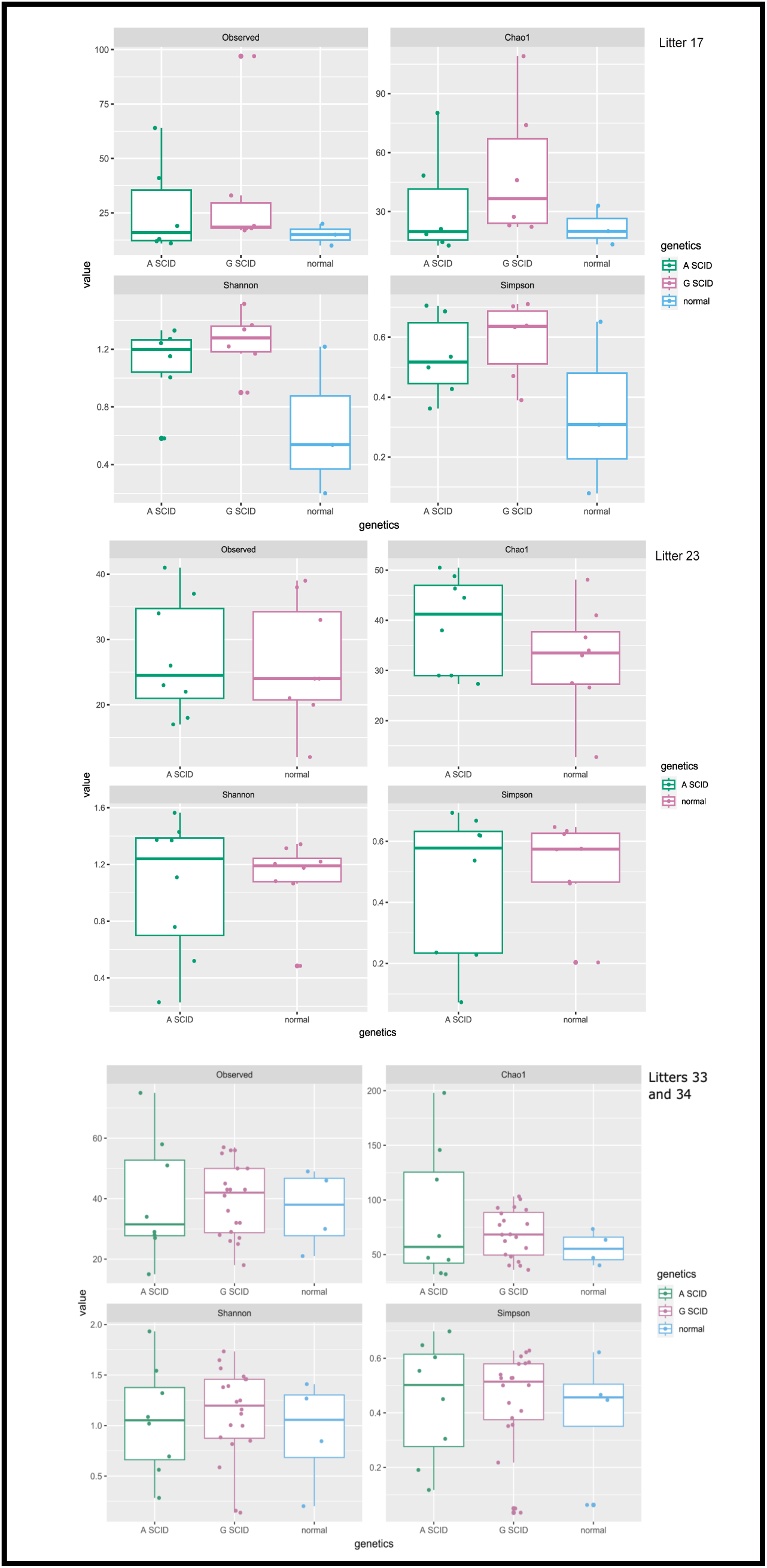
There was no significant effect of genotype on the alpha diversity of SCID piglet genotype on piglet gut microbiota, regardless of piglet age. This figure displays the number of observed species (Observed), species richness (Chao1), diversity (Shannon), and evenness (Simpson). **(A)** There were no significant differences (P>0.05) in alpha diversity measures between the three genotypes in litter 17. **(B)** There were no significant differences (P>0.05) in alpha diversity measures between the two genotypes in litter 23. **(C)** There were no significant differences (P>0.05) in alpha diversity measures between the three genotypes in litter 33 and 34.

**Supplementary Figure 3.**
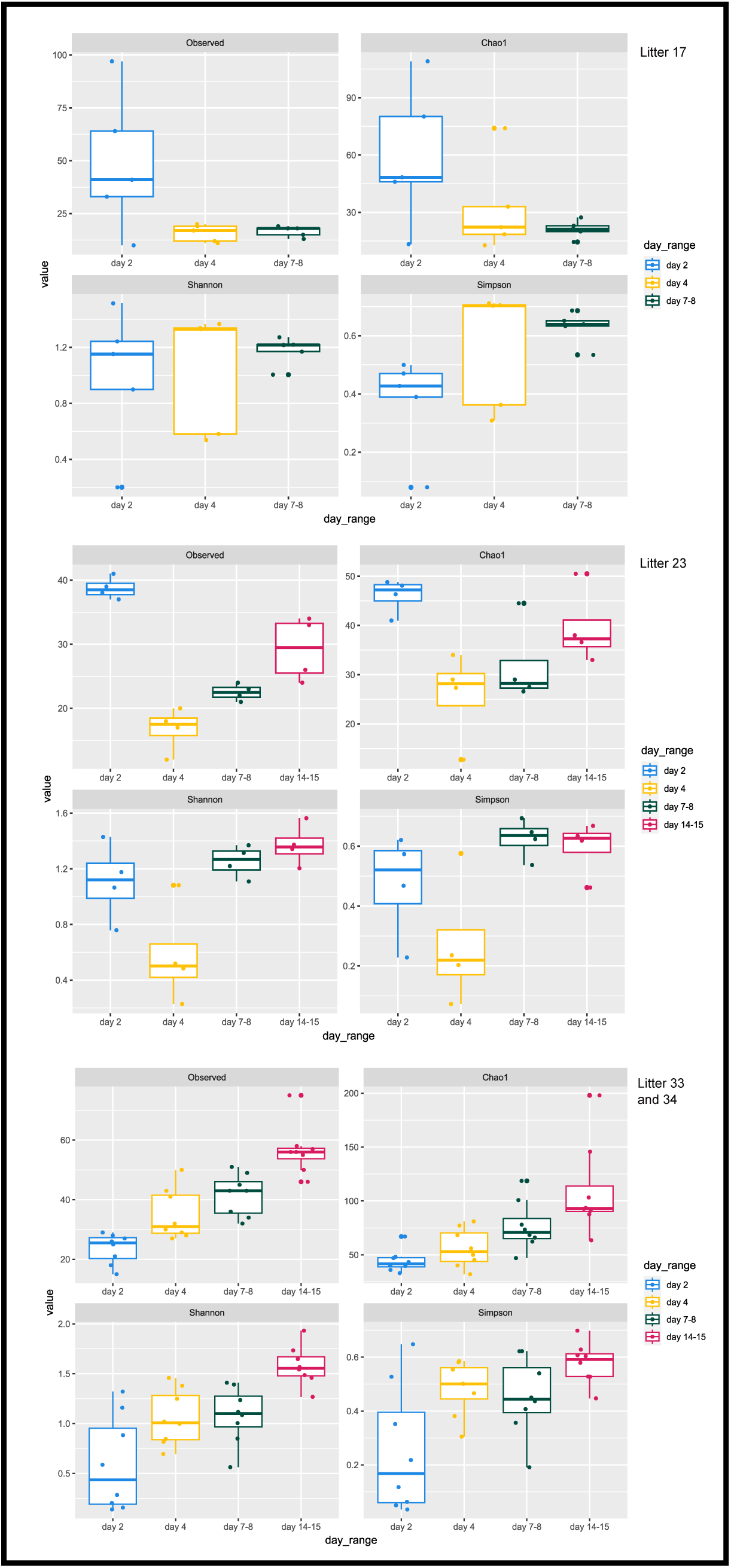
There was a significant difference in alpha diversity of piglet gut microbiota due to piglet age, regardless of genotype. The figure compares the number of observed species (Observed), species richness (Chao1), diversity (Shannon), and evenness (Simpson) between the sampling timepoints. **(A)** Alpha diversity estimators for litter 17 revealed a significant difference in evenness (P = 0.026) over time due to piglet age across the three timepoints. Pairwise comparisons for evenness revealed a significant difference (P=0.023) between day 2 and day 7-8 in piglet microbiota due to piglet age. This litter received antibiotics prior to collection of day 14-15 swabs so they were omitted from analysis. **(B)** Alpha diversity estimators for litter 23 revealed a significant difference in observed species (P = 0.002), species richness (P = 0.028), and diversity (P = 0.014) across the four timepoints due to piglet age. Pairwise comparisons show there was a significant difference in observed species between day 2 and day 4 (P=0.008), between day 2 and day 7 (P=0.007), and between day 4 and day 14 (P=0.019). There was also a significant difference in diversity between day 4 and day 14 (P=0.010). **(C)** Alpha diversity estimators for litters 33 and 34 revealed a significant difference in the number of observed species (P <0.001), species richness (P < 0.001), diversity (P = 0.001), and evenness (P = 0.017) due to piglet age across the four timepoints due to piglet age. Pairwise comparisons show there was a significant difference in observed species between day 2 and day 7-8 (P< 0.001), day 2 and day 14 (P<0.001), and day 4 and day 14 (P<0.001). There was a significant difference in species richness between day 2 and day 14 (P<0.001) and between day 4 and day 14 (P<0.001). There was a significant difference in diversity between day 2 and day 7-8 (P=0.029), day 2 and day 14 (P=0.001), and between day 4 and day 14 (P=0.036). Lastly, there was a significant difference in evenness between day 2 and day 7-8(P=0.048) and between day 2 and day 14 (P=0.014).

**Supplementary Table 1:**
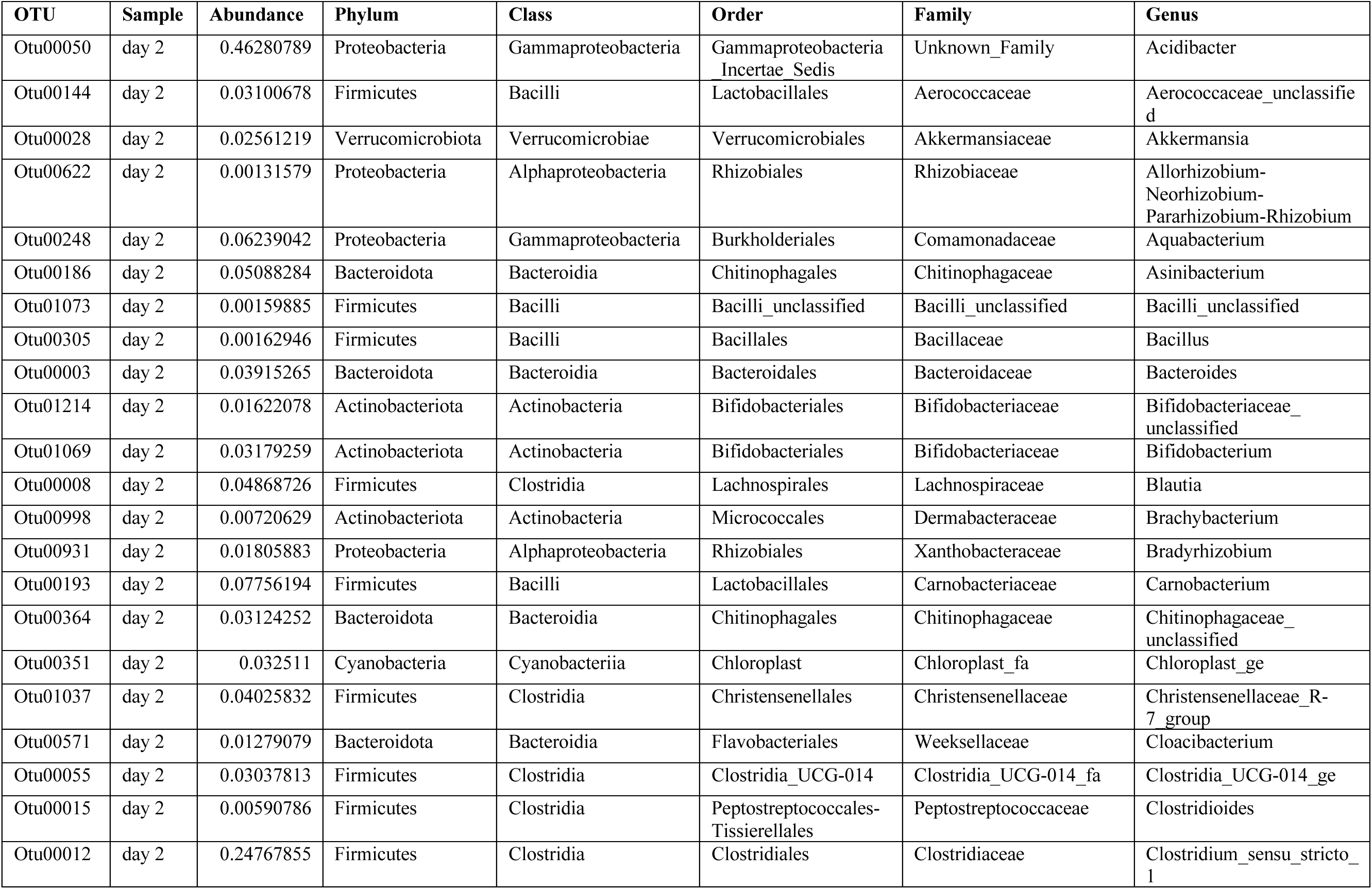
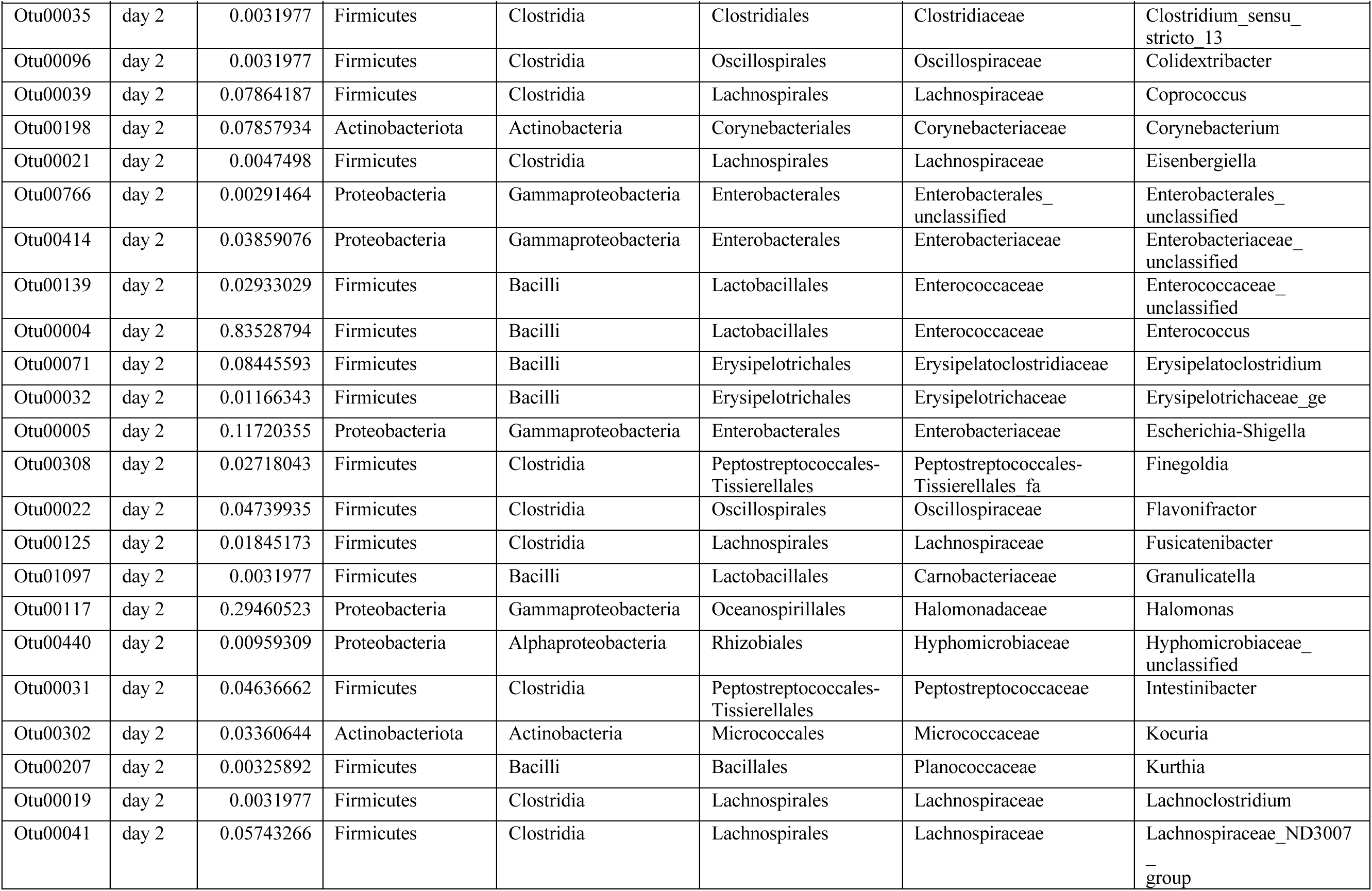
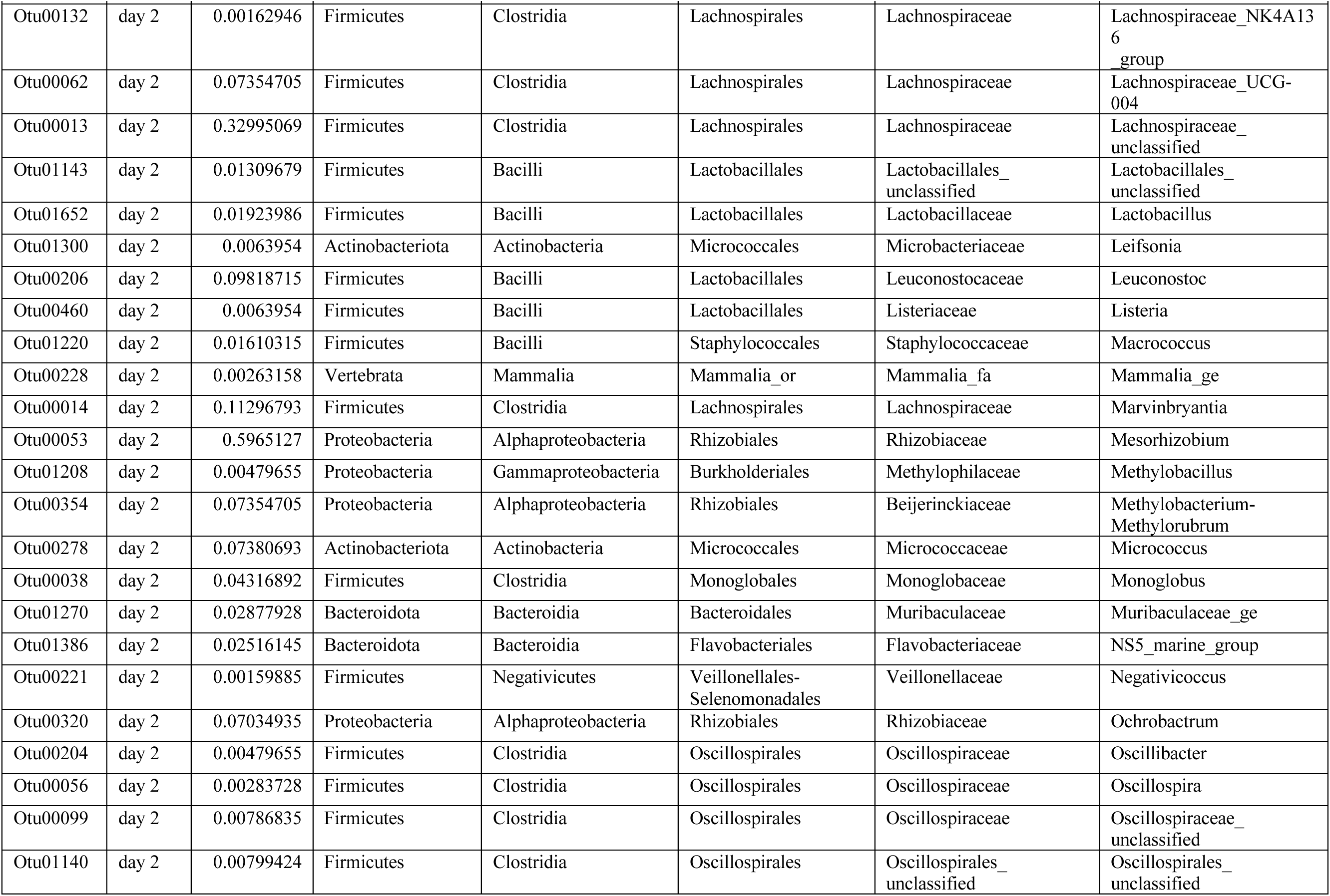
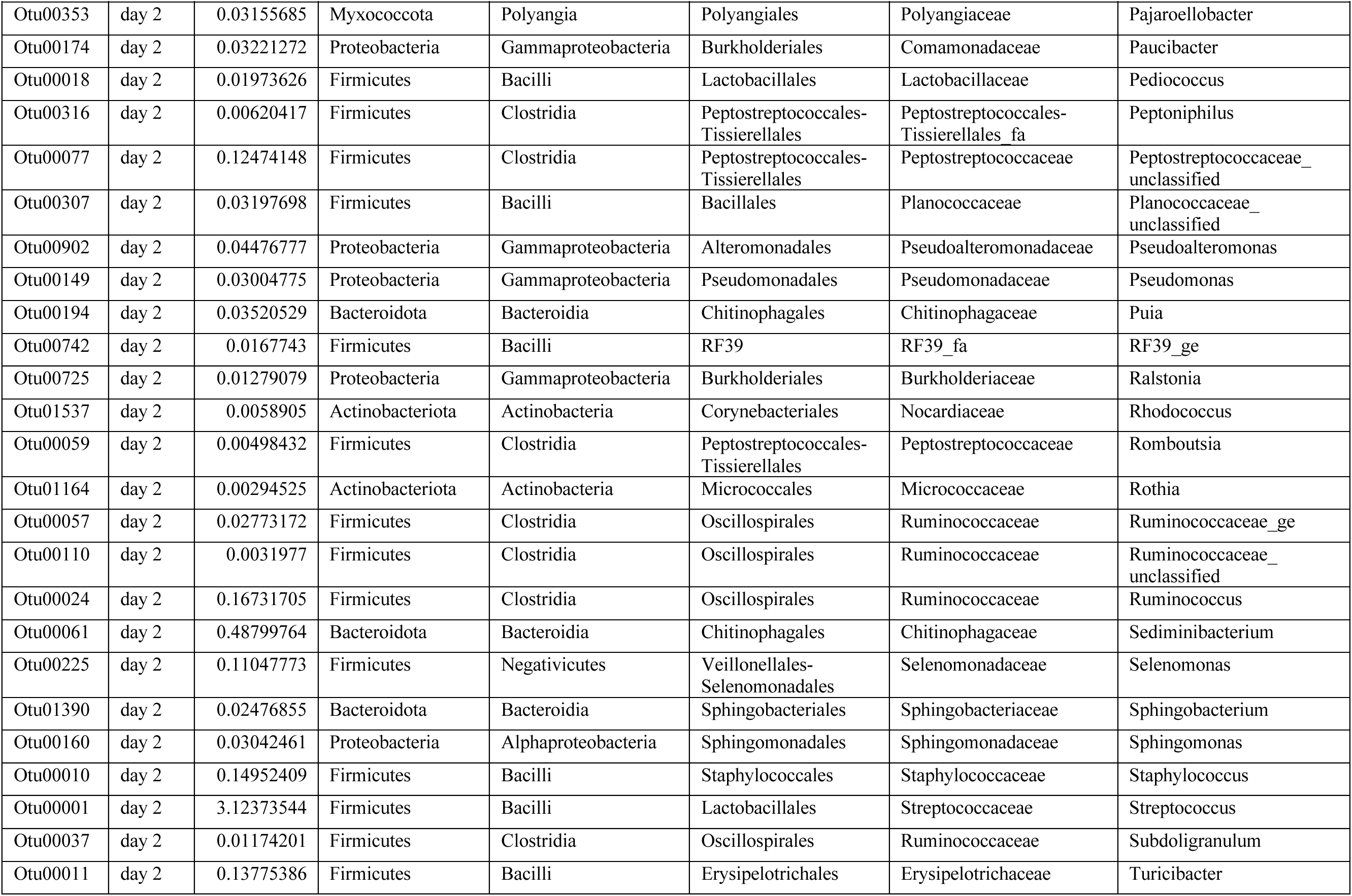
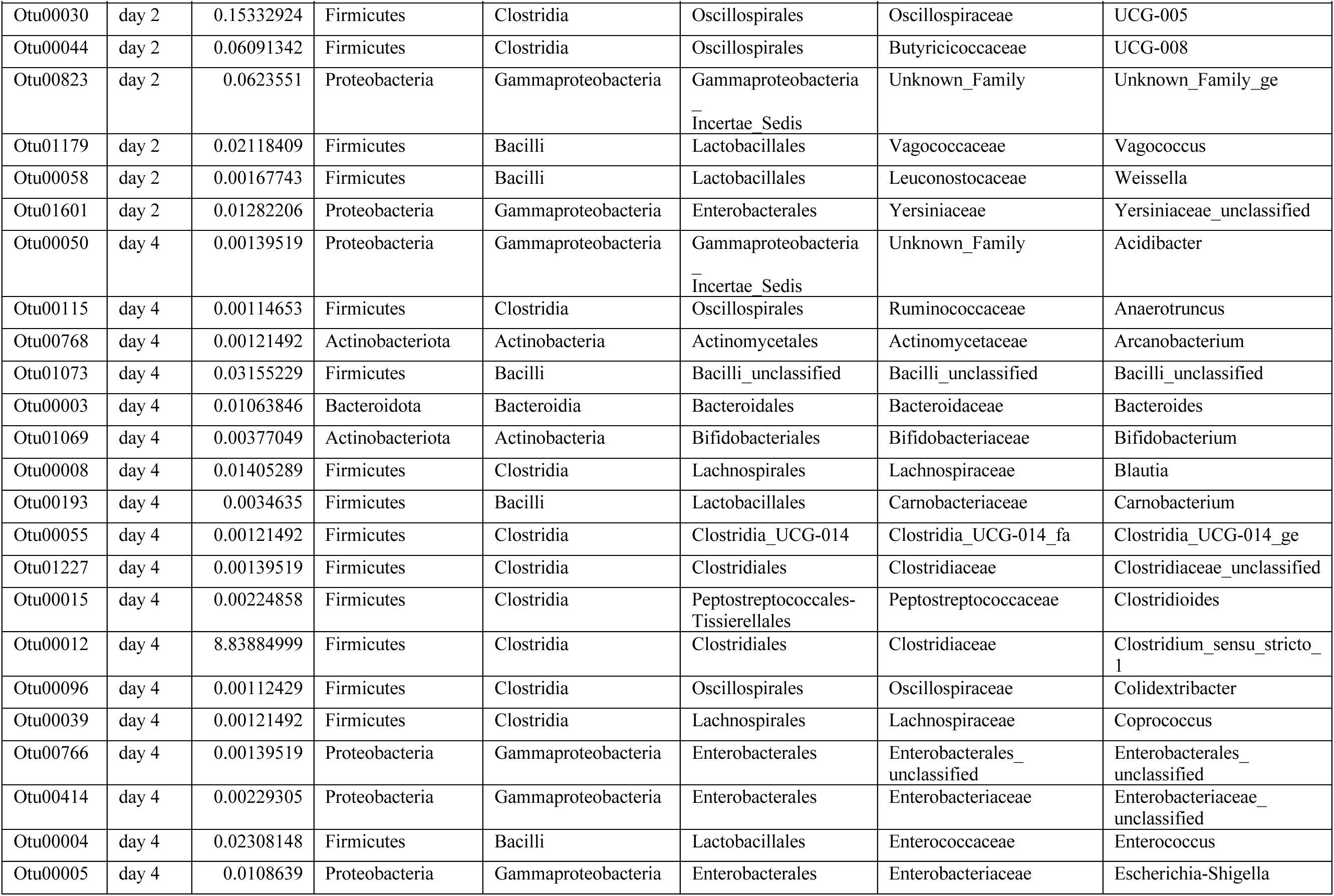
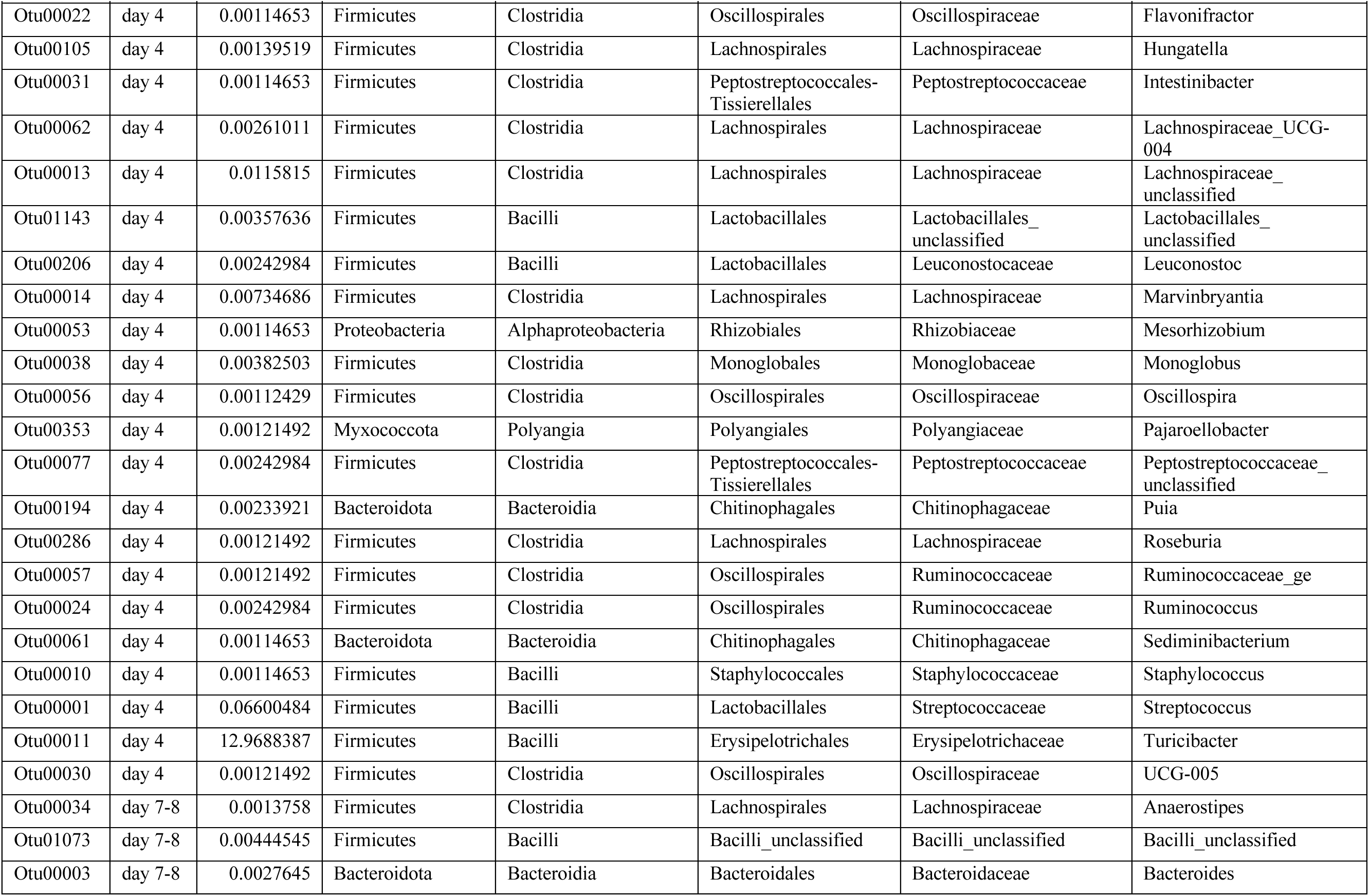
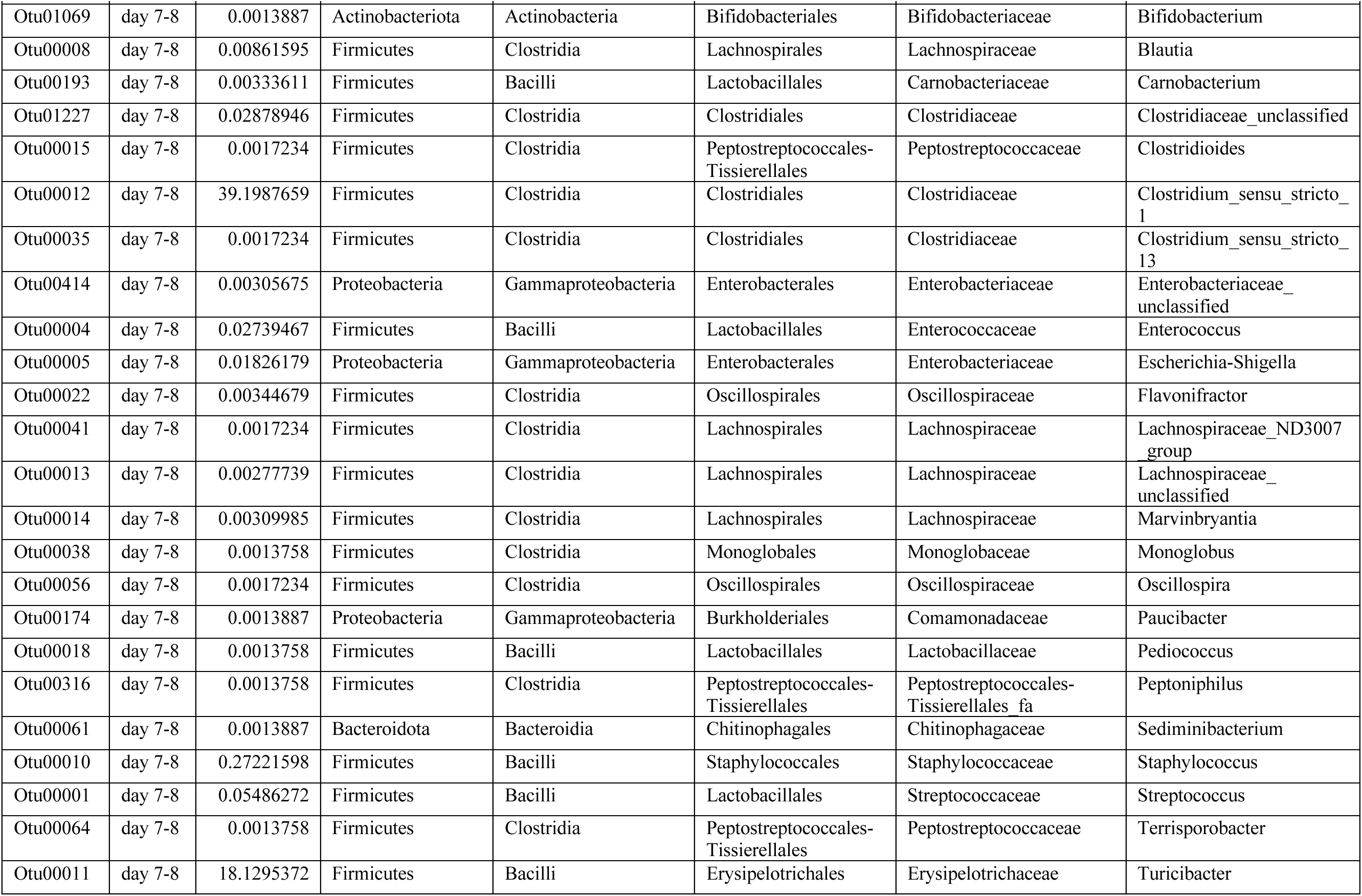
List of “Other” bacteria as seen in Figure 4 for litter 17.

**Supplementary Table 2:**
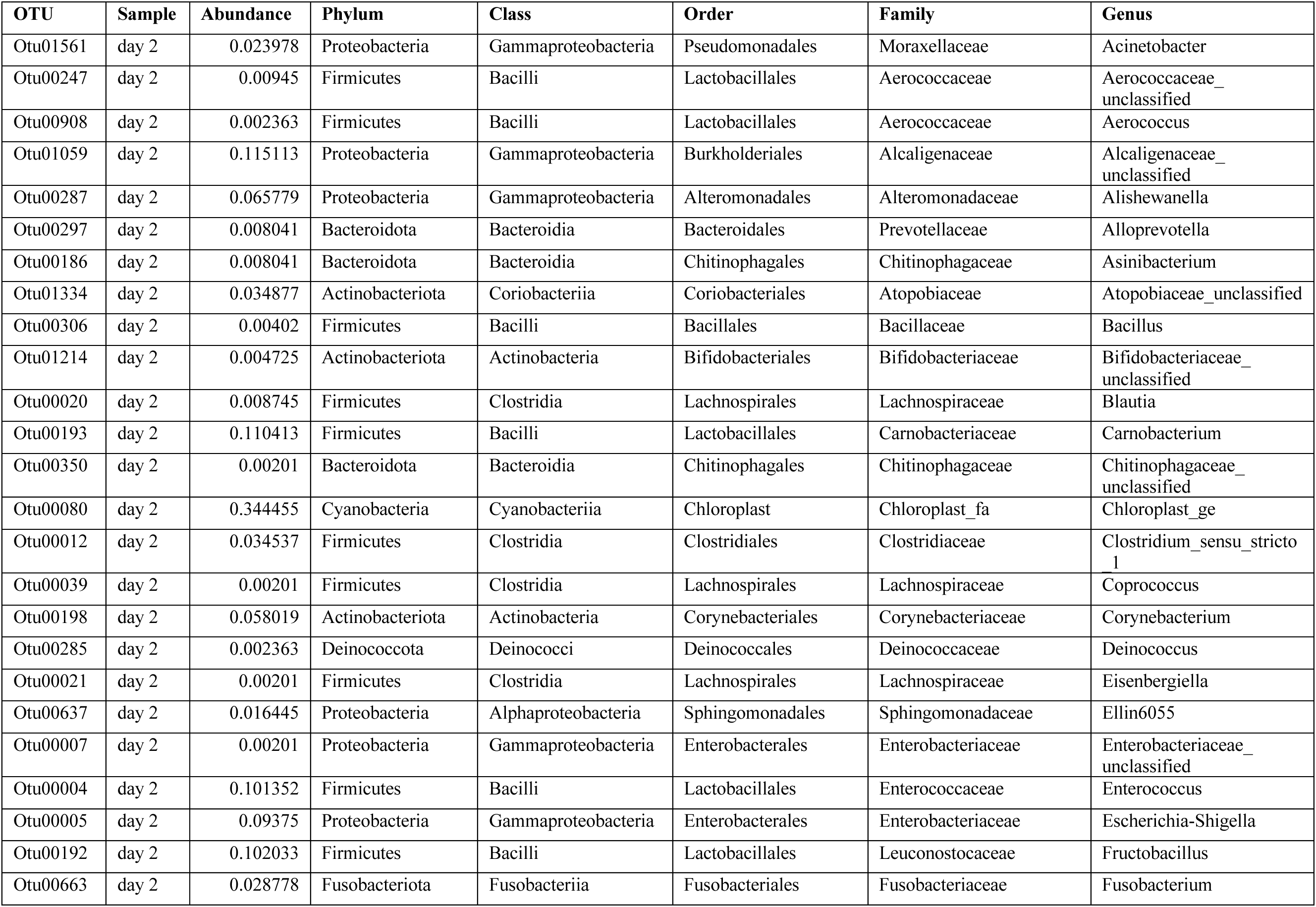
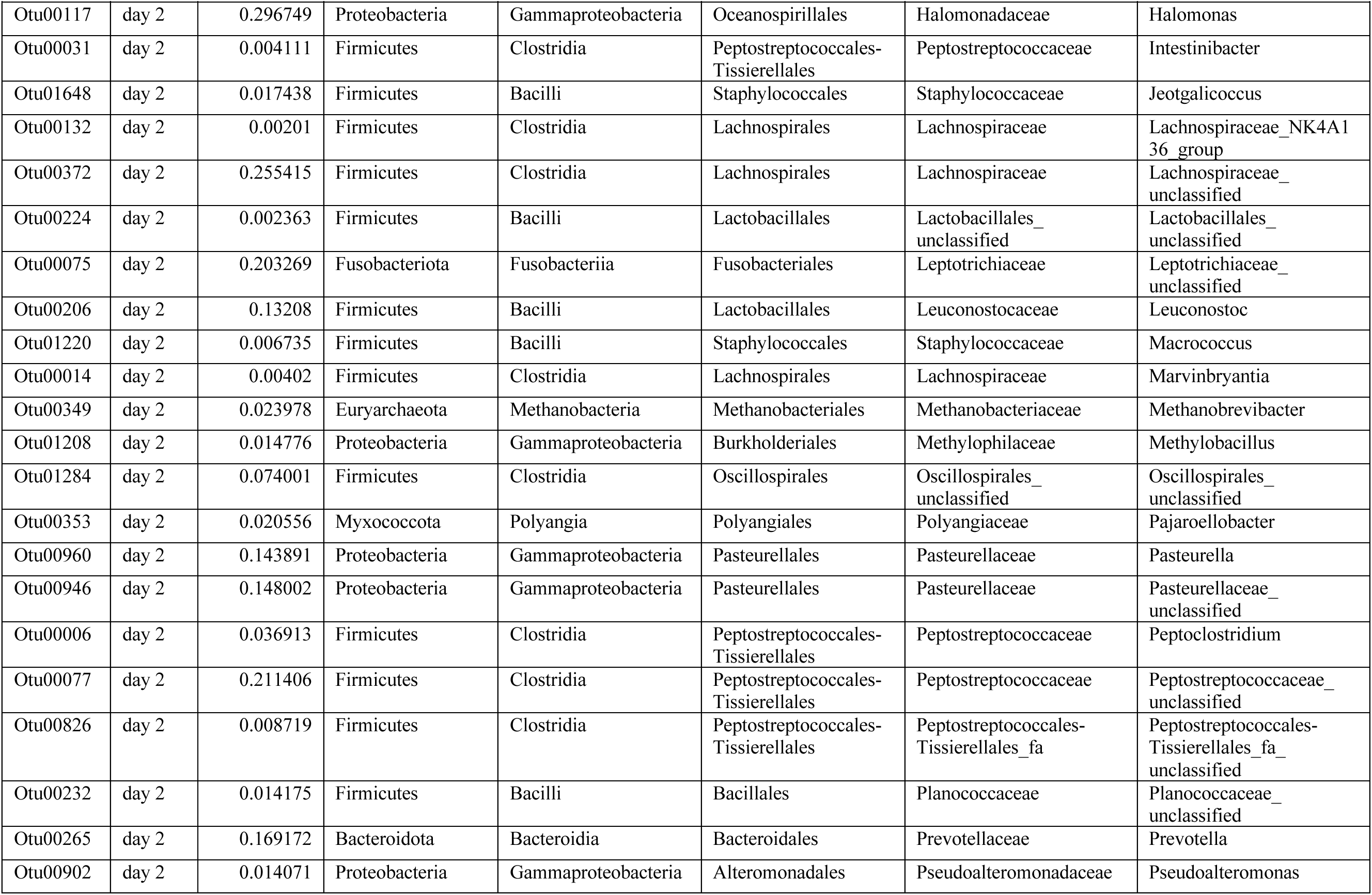
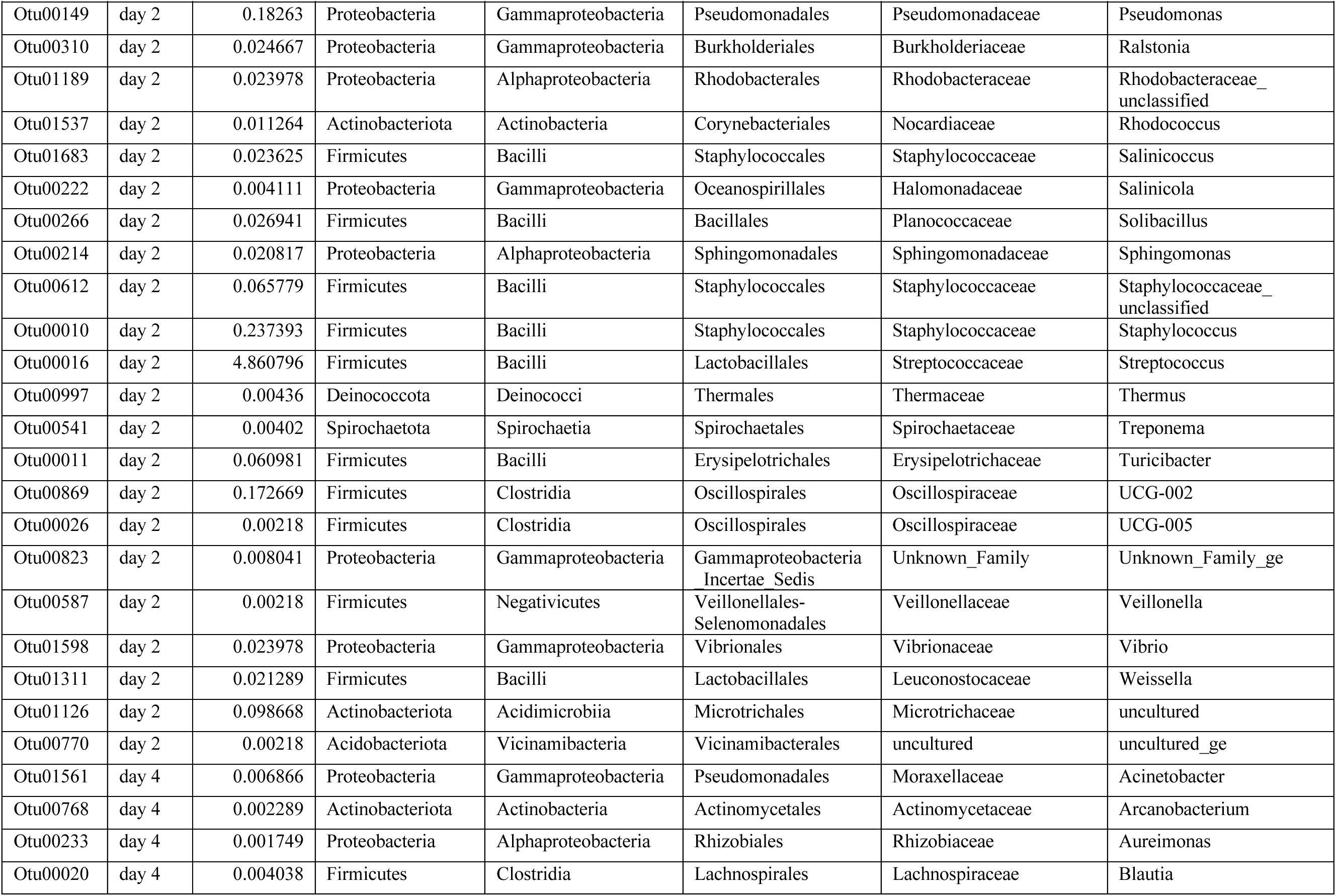
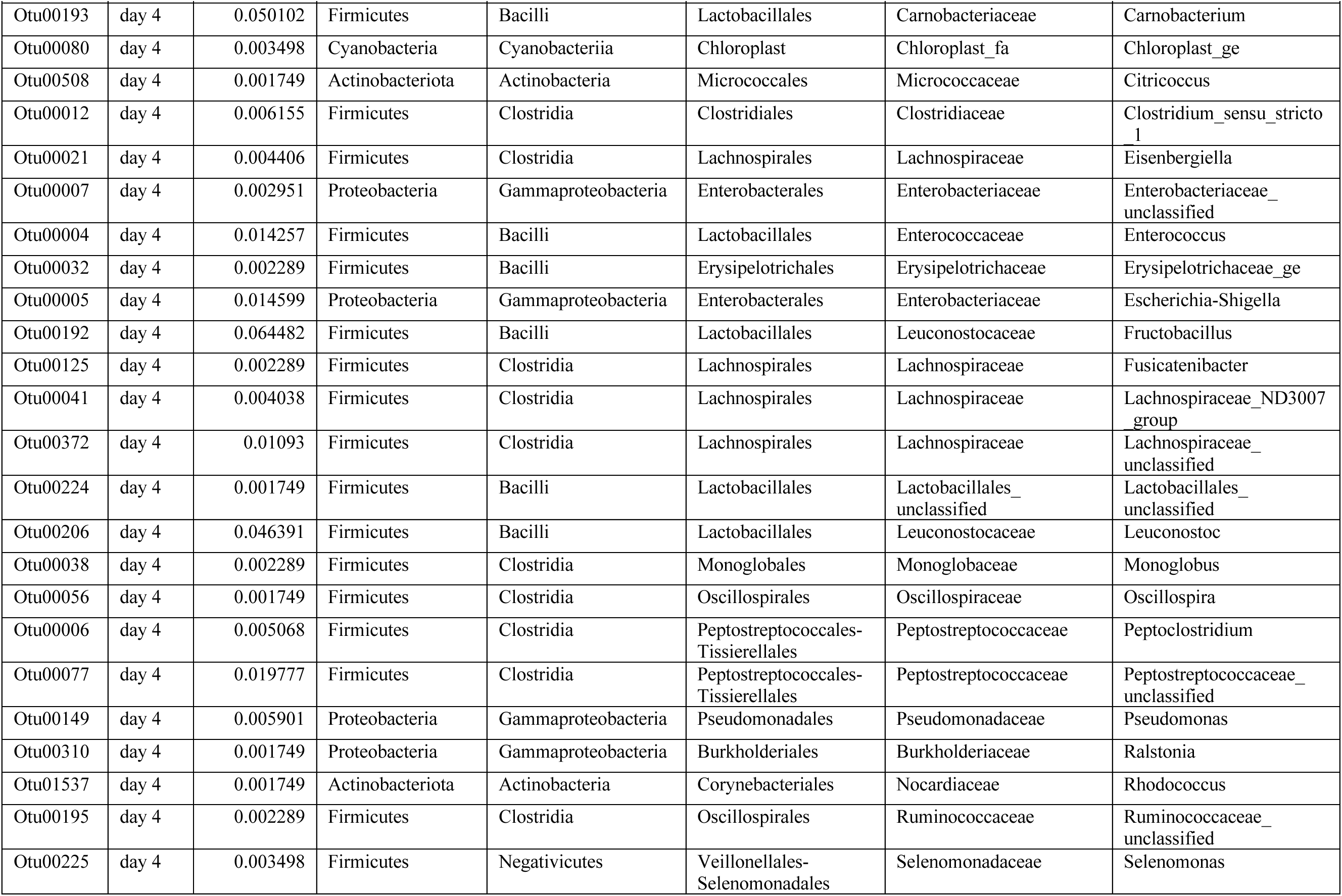
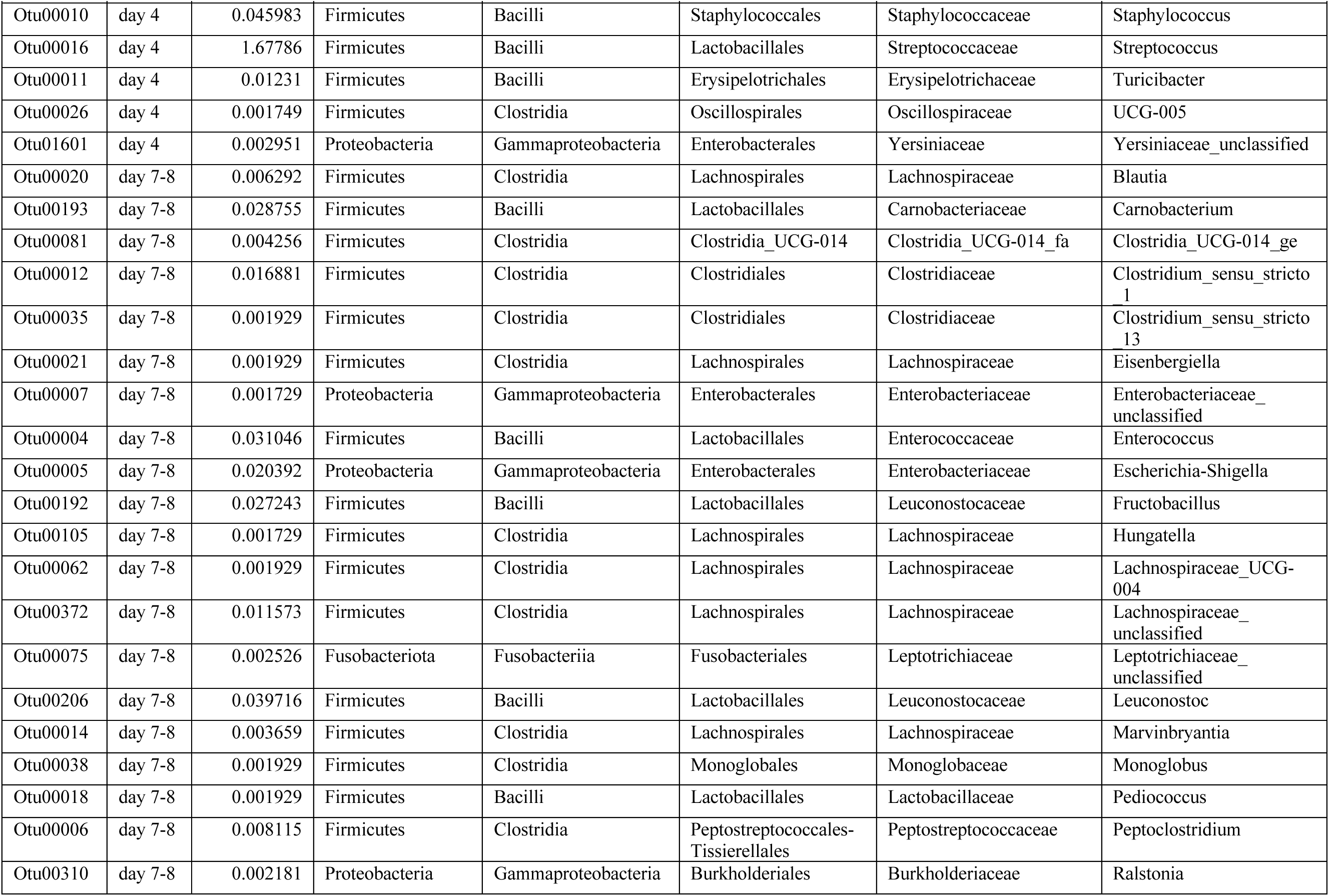
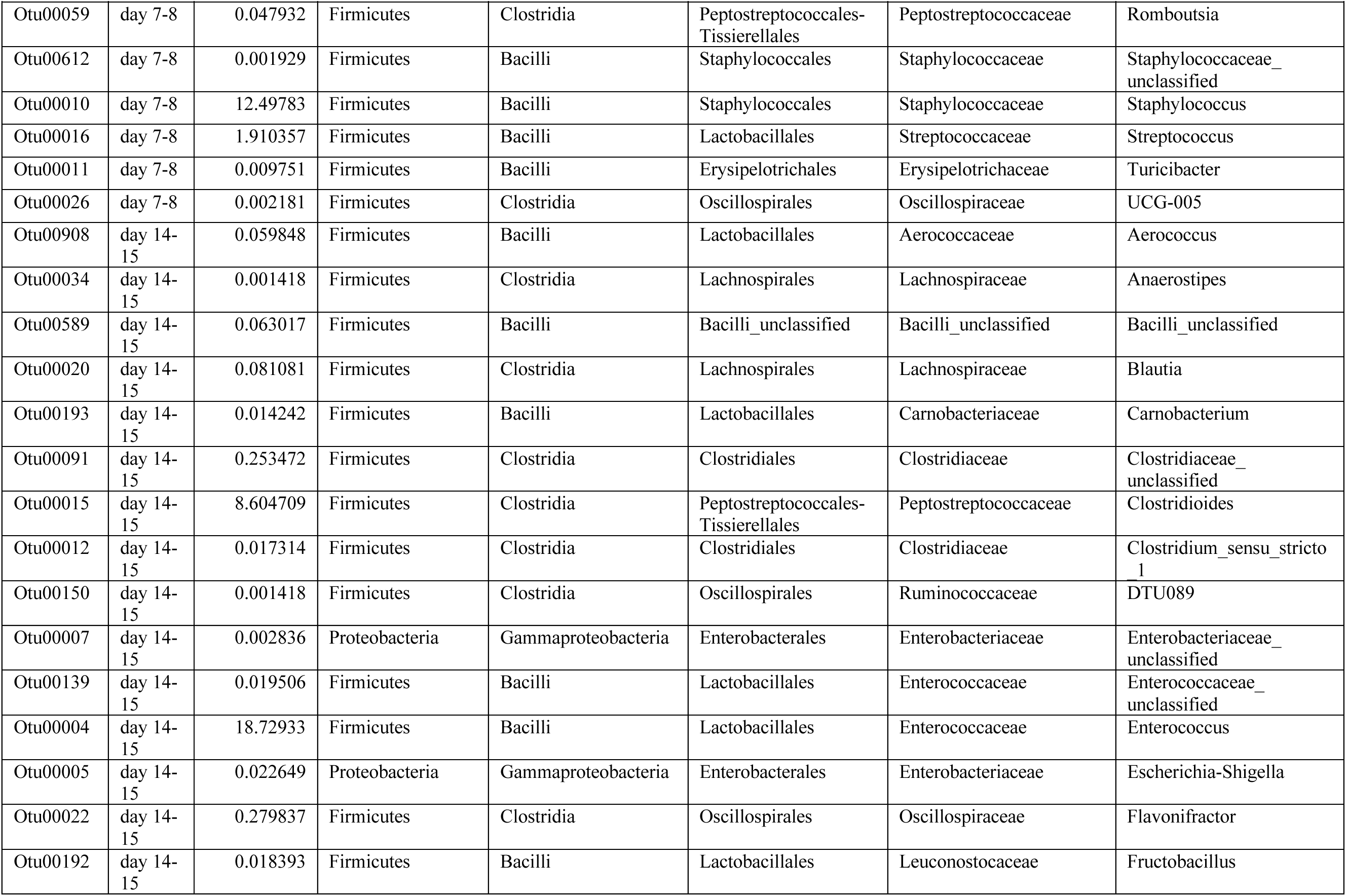
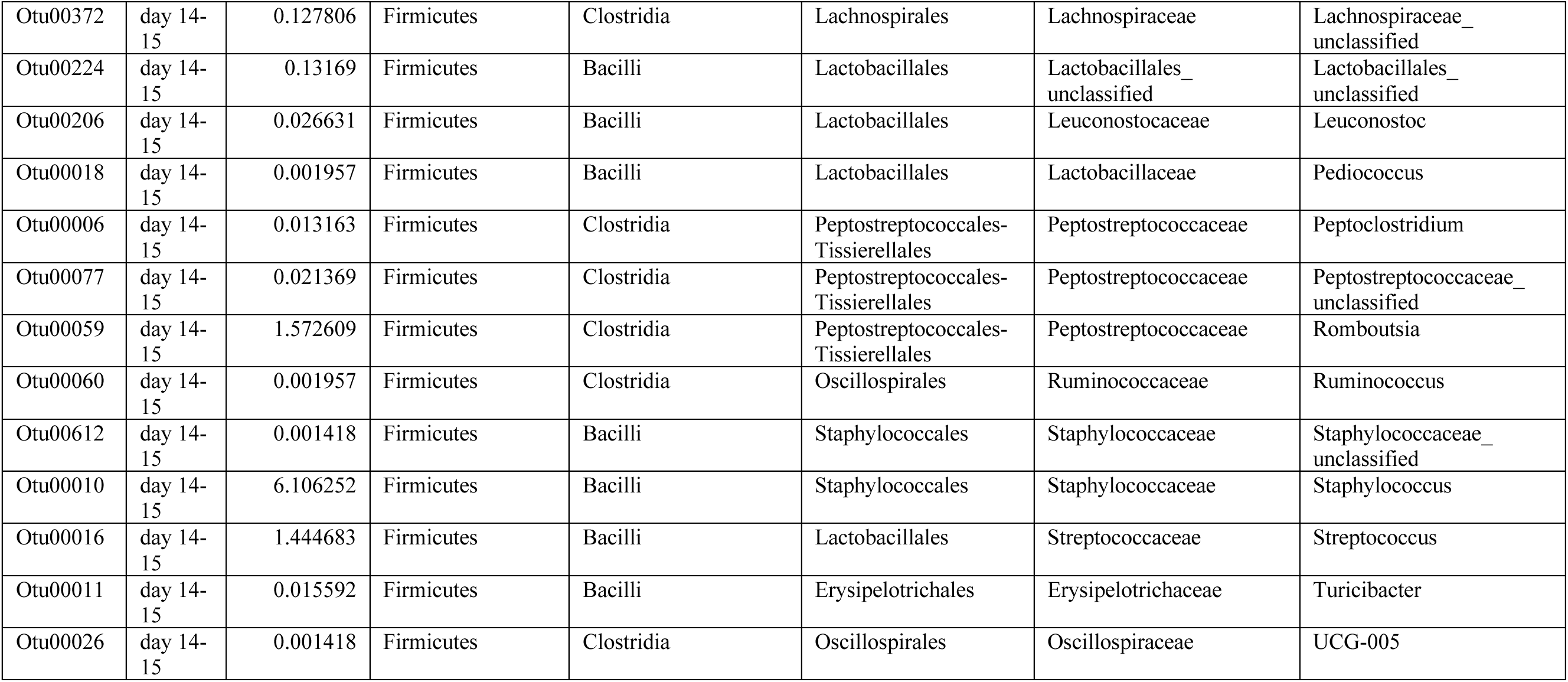
List of “Other” bacteria as seen in Figure 4 for litter 23.

**Supplementary Table 3:**
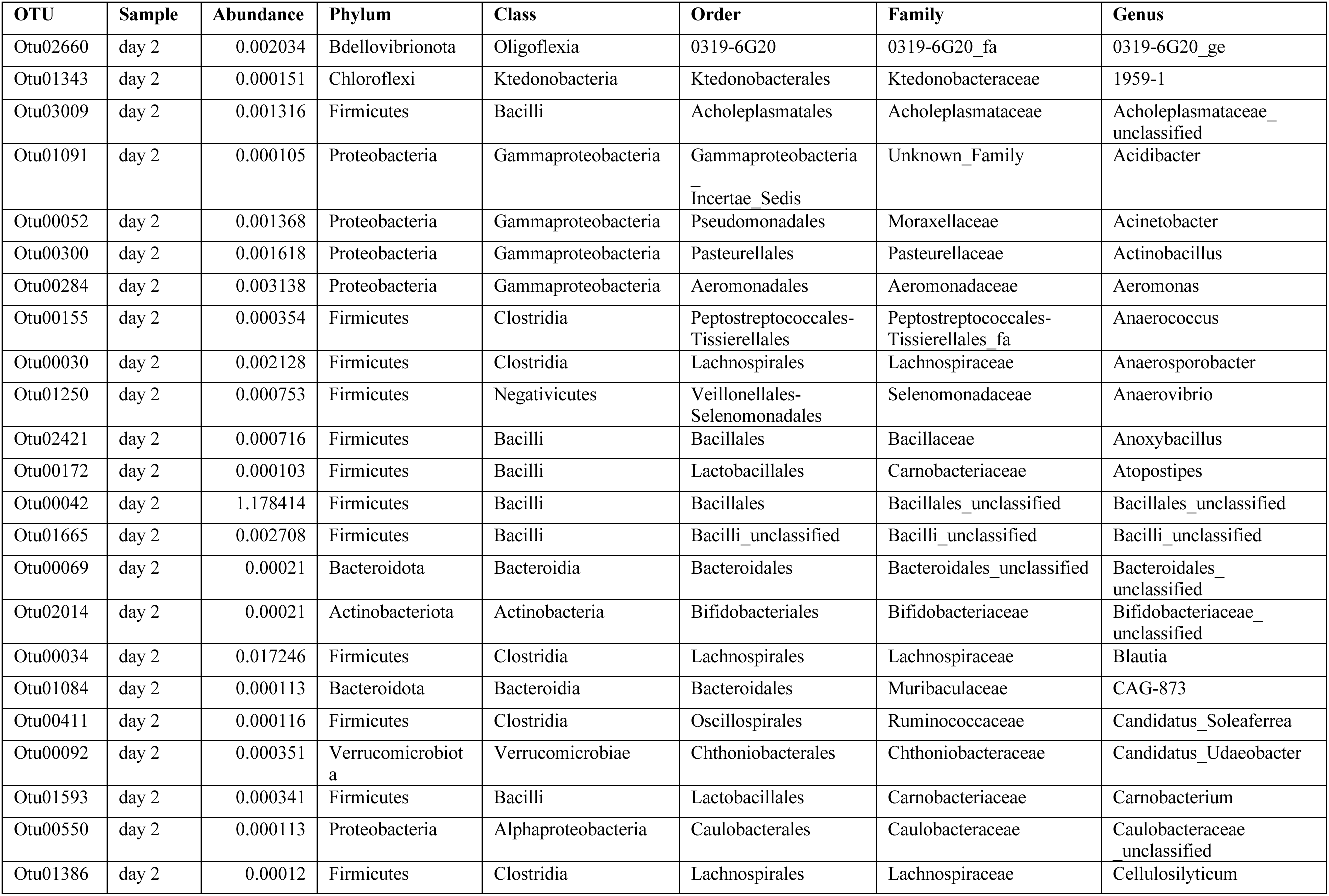
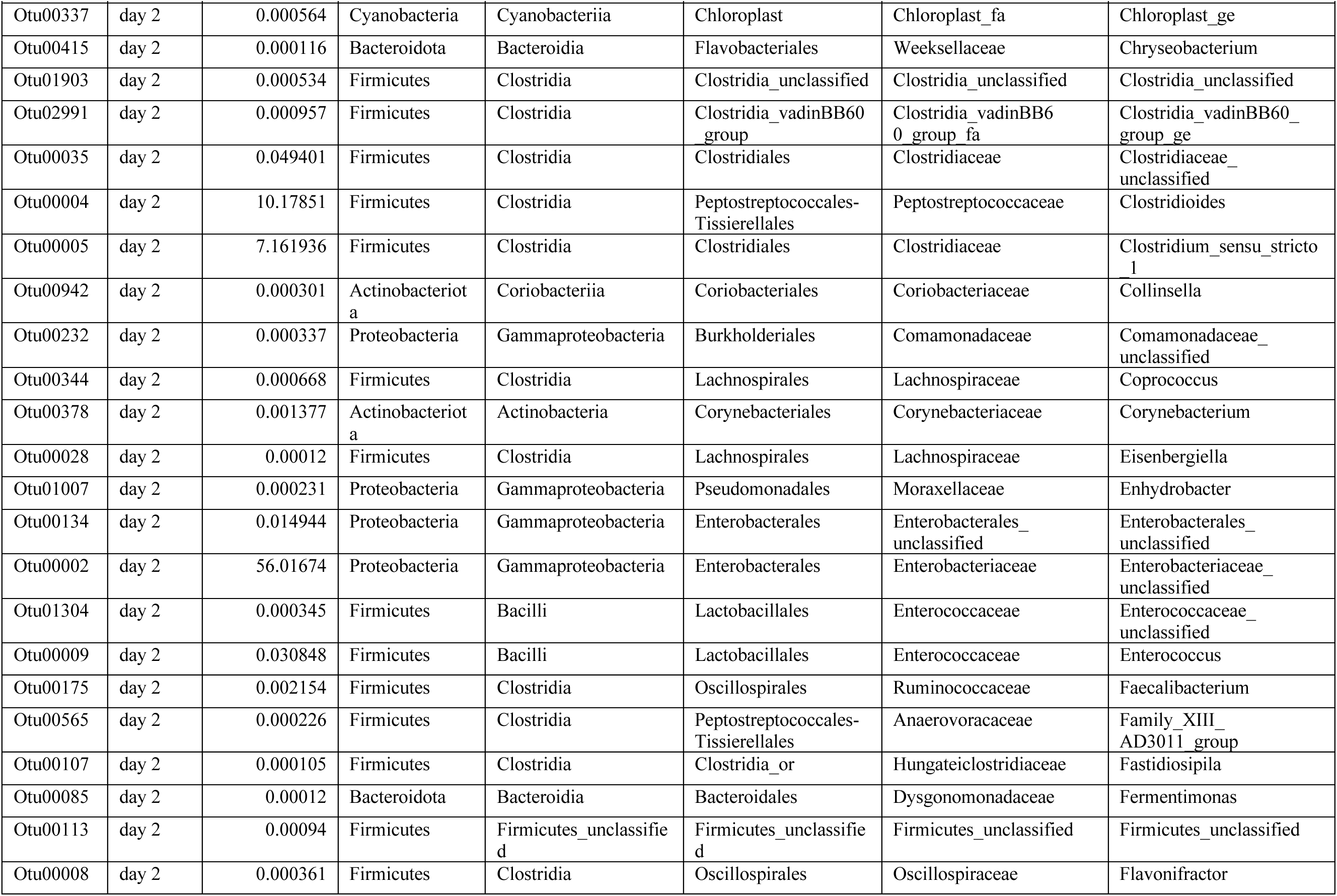
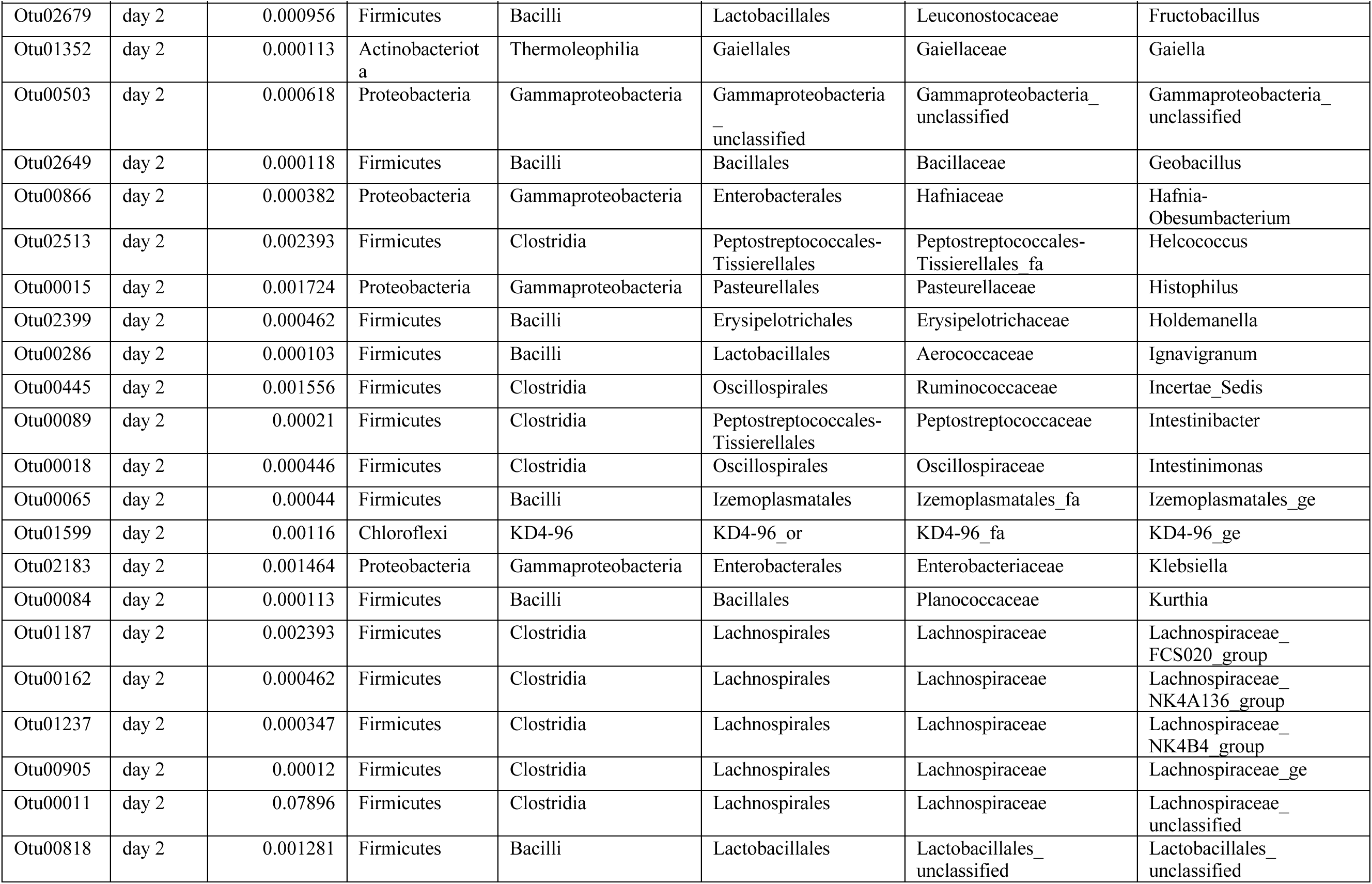
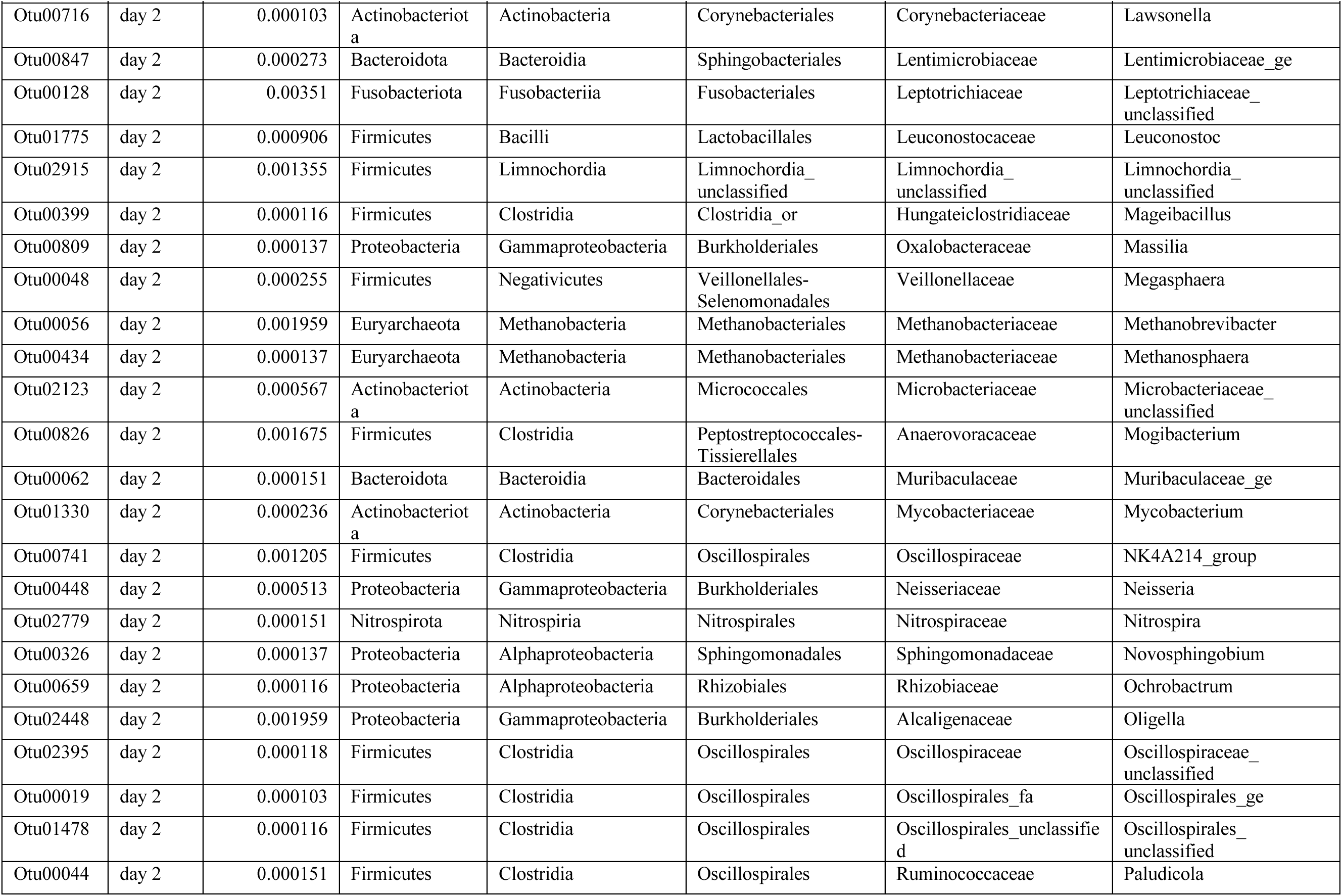
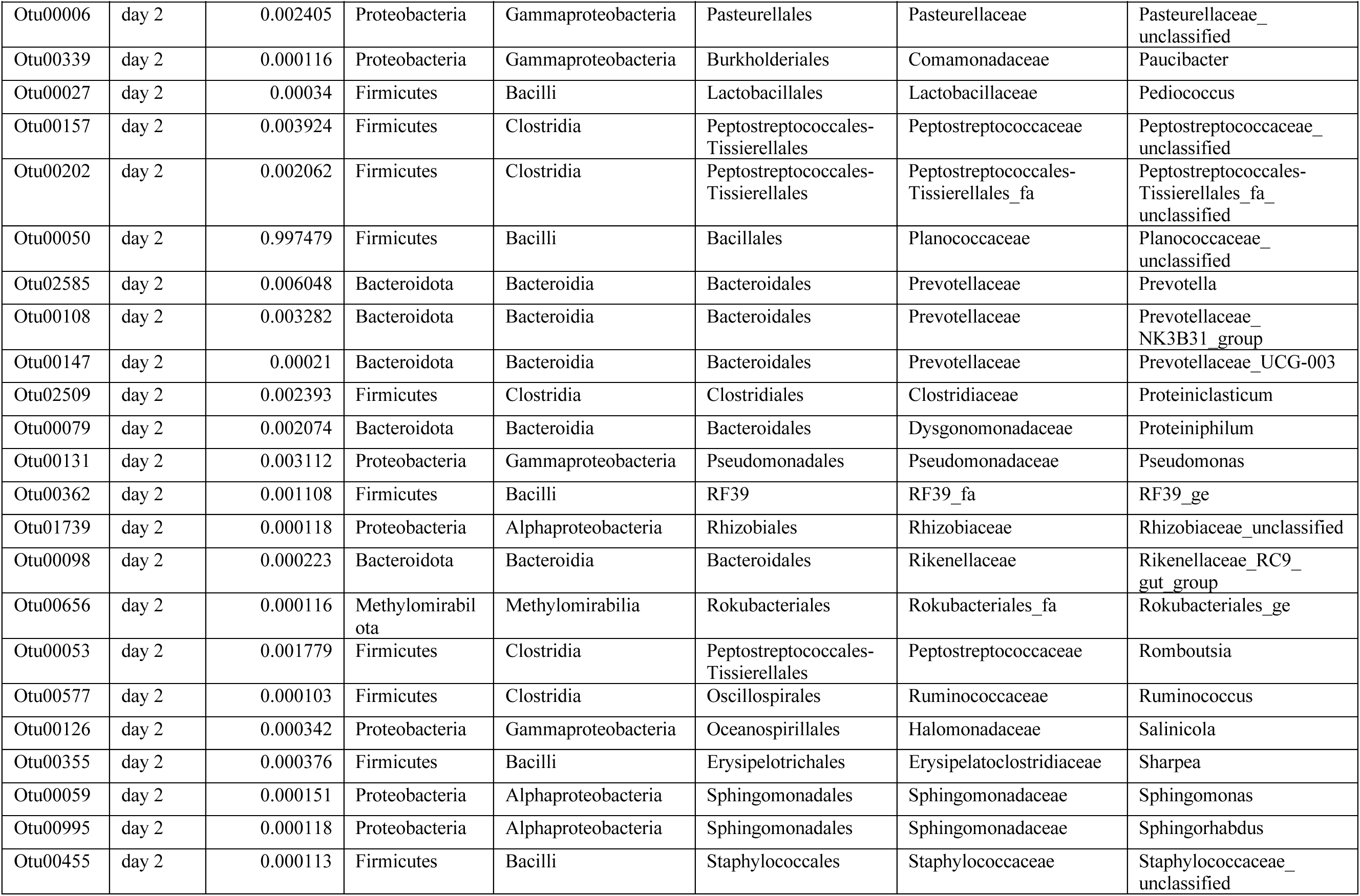
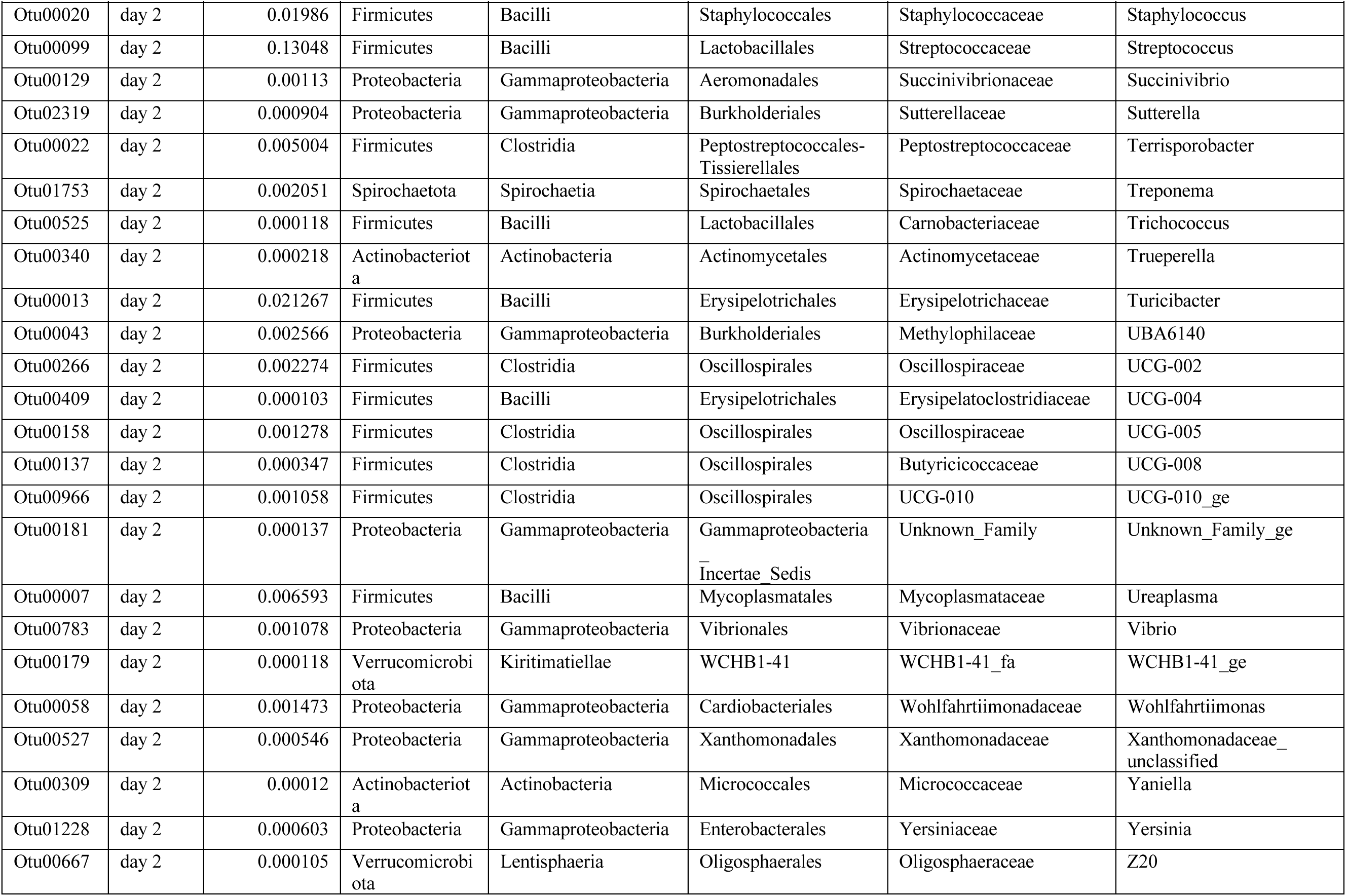
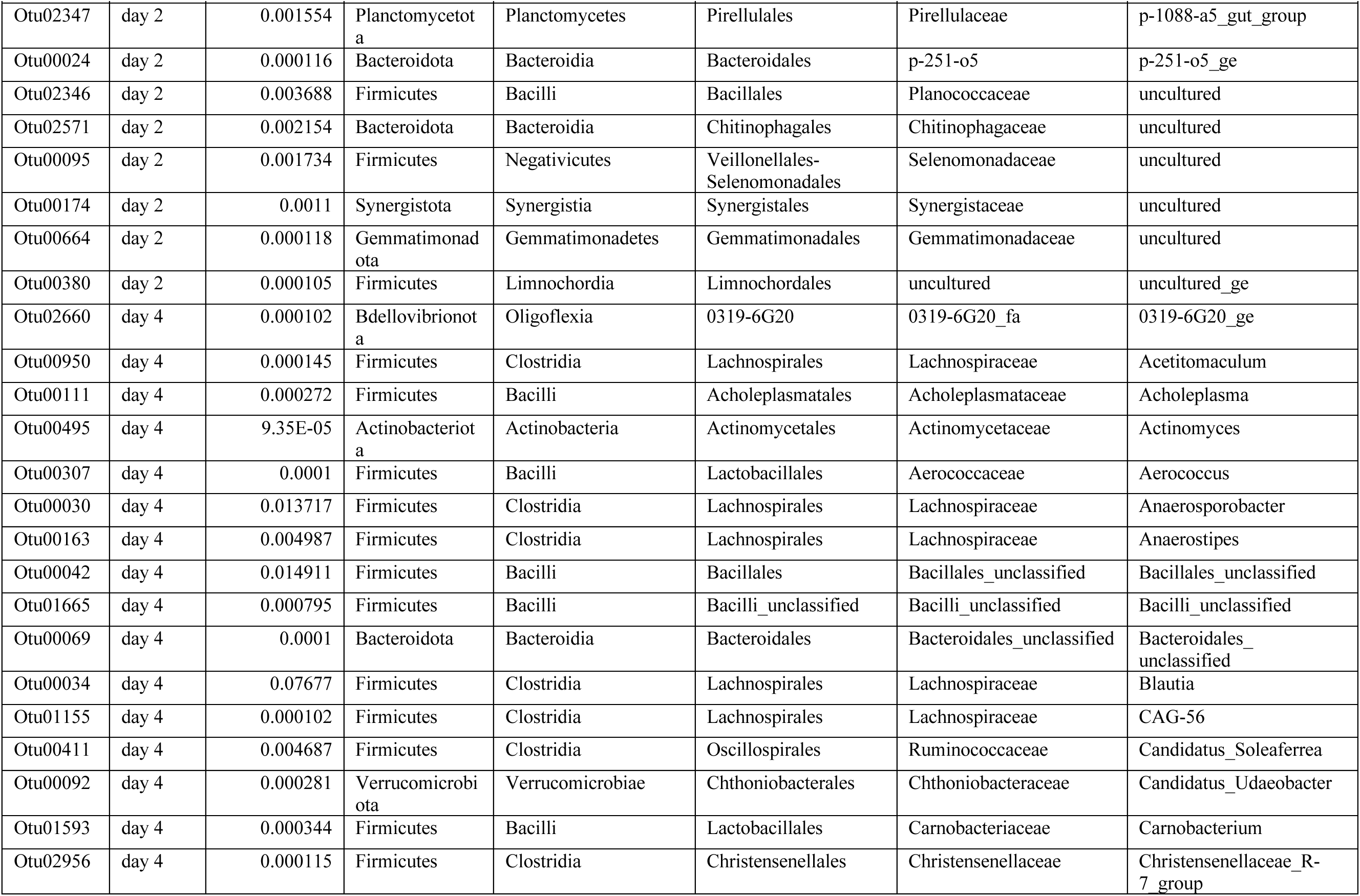
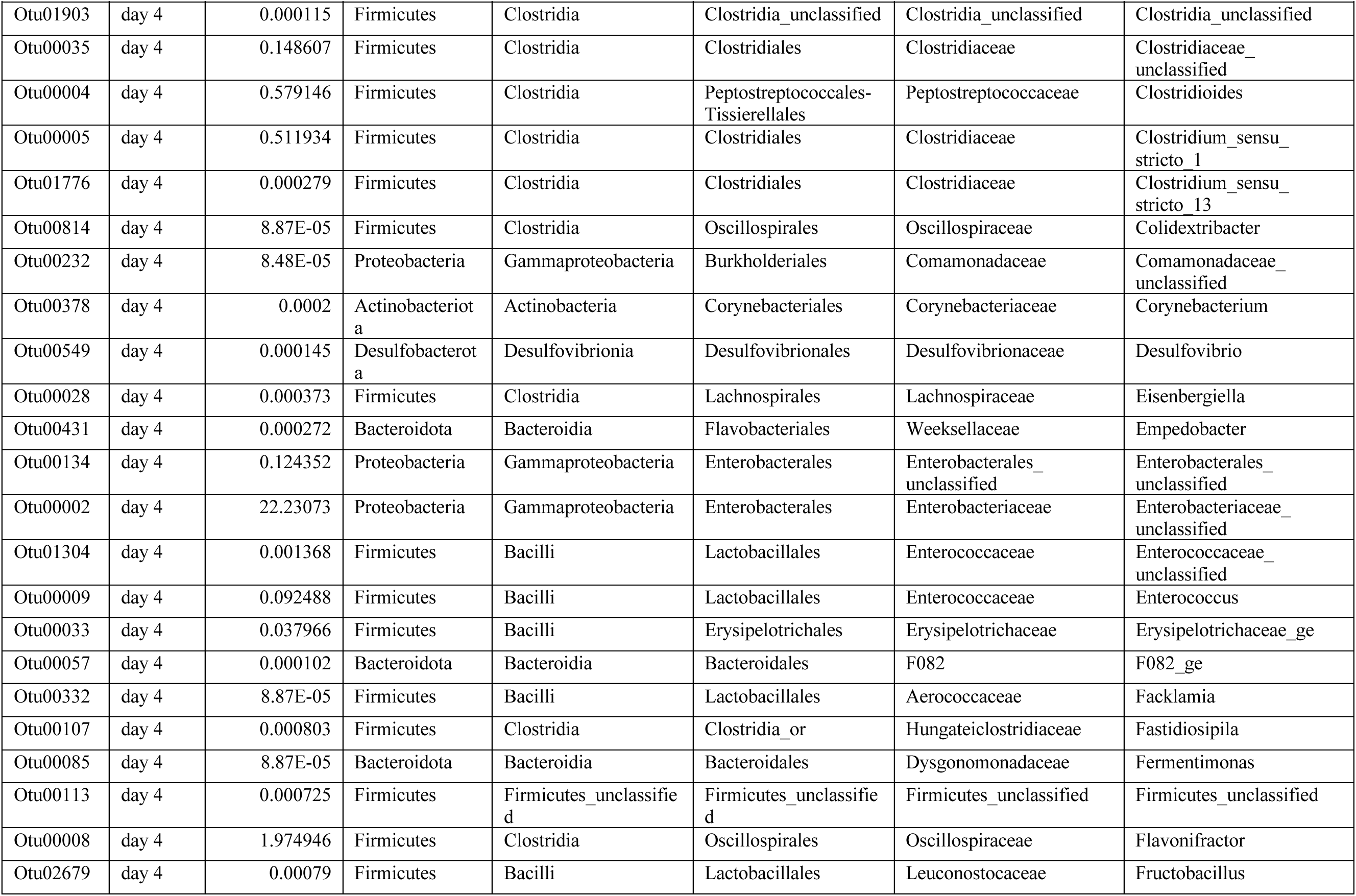
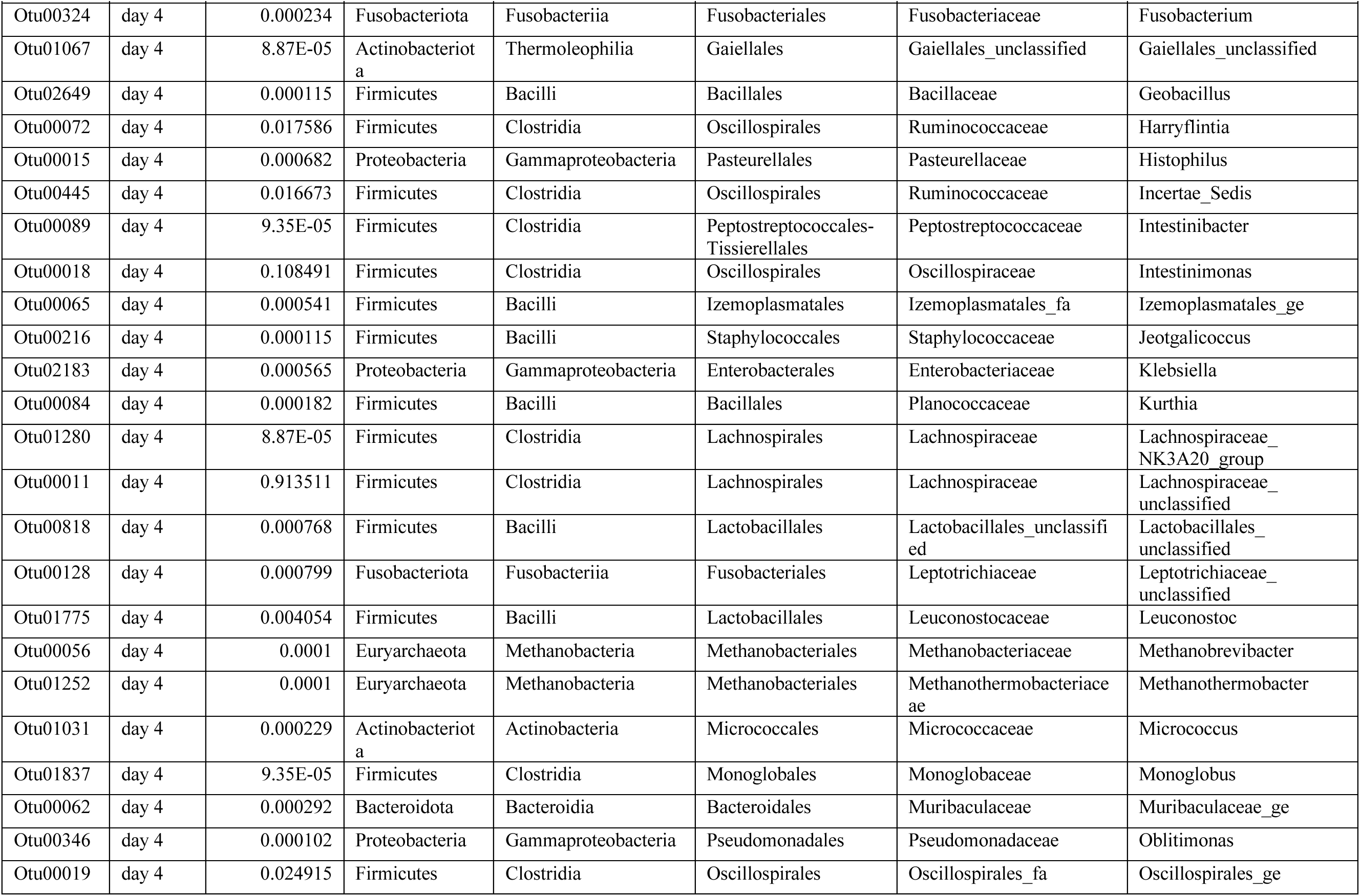
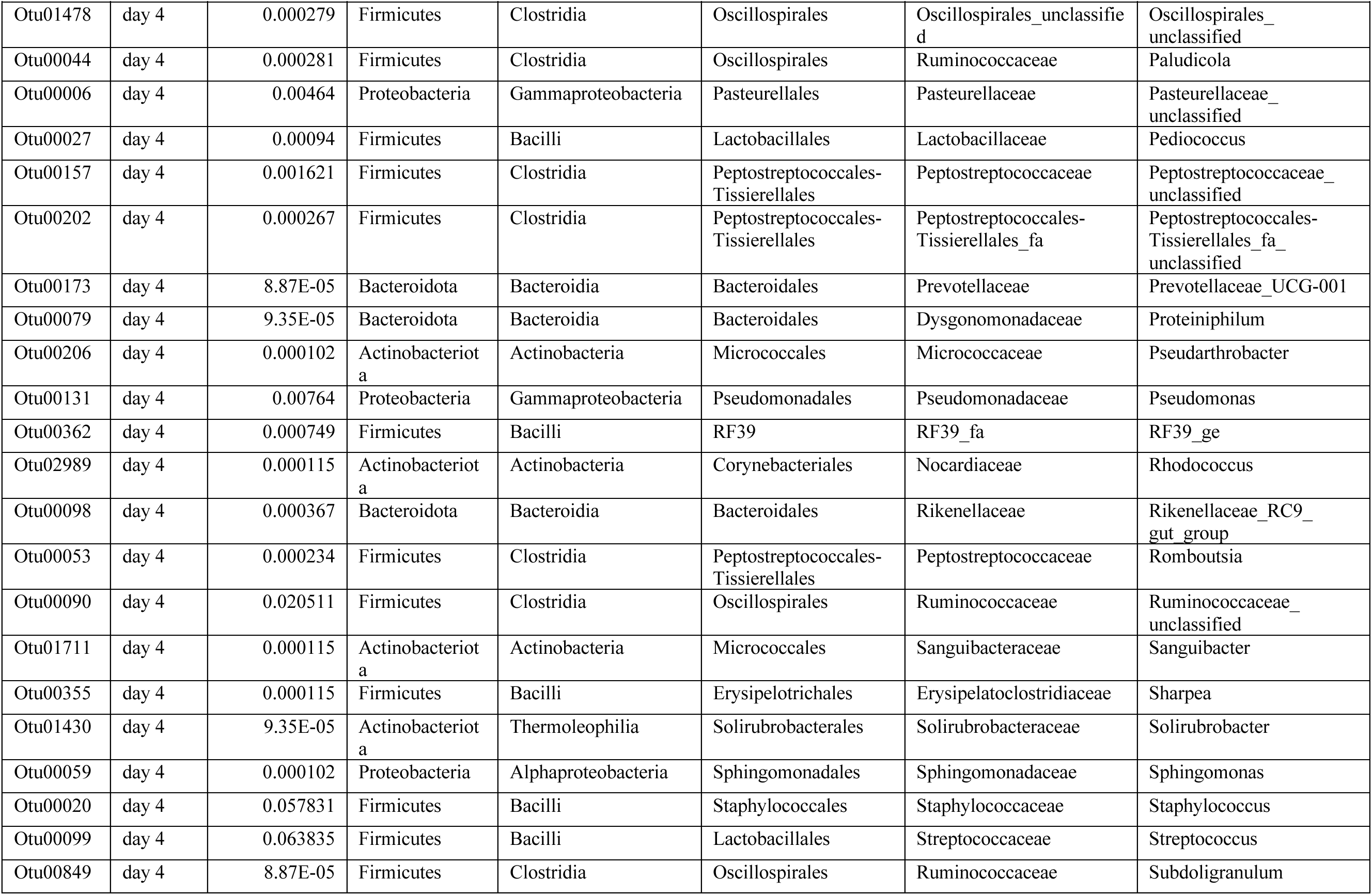
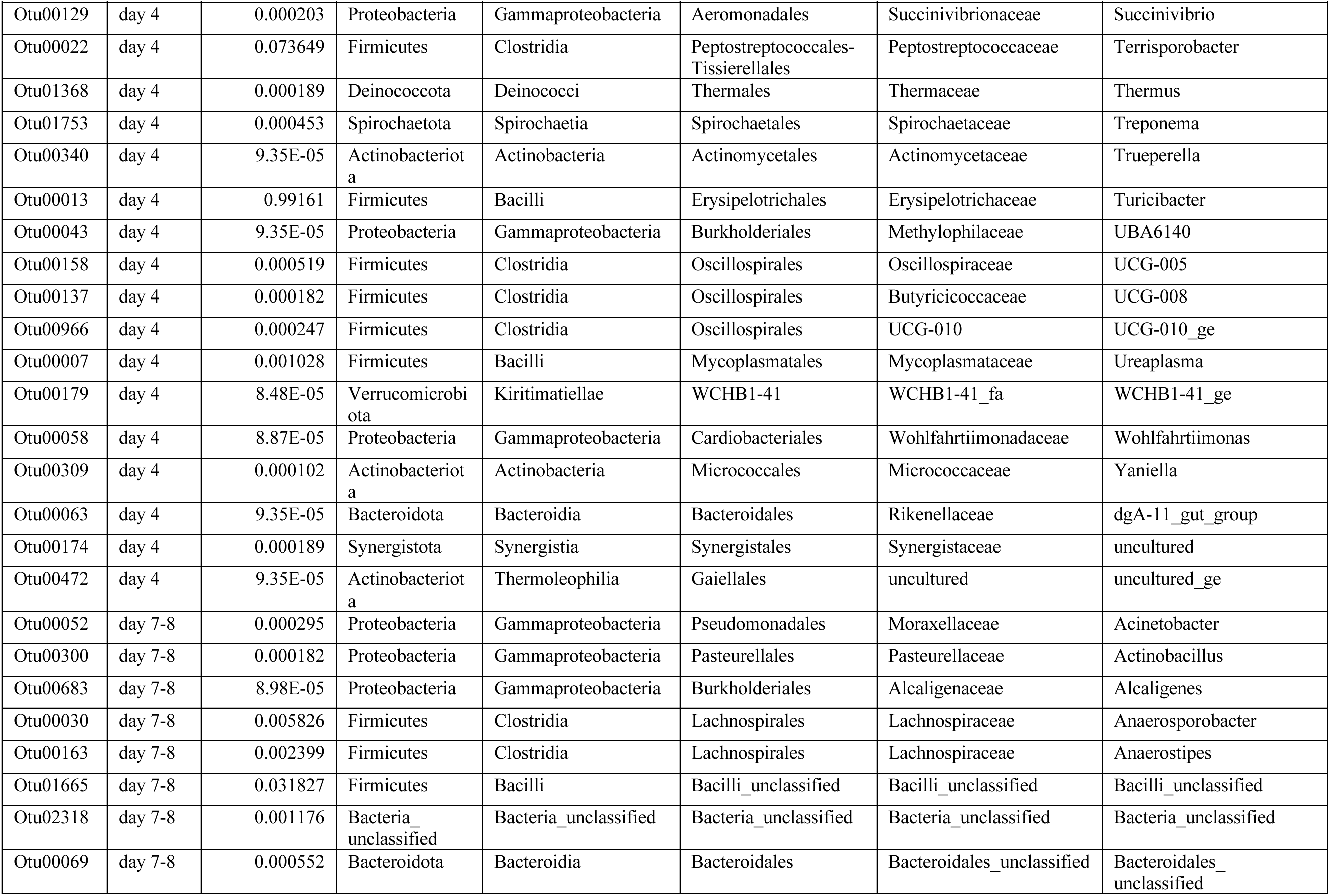
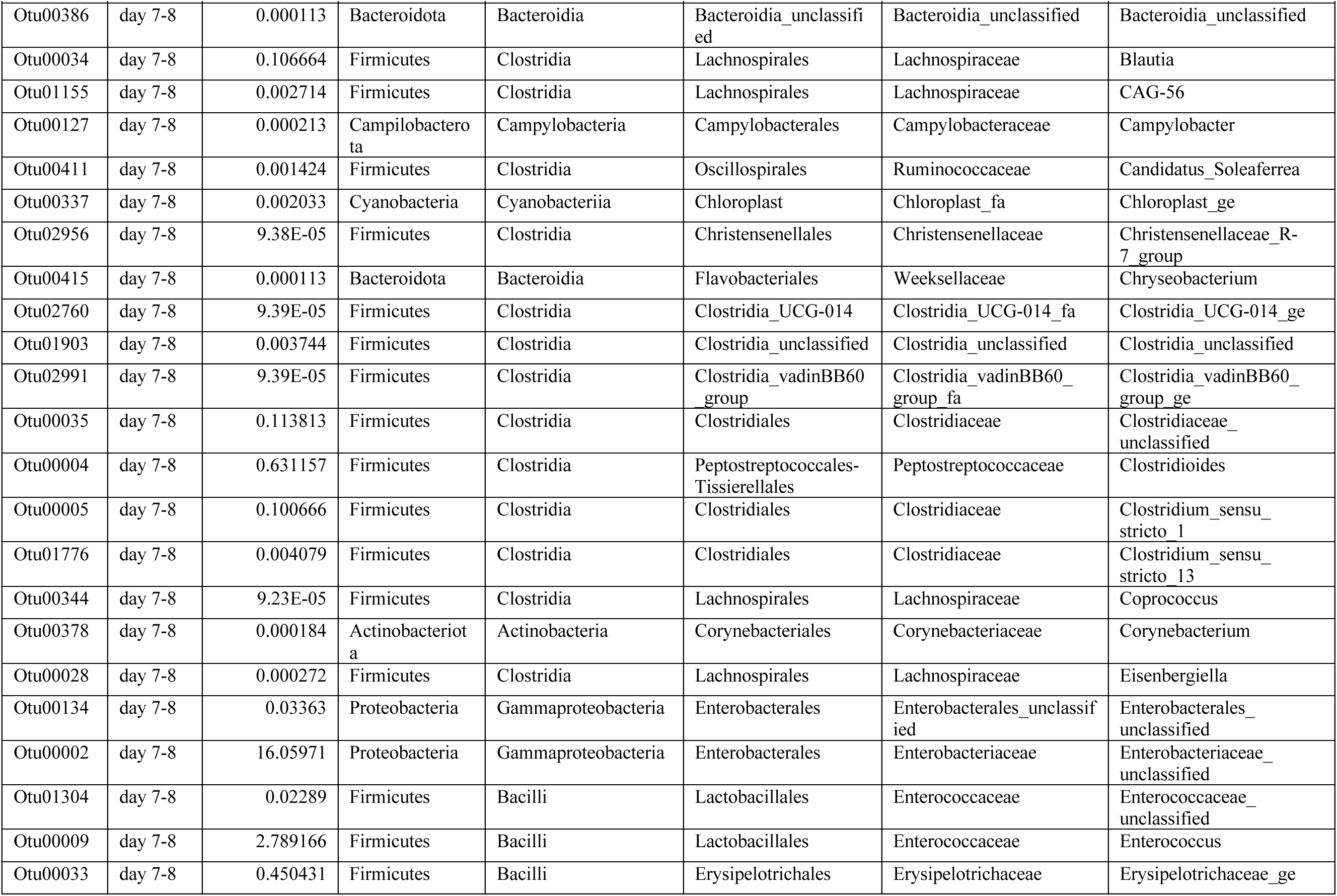
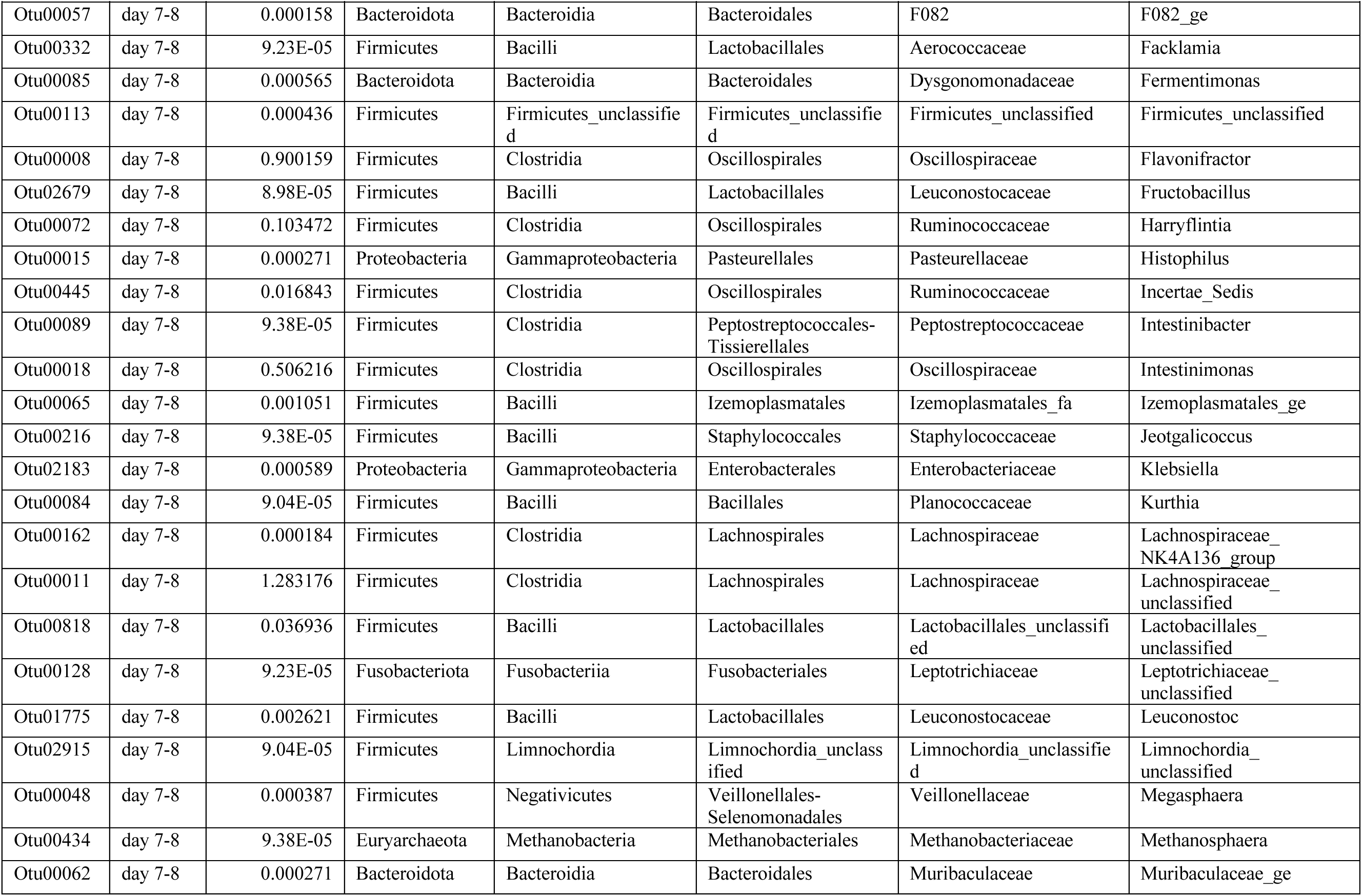
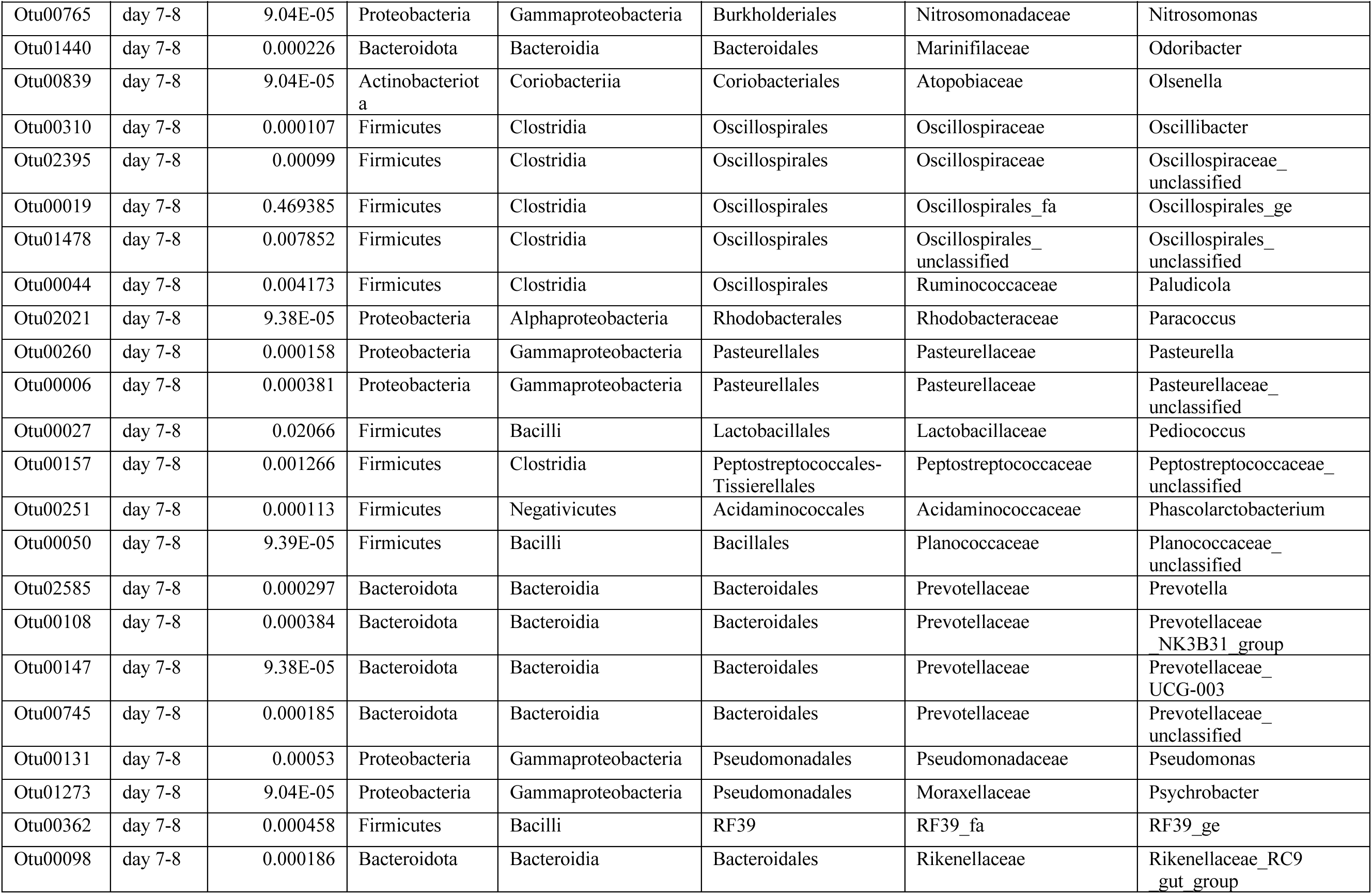
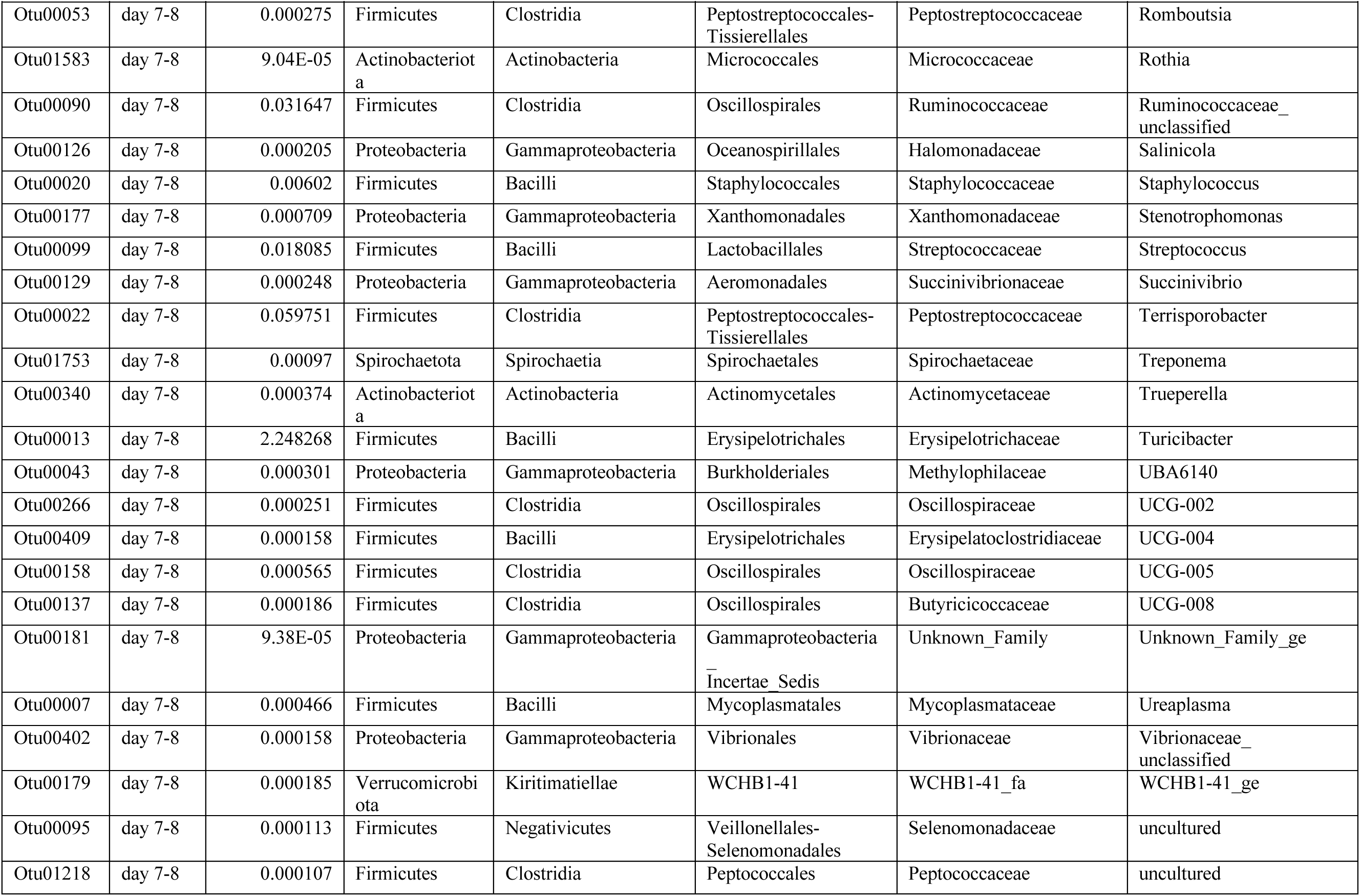
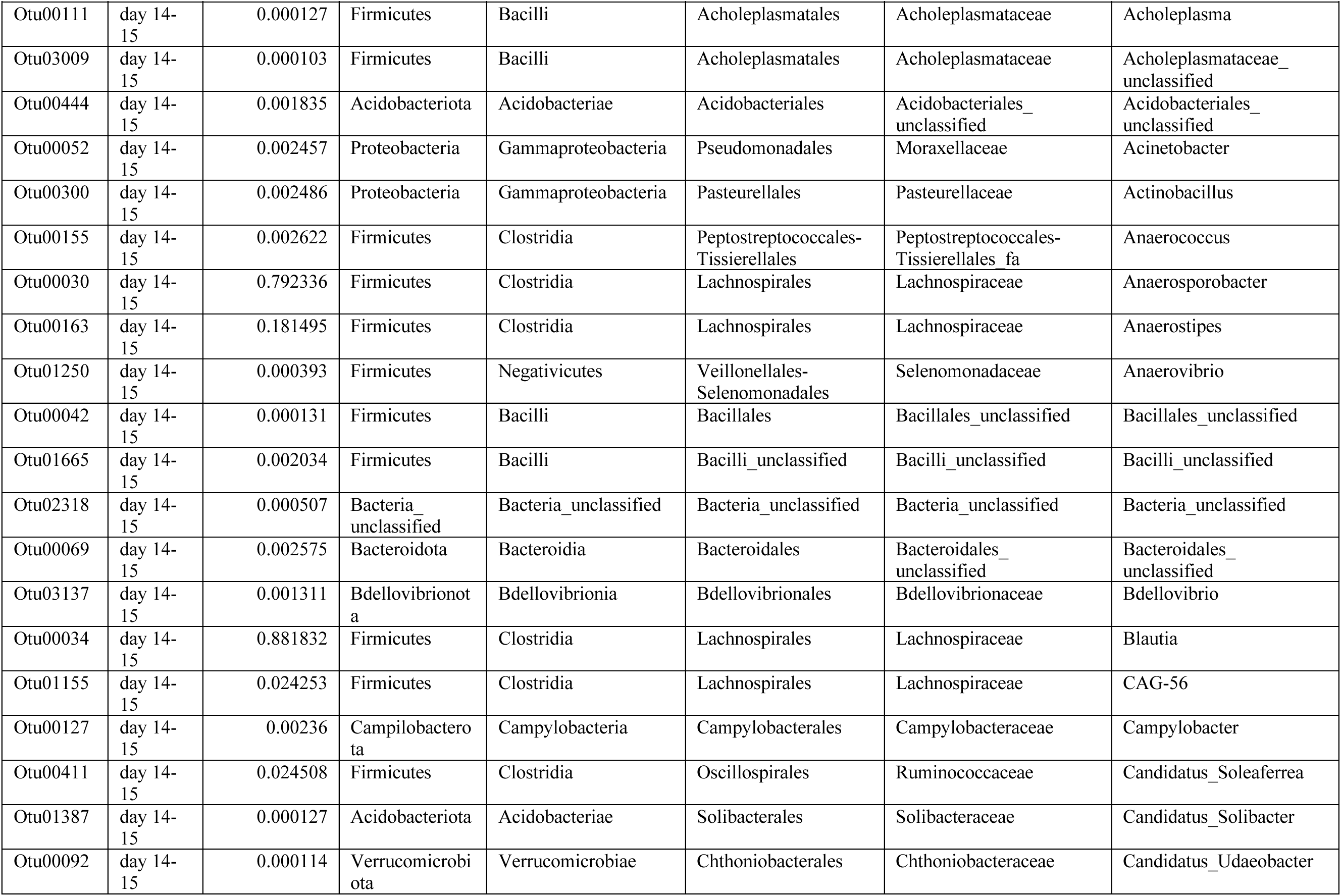
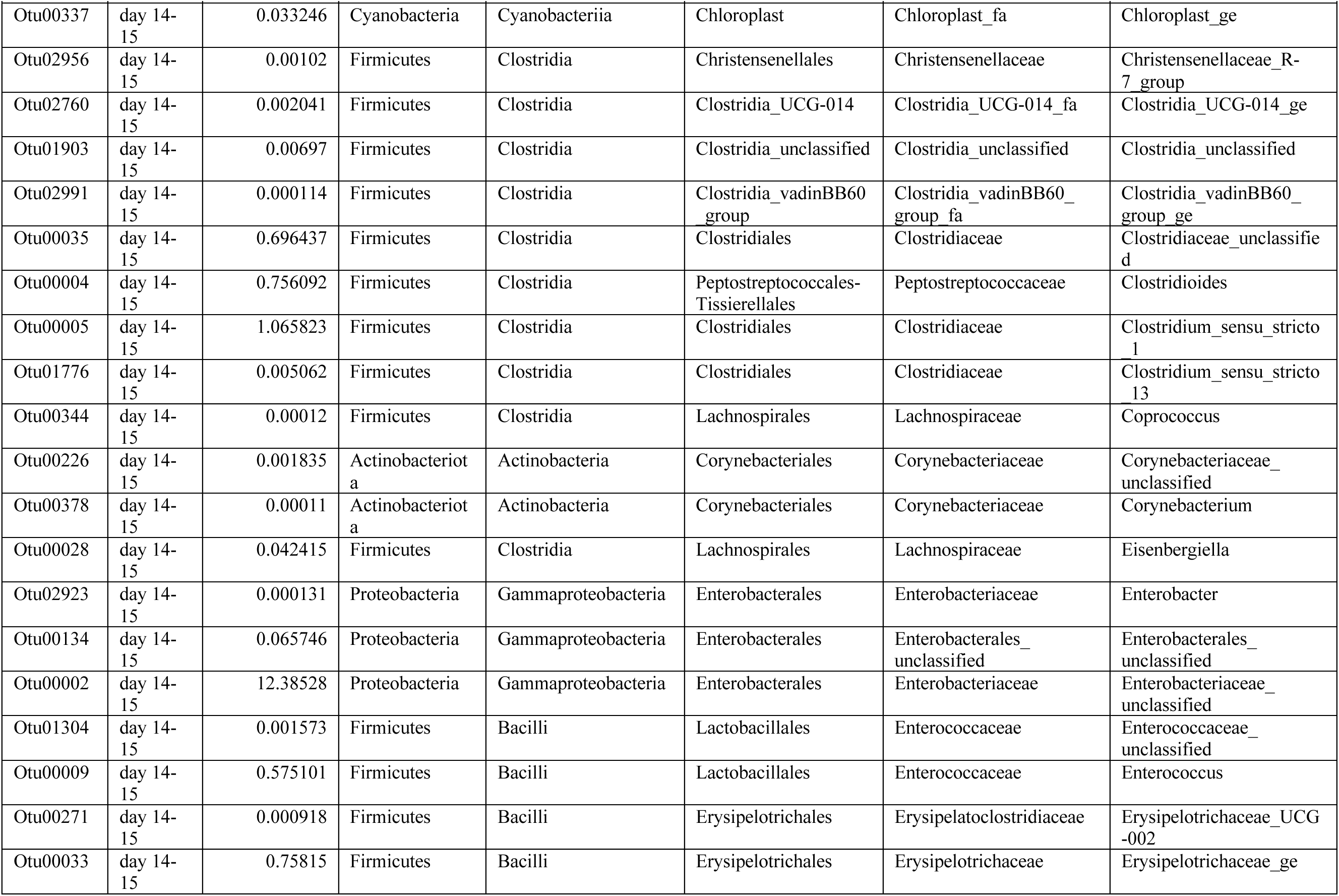
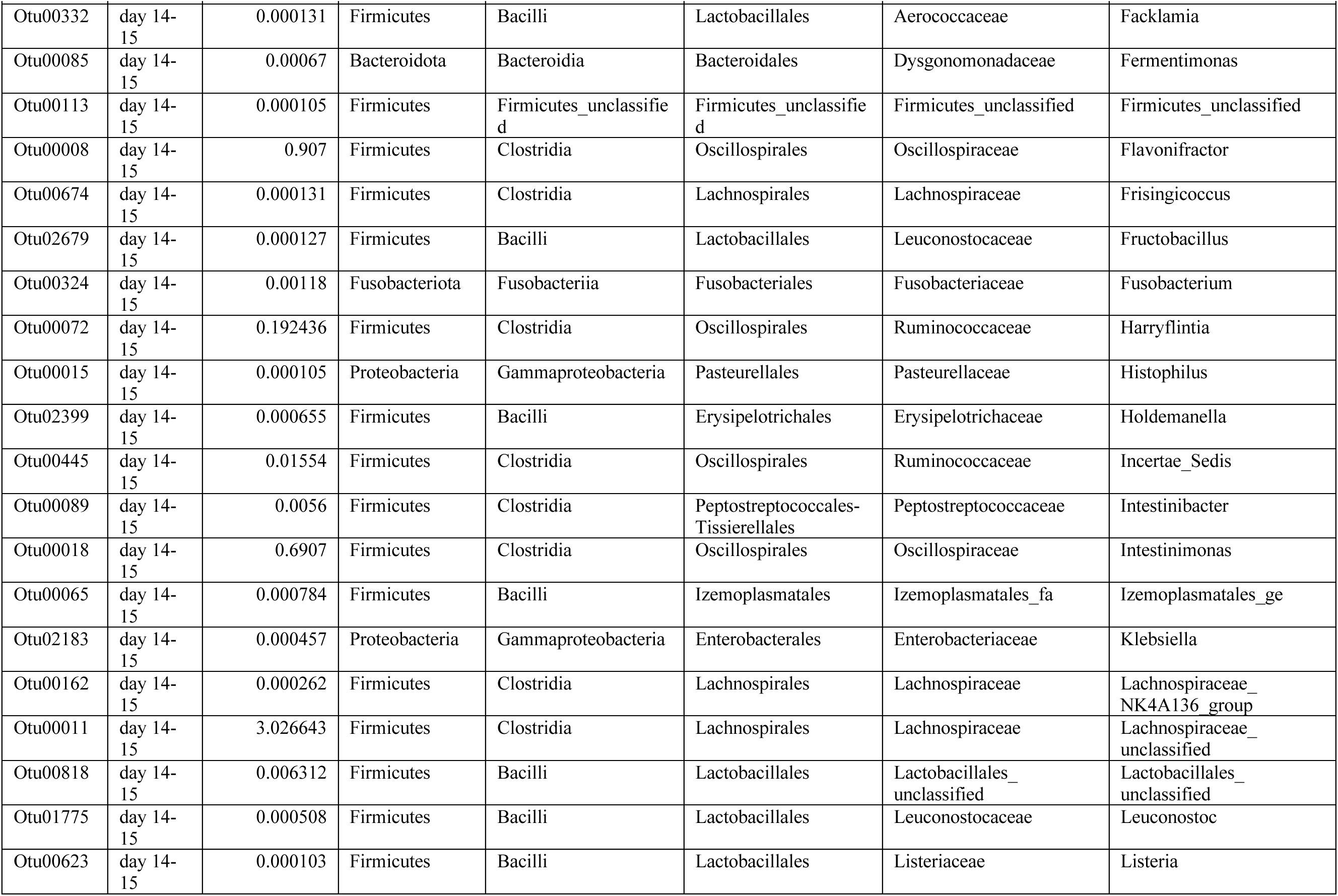
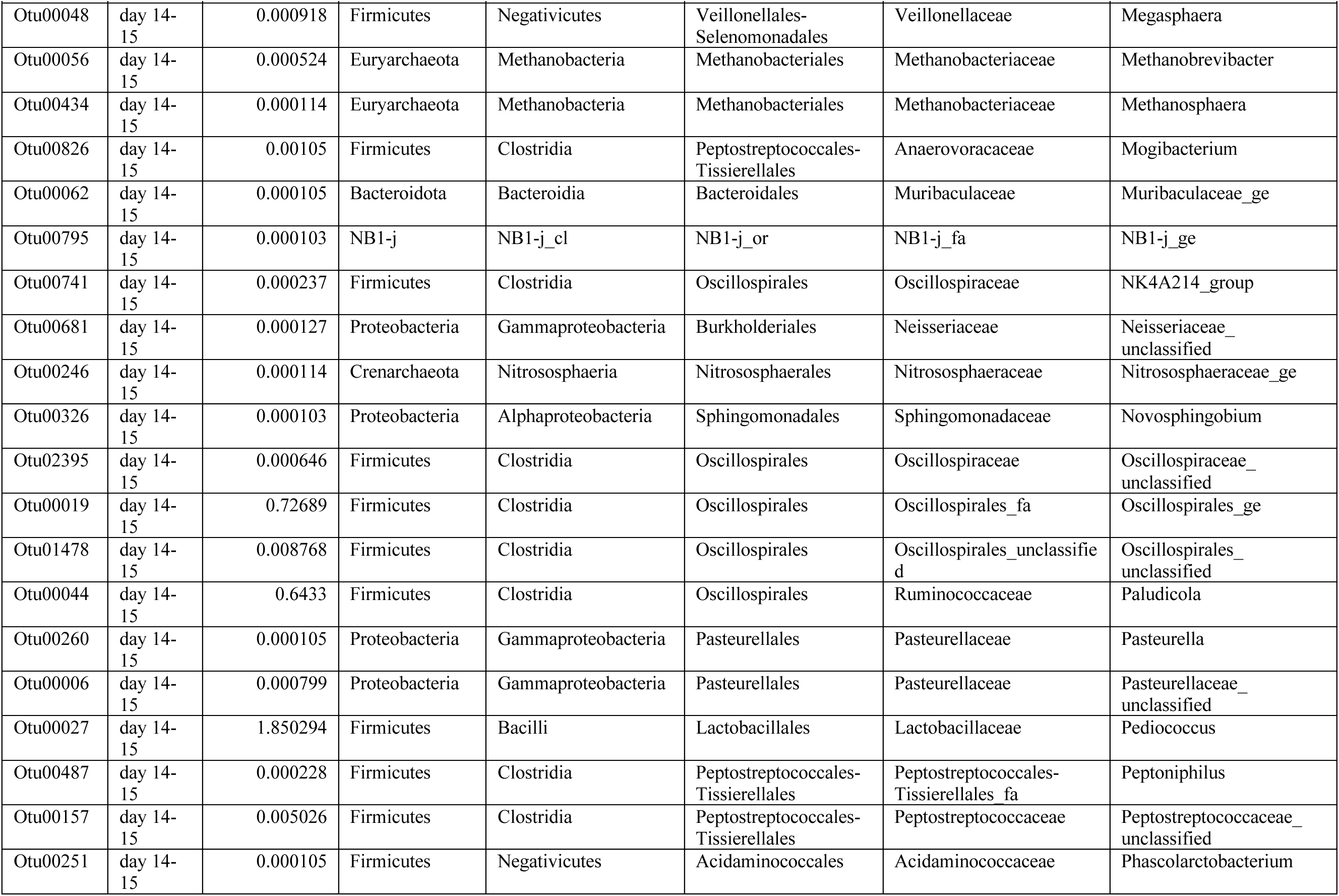
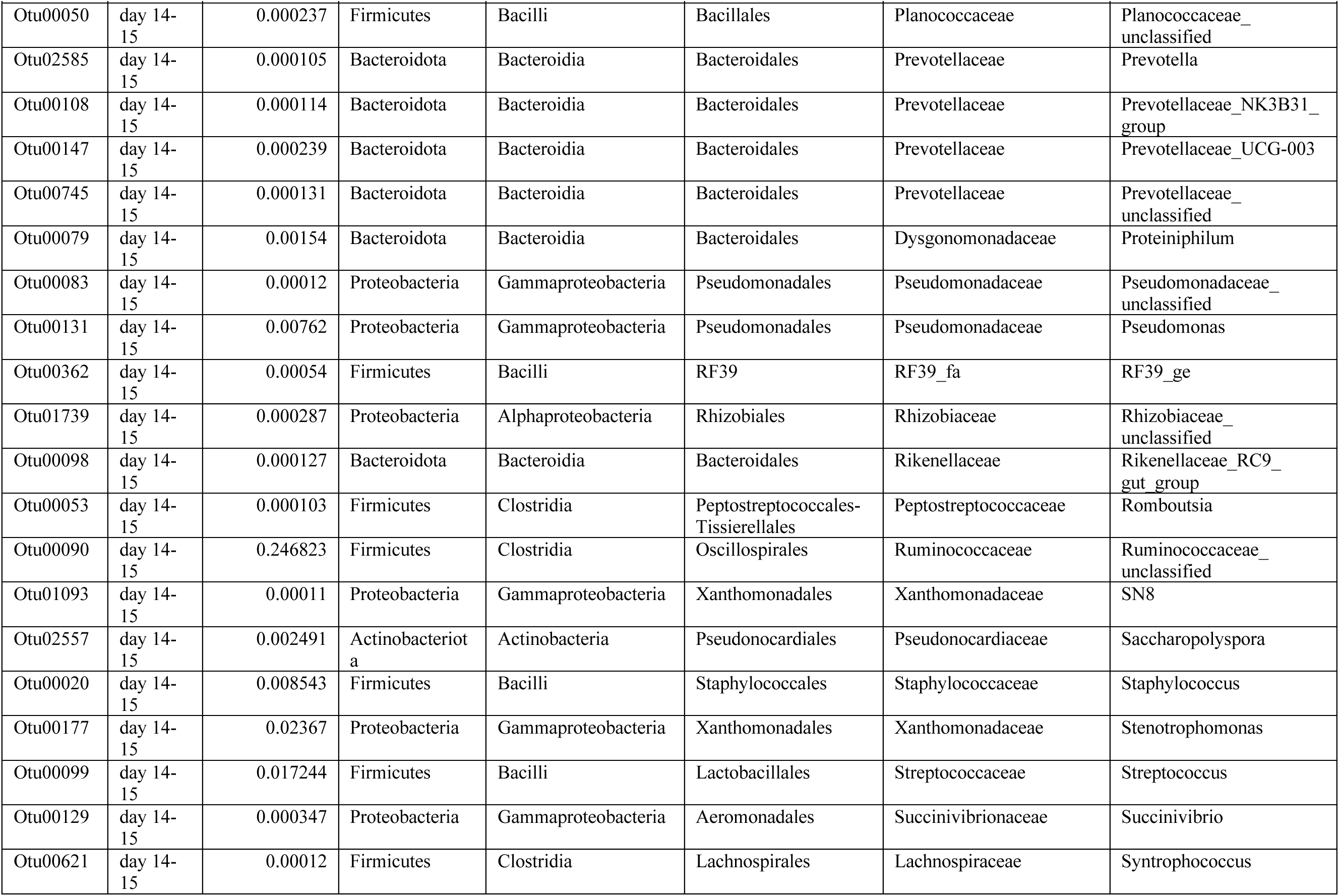
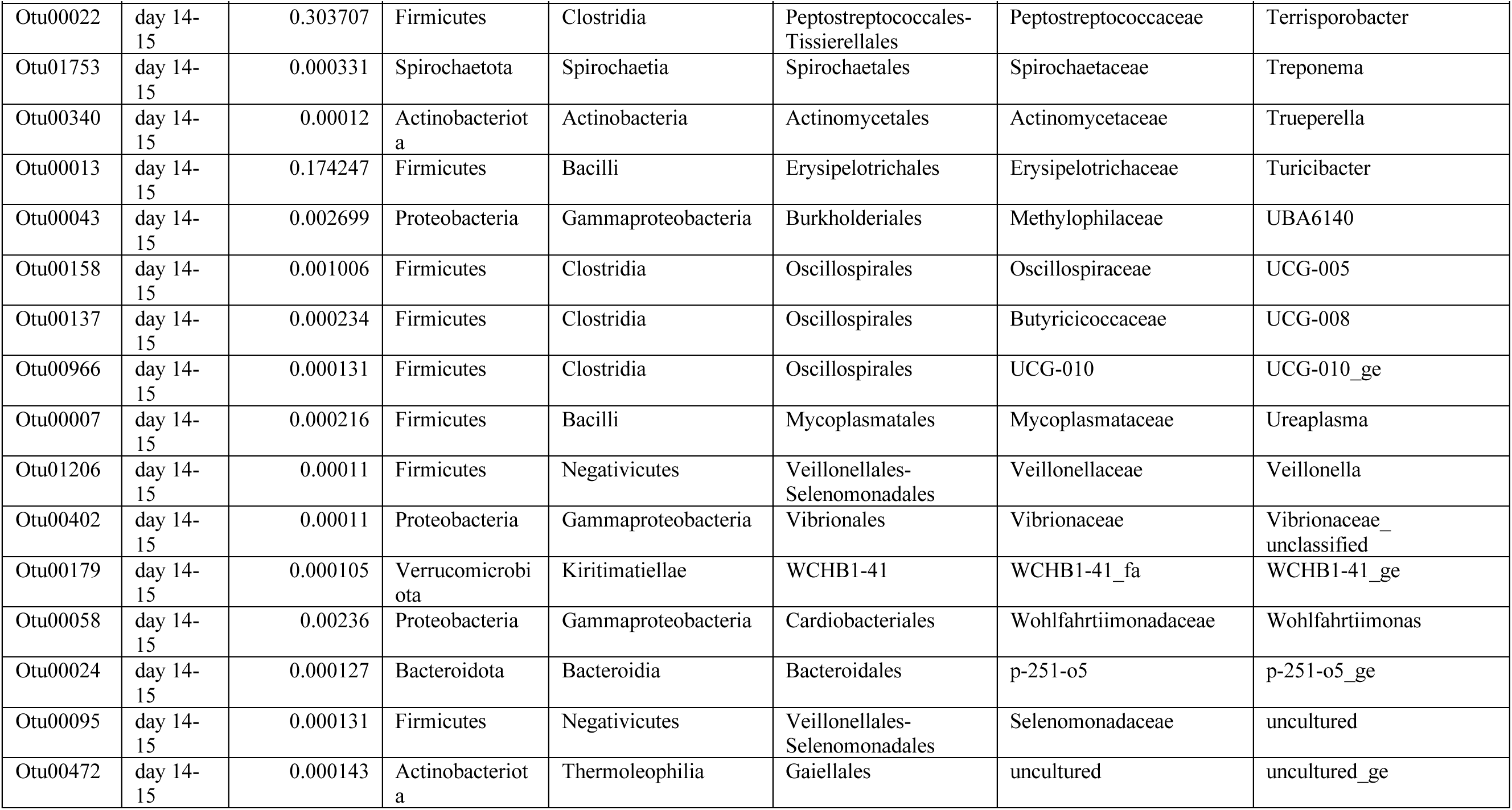
List of “Other” bacteria as seen in Figure 4 for litters 33 and 34.

## Notes

### Competing Interest Statement

The authors have declared no competing interest.

